# Mapping the Modification Landscape of MHC-I Epitopes: A Framework for Immunogenic Peptidomimetic Antigen Design

**DOI:** 10.64898/2026.03.06.710184

**Authors:** Sarah E. Newkirk, Joey J. Kelly, Nita Hourn, Sobika Bhandari, Naomi Spencer, Marcos M. Pires

## Abstract

Peptide-based cancer vaccines offer a promising strategy to target tumor-specific neoantigens. This approach is increasingly critical as post-translationally modified peptides, driven by altered tumor metabolism, emerge as a unique class of neoantigens. Because these chemically distinct epitopes cannot be genetically encoded by mRNA or viral platforms, synthetic peptide vaccines are poised to be the primary route to target these types of neoantigens. Yet, their clinical translation is restricted by poor metabolic stability, limited intracellular permeability, and structural requirements for MHC-I binding and T cell receptor (TCR) recognition. Although peptidomimetic modifications have been widely explored to improve pharmacokinetics, their impact on antigen presentation and immune recognition remains poorly understood. Here, we undertook a comprehensive evaluation of peptidomimetics geared at MHC-I neoantigens and generated a diverse library of systematically modified peptides that incorporate backbone *N*-methylation, peptoid substitution, and stereochemical inversion. Integrated assays revealed a highly position-dependent tolerance to peptidomimetic modifications, while subsequent combinatorial designs demonstrated non-additive effects on the balance between immunogenicity and pharmacokinetics. Collectively, these findings establish design principles and provide a framework for balancing immune recognition with enhanced stability and permeability in peptidomimetic antigen design.

## INTRODUCTION

The adaptive immune system continuously monitors cells for signs of infection, malignancy, or other aberrations through sophisticated molecular recognition mechanisms. Central to this surveillance is the presentation of intracellular peptides to patrolling cytotoxic T cells, enabling the immune system to distinguish healthy cells from those that are compromised. Major histocompatibility complex class I (MHC-I) molecules are membrane proteins expressed on nearly all nucleated cells, serving a role in immune surveillance.^1^ Their principal function is to present intracellularly derived peptides, typically 8-10 amino acids in length, on the cell surface bound to the MHC-I binding groove. These peptide-MHC complexes (pMHCs) are recognized by T cell receptors (TCRs) on CD8+ cytotoxic T lymphocytes (CTLs), triggering T cell activation and subsequent destruction of the infected or abnormal cells.^2^

In cancer, tumor-associated and tumor-specific peptides can arise from proteomic alterations such as somatic mutations,^3^ aberrant splicing,^4–7^ or post-translational modifications (PTMs),^8^ generating neoantigens. Notably, altered tumor metabolism frequently drives the formation of non-enzymatic PTMs, yielding chemically distinct epitopes that differ fundamentally from normal self-peptides. Because these unique structural modifications cannot be genetically encoded by mRNA or other genetic-based viral platforms, harnessing them to elicit potent anticancer immune responses strictly requires synthetic peptide-based vaccination.^9^ This immunological distinction has positioned neoantigens as valuable targets for personalized cancer immunotherapy. In parallel with efforts to identify immunogenic peptides, complementary strategies have emerged to modulate MHC-I antigen presentation itself, including the use of small molecules to regulate MHC-I surface expression.^10^

Efforts to modulate MHC-I presentation are further amplified by a growing understanding of the exact structures of the peptides being presented on cancer cells analyzed by immunopepidomic. Recent seminal profiling of the tumor MHC-I immunopeptidome has fundamentally reshaped this landscape. Utilizing an integrated antigen discovery pipeline, Kacen et al. ^8^ demonstrated that post-translational changes driven by the tumor proteome dramatically expand the targetable antigen landscape. Crucially, their work revealed that PTM-driven neoantigens can be shared not only between patients but across diverse cancer types. This paradigm-shifting discovery positions PTM-modified epitopes as highly exciting candidates for cancer vaccination, as it establishes the feasibility of scalable, "off-the-shelf" synthetic peptide vaccines that bypass the logistical bottlenecks of strictly individualized therapies.

To translate these candidates into the clinic, a promising strategy utilizes these synthetic peptides to target antigen-presenting cells (APCs), allowing them to naturally process and present MHC-I-restricted epitopes to prime tumor-specific T cells (**Fig. 1a**).^11–13^ These vaccines offer distinct advantages, including tumor specificity, higher safety, ease of synthesis, chemical stability, and compatibility with adjuvants or delivery systems.^14^ Their feasibility has been demonstrated in clinical trials for solid tumors, particularly melanoma, often in combination with immune checkpoint inhibitors or dendritic cell-based vaccination strategies.^15–20^

**Figure 1.**
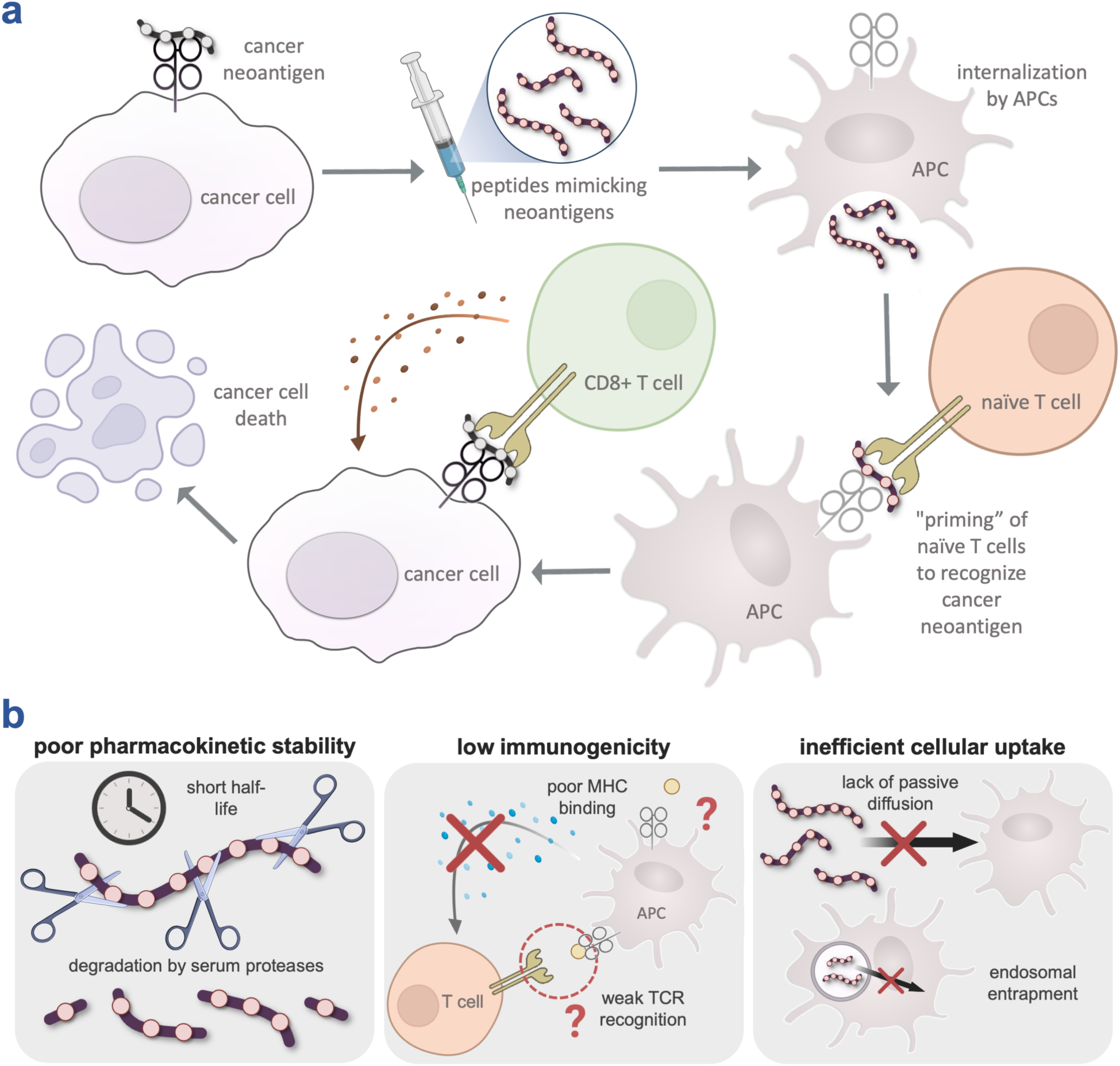
**(a)** Schematic illustrating the general mechanism of peptide vaccines for cancer. Cancer vaccines activate the host immune system by delivering synthetic peptides to antigen-presenting cells (APCs), allowing them to naturally process and present MHC-I-restricted epitopes to prime tumor-specific T cells. **(b)** Schematic depicting the three central challenges of synthetic peptide vaccines: poor pharmacokinetic stability, low immunogenicity, and inefficient cellular uptake.

We herein broadly and systematically investigate the impact of various peptidomimetic modifications on the fundamental mechanics of antigen presentation, covering MHC-I binding, TCR recognition, cellular permeability, and serum stability. More specifically, we generated and characterized a panel of modified analogs of SIINFEKL, a canonical MHC-I epitope derived from ovalbumin. Through integrated biochemical and cellular assays, we determine how *N*-methylation, peptoid, and ᴅ-amino acid substitutions modulate peptide behavior across these parameters. Our findings provide a framework for optimizing design of peptidomimetic vaccines with retained immunological properties.

## RESULTS AND DISCUSSION

Synthetic peptide vaccines face three central challenges: poor pharmacokinetic stability, low immunogenicity, and inefficient cellular uptake (**Fig. 1b**). First, rapid recognition and cleavage by serum proteases compromise pharmacokinetics and can diminish sustained antigen exposure.^21^ A second major limitation of peptide vaccines is their inherently low immunogenicity. Effective immune recognition requires two stringent molecular interactions: (1) the peptide must bind to MHC-I with sufficient affinity to form a stable complex, and (2) the resulting complex must be recognized by a specific TCR.^1, 2^ Even minor alterations to key anchor residues within the peptide can drastically reduce MHC binding, while changes at solvent-exposed positions can abolish TCR recognition.^22, 23^

Consequently, the limited antigenicity of short peptides typically necessitates co-administration with immunostimulatory adjuvants or carrier systems to elicit robust immune responses.^24^ A third, often underappreciated challenge is limited cellular uptake, as variations in peptide permeability may influence antigen fate following immunization. Due to their size and polarity, exogenous peptides passively diffuse across lipid bilayers inefficiently, instead remaining extracellular and vulnerable to degradation.^25–27^ As a result, only a small fraction reaches the cytosol for loading onto MHC-I molecules.^28^ This poor membrane permeability renders them generally unsuitable for oral administration, as they cannot traverse the intestinal epithelium to enter systemic circulation effectively. Furthermore, even when internalized by antigen-presenting cells (APCs) via endocytosis, peptides often remain trapped in endosomes, struggling to escape into the cytosol where the proteasomal processing required for MHC-I cross-presentation occurs.^29^ Permeability may also exert an indirect effect via uptake by APCs of antigen-loaded cell debris. Permeable peptides can enter non-APCs and become intracellular antigens. During immunization, inflammation at injection sites recruits innate immune cells and promotes necrosis. When those non-APC “donor” cells die, their contents (including internalized peptides) are released and captured by APCs through cross-priming pathways.^30, 31^ Specialized dendritic cell subsets are particularly adept at phagocytosing dying cells and diverting internalized antigens into the MHC-I processing pathway.^32, 33^ Thus, enhanced peptide permeability may expand the available pool of antigenic material through both direct and indirect mechanisms.

Multiple strategies have been considered to address bottlenecks to peptide vaccines, primarily focusing on structural modifications to enhance peptide stability and immunogenicity.^34^ These approaches, broadly, aim to enhance protease resistance and bioavailability while preserving the physicochemical properties required for antigen presentation.^13^ To overcome these challenges, multiple stabilization strategies have been explored.^35^ These include cyclization,^36–38^ PEGylation,^39, 40^ side chain substitutions (e.g., ʟ- to ᴅ-amino acid inversion,^41–43^ retro-inversion^44–48^), backbone modifications (e.g., α- to β-amino acid replacement^49–53^, thioamide substitution,^54^ *N*-methylation,^55, 56^ reduction^57, 58^), combined backbone and side chain alterations (*N*-alkyl glycines or peptoids)^59–61^, and modifications at the termini (*N*-terminal methylation and *C*-terminal amidation).^62^ Examples of such modifications have been applied to clinically relevant epitopes such as MAGE-1.A1,^63^ Melan-A/MART-1,^50, 64, 65^ and UTA2-1,^66^ improving antigenicity and proteolytic stability. While increasing peptide stability is a standard strategy for enhancing vaccine efficacy, optimizing cellular uptake has received less attention, despite its critical role. The relationship between structural modifications and peptide accumulation in mammalian cells is well understood; however, these principles have not yet been evaluated in the context of MHC presentation. Specifically, *N*-methylation and stereoinversion are strategies commonly employed to optimize peptide stability and permeability.^67–69^ Since prior studies,^70^ including our own work,^71^ demonstrate that these modifications can drive peptide accumulation, they represent a promising, underexplored avenue for enhancing vaccine delivery.

### Design of Peptidomimetic Libraries and Effect of Peptidomimetic Substitutions on pMHC-I Stability

To establish a consistent framework for our libraries, we selected the base model epitope SIINFEKL (**ovaWT**, **Fig. 2a**) derived from the protein ovalbumin (OVA). SIINFEKL is widely employed in antigen presentation studies due to its well-characterized interactions with the murine MHC-I molecule H-2K^b^ and SIINFEKL-specific TCRs, providing a robust system for comprehensively analyzing structural modifications.^72^ To assess the influence of peptidomimetic modifications on pMHC-I stability, we used RMA-S cells, a cell line lacking functional transporters associated with antigen processing (TAP).^73, 74^ TAP transports cytosolic peptides into the endoplasmic reticulum (ER) for loading onto MHC-I molecules. In the absence of TAP, endogenous antigen presentation is compromised, and unloaded MHC-I molecules become unstable, leading to significantly reduced surface expression of the H-2K^b^ haplotype (**Fig. 2b**). However, lowering the incubation temperature promotes the transient surface expression of these empty MHC-I molecules. Subsequent incubation with high-affinity extracellular peptides stabilizes the H-2K^b^ complexes, allowing them to remain on the cell surface even after returning to physiological temperature (37°C). Previous studies have demonstrated a strong correlation between the abundance of surface pMHC-I complexes and peptide affinity for H-2K^b^.^75–78^ Thus, this system provides a controlled cell-based platform to isolate the effects of peptidomimetic modifications on pMHC-I binding.

**Figure 2.**
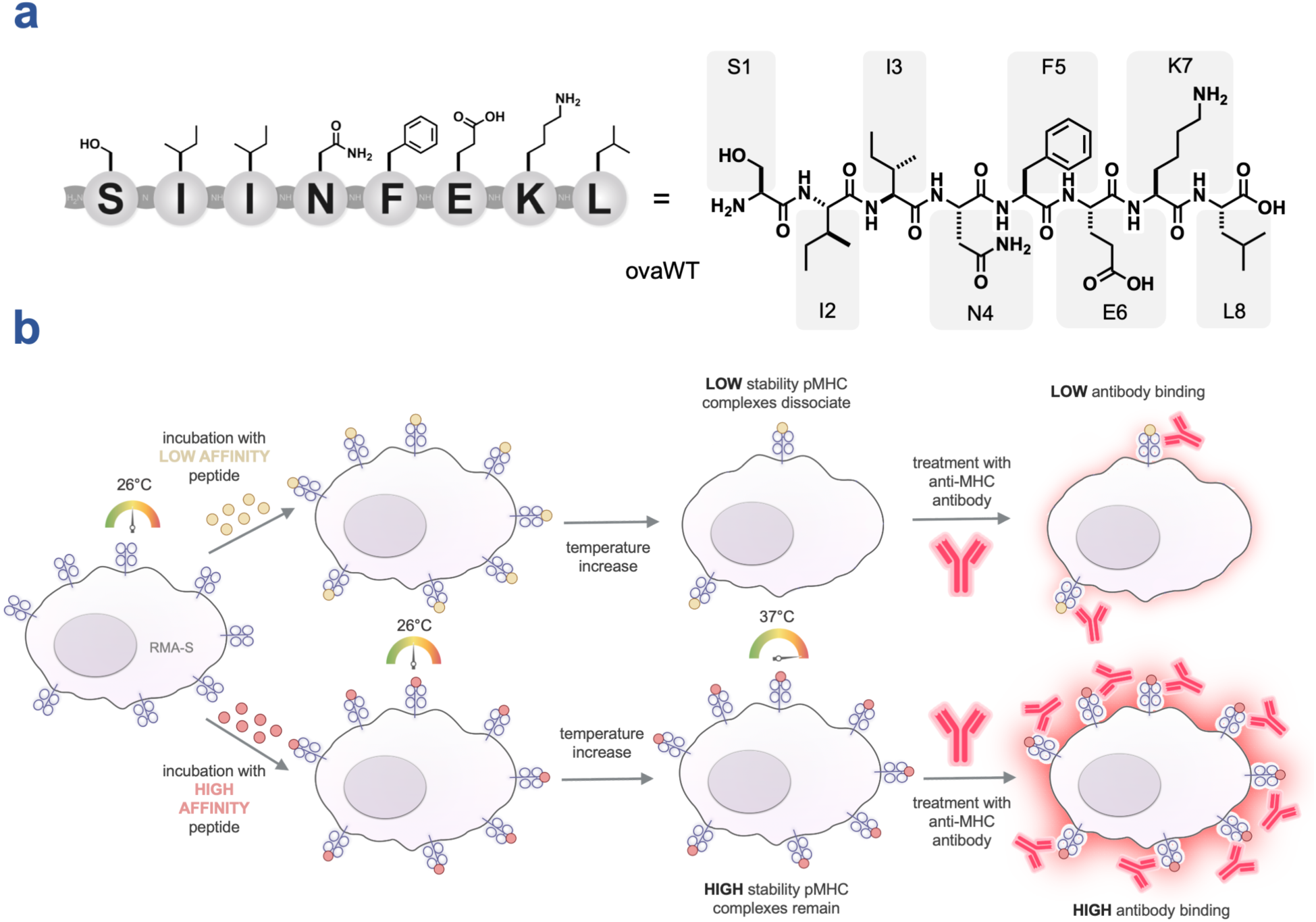
**(a)** Chemical structure of **ovaWT**. **(b)** Schematic illustrating the RMA-S stabilization assay.

Building on the base **ovaWT** peptide, we first generated a library of *N*-methylated peptides (**Fig. 3a**), in which each variant contained a single backbone methylation systematically introduced across the entire **ovaWT** sequence (**ovaNmet1–8**) (**Fig. 3b**). As described briefly above, *N*-methylation can sterically hinder proteolytic access to the backbone and eliminate the amide hydrogen bond donor, thereby reducing the energetic penalty associated with desolvation during membrane traversal. Although these effects can synergistically enhance metabolic stability and passive permeability, they may also fundamentally reshape the peptide’s conformational landscape. The introduced methyl group at the backbone nitrogen restricts rotation about adjacent torsion angles and increases the propensity for *cis*-amide isomer formation. This modification effectively rigidifies the peptide backbone and biases conformational sampling toward more pre-organized states. Given these structural and biophysical consequences, it is critical to empirically and systematically evaluate the impact of this modification in the context of MHC biology.

**Figure 3.**
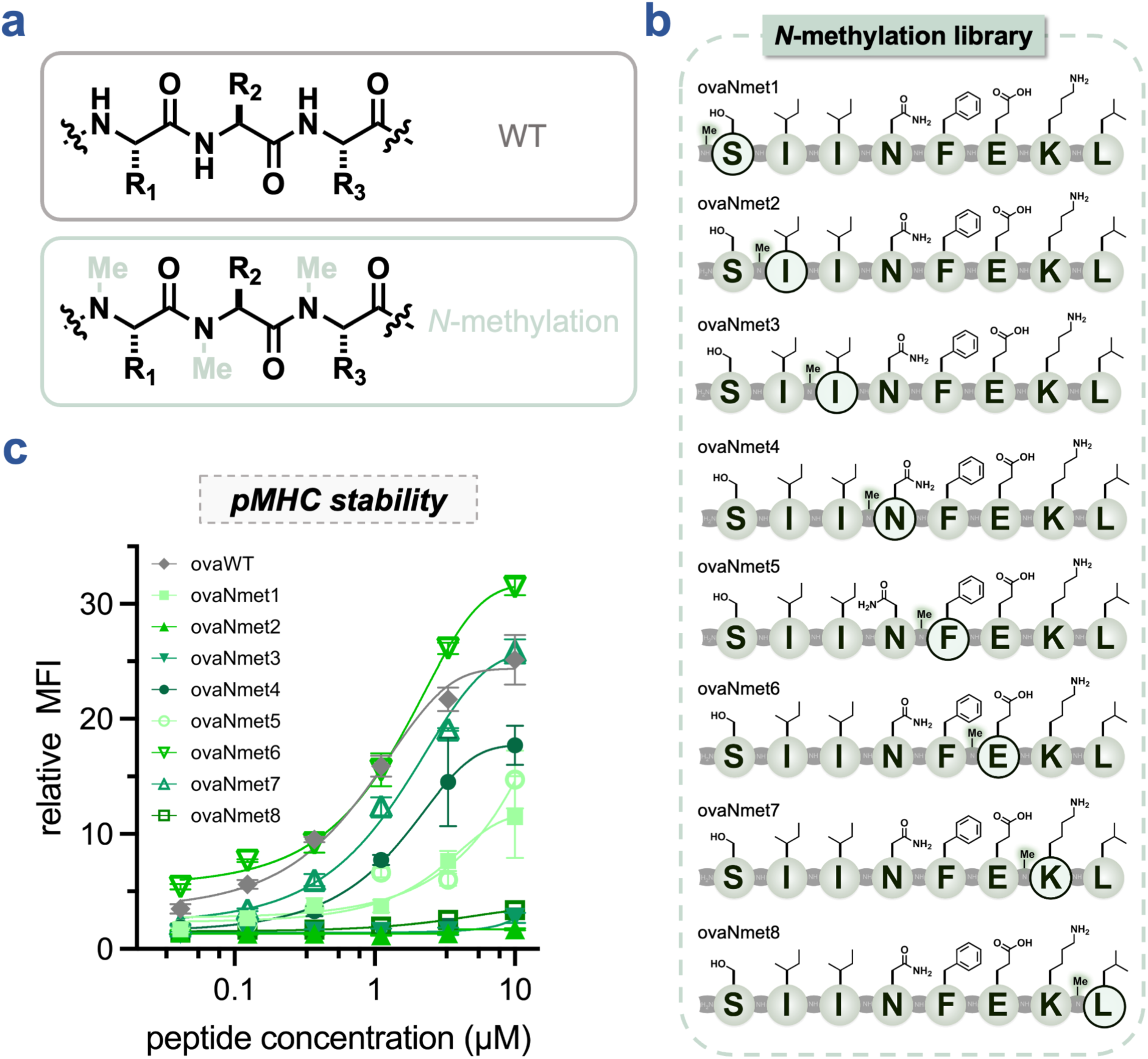
**(a)** Schematic depicting generic tripeptide with *N-*methylation modifications. **(b)** Chemical structures of the singly substituted *N*-methylation library. **(c)** Dose-response curves from flow cytometry analysis of the RMA-S stabilization assay. RMA-S cells were incubated with the indicated concentration of **ovaWT** and peptidomimetics from singly-substituted *N*-methylation library. H-2K^b^ expression was analyzed via flow cytometry by APC anti-mouse H-2K^b^ antibody. MFI is the mean fluorescence intensity of the level of fluorescence relative to the DMSO control. Data are represented as mean ± SD (*n* = 3), and Boltzmann sigmoidal curves were fitted to the data using GraphPad Prism.

Our data revealed that several of the *N*-methylated variants retained measurable ability to stabilize the pMHC complex (**Fig. 3c**). In particular, **ovaNmet6 and ovaNmet7**, featuring backbone *N*-methylation at the glutamic acid and lysine residues, respectively, exhibited pMHC stability comparable to the unmodified **ovaWT**. In contrast, three of the *N*-methylated derivatives completely abolished MHC binding: both isoleucine variants (**ovaNmet2** and **ovaNmet3**) and the leucine variant (**ovaNmet8**). These findings are consistent with the established characteristics of the **ovaWT**-H-2K^b^ complex. According to its crystal structure, three of the eight side chains (isoleucine at position 2, phenylalanine at position 5, and leucine at position 8) are completely buried within the binding pocket, while two residues (serine at position 1, isoleucine at position 3) are largely buried, leaving three (asparagine at position 4, glutamic acid at position 6, and lysine at position 7) solvent-exposed.^79^ Prior mutagenesis studies have shown that alanine substitutions within **ovaWT** at positions P3, P5, and P8 reduce stability in the RMA-S assay, whereas substitutions at P4, P6, and P7 impair T cell recognition.^80^ These precedents align with our own observations: *N*-methylation at positions 2, 3, and 8 (residues buried in the binding groove) were found to be intolerable to modification. Although the effect may be primarily steric, changes in hydrogen bond engagement or *cis-trans* isomerization likely also contribute. Notably, although phenylalanine at position 5 serves as a critical anchor residue, backbone *N-*methylation at this site resulted in only a moderate reduction in pMHC stability. Taken together, these results highlight the more stringent structural constraints imposed on buried residues, standing in contrast to the potential plasticity of solvent-exposed positions.

Next, we constructed a peptoid library (**Fig. 4a**) by incorporating a single *N*-substituted glycine unit at varying positions along the **ovaWT** backbone (**ovaNalk1–8**) (**Fig. 4b**). Of note, for **ovaNalk1**, serine could not be directly translated into a peptoid residue, as β-hydroxy peptoid side chains are unstable during solid-phase synthesis due to favorable intramolecular cyclization;^81^ therefore, a homoserine analog was used instead. Unlike peptides, peptoids feature side chains attached to the backbone nitrogen rather than the α-carbon, allowing for the retention of the side chain identity while altering the spatial orientation.^82^ This backbone reconfiguration confers near-complete resistance to proteolysis, as the modified amide linkage is poorly recognized by proteolytic enzymes. Moreover, substitution at the backbone nitrogen eliminates the amide hydrogen bond donor, significantly reducing the energetic penalty of desolvation and thereby potentially enhancing membrane permeability. However, these advantages are accompanied by conformational consequences. The absence of backbone chirality and hydrogen bonding capacity results in a conformationally flexible (“floppy”) scaffold. Peptoids typically exhibit low rotational barriers about backbone torsion angles and high *cis/trans* amide heterogeneity, leading to a highly dynamic conformational ensemble.

**Figure 4.**
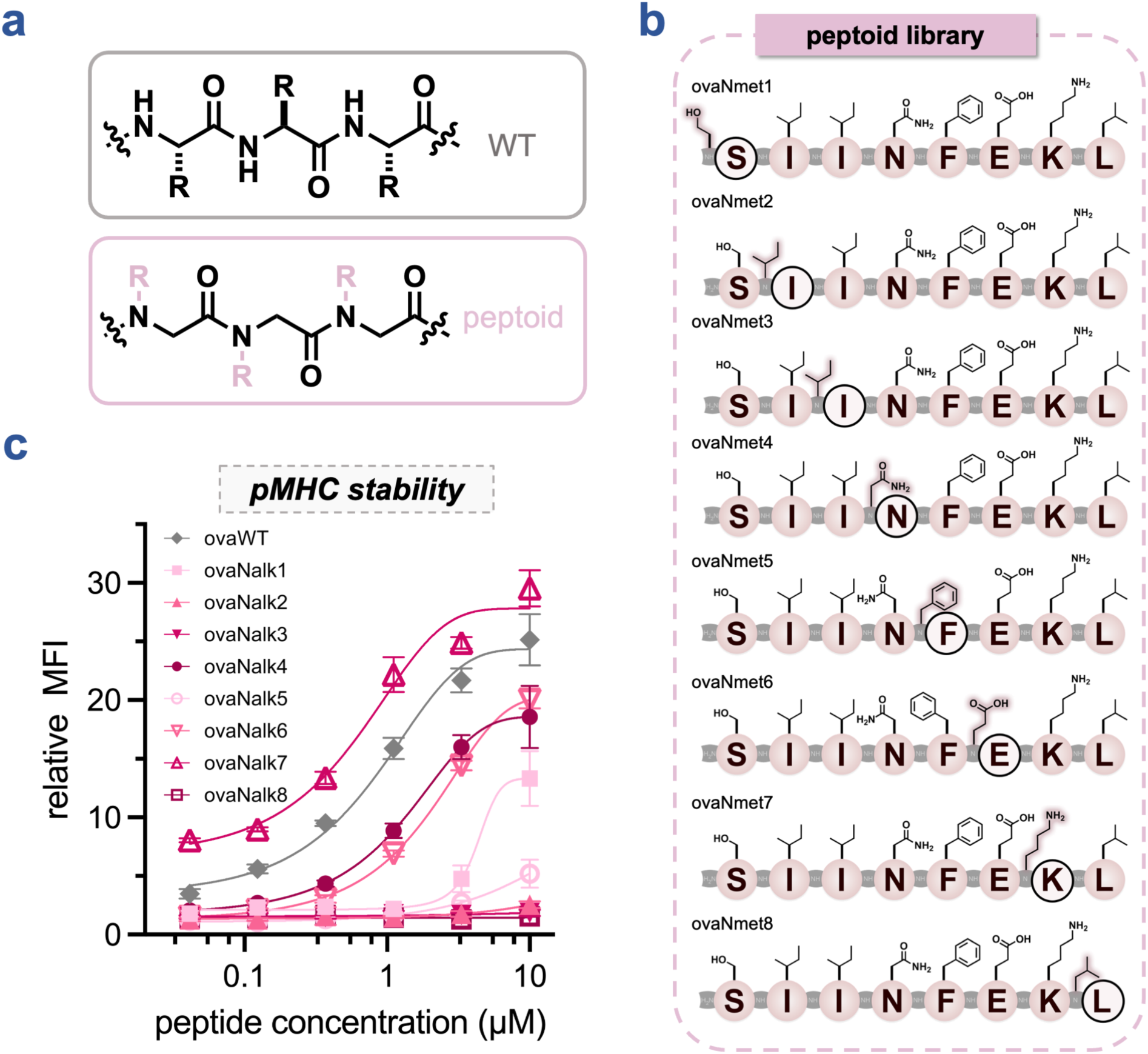
**(a)** Schematic depicting generic tripeptide with peptoid modifications. **(b)** Chemical structures of the singly substituted peptoid library. **(c)** Dose-response curves from flow cytometry analysis of the RMA-S stabilization assay. RMA-S cells were incubated with the indicated concentration of **ovaWT** and peptidomimetics from singly-substituted peptoid library. H-2K^b^ expression was analyzed via flow cytometry by APC anti-mouse H-2K^b^ antibody. MFI is the mean fluorescence intensity of the level of fluorescence relative to the DMSO control. Data are represented as mean ± SD (*n* = 3), and Boltzmann sigmoidal curves were fitted to the data using GraphPad Prism.

We evaluated our peptoid-modified series using the RMA-S assay (**Fig. 4c**). Prior work has demonstrated that even a single peptide-to-peptoid substitution at a solvent-exposed residue of an MHC-II ligand can substantially diminish binding, suggesting that peptoid analogs might not be tolerated.^83^ In contrast, we observed that peptoid substitutions at asparagine (**ovaNalk4**), glutamic acid (**ovaNalk6**), and lysine (**ovaNalk7**) preserved pMHC stability similar to unmodified **ovaWT**, whereas substitution at serine (**ovaNalk1**) caused a moderate decrease. Consistent with trends observed in the *N-*methylation library, peptoid modification at either isoleucine (**ovaNalk2** and **ovaNalk3**) completely abolished pMHC stability. Moreover, unlike the *N*-methylated series, peptoid substitution at the phenylalanine anchor residue (**ovaNalk5**) also eliminated pMHC complex stability, consistent with expectations from the crystal structure. Collectively, these results underscore a distinct structure-activity relationship in which solvent accessibility governs the permissibility of peptoid substitution, sharply contrasting with the stringent steric and conformational constraints imposed at primary anchor residues.

Finally, we synthesized a diastereomeric library (**Fig. 5a**) by inverting the stereochemistry at each residue of **ovaWT** (**ovaD1–8**) (**Fig. 5b**). Incorporation of D-amino acids confers substantial resistance to proteolysis by introducing a stereochemical mismatch with the chiral active sites of endogenous proteases, which have evolved to recognize L-residues. However, unlike *N*-methylation or peptoid substitutions, stereoinversion preserves the backbone amide hydrogen bond donor; thus, the energetic cost of desolvation remains chemically equivalent to that of the L-enantiomer.

**Figure 5.**
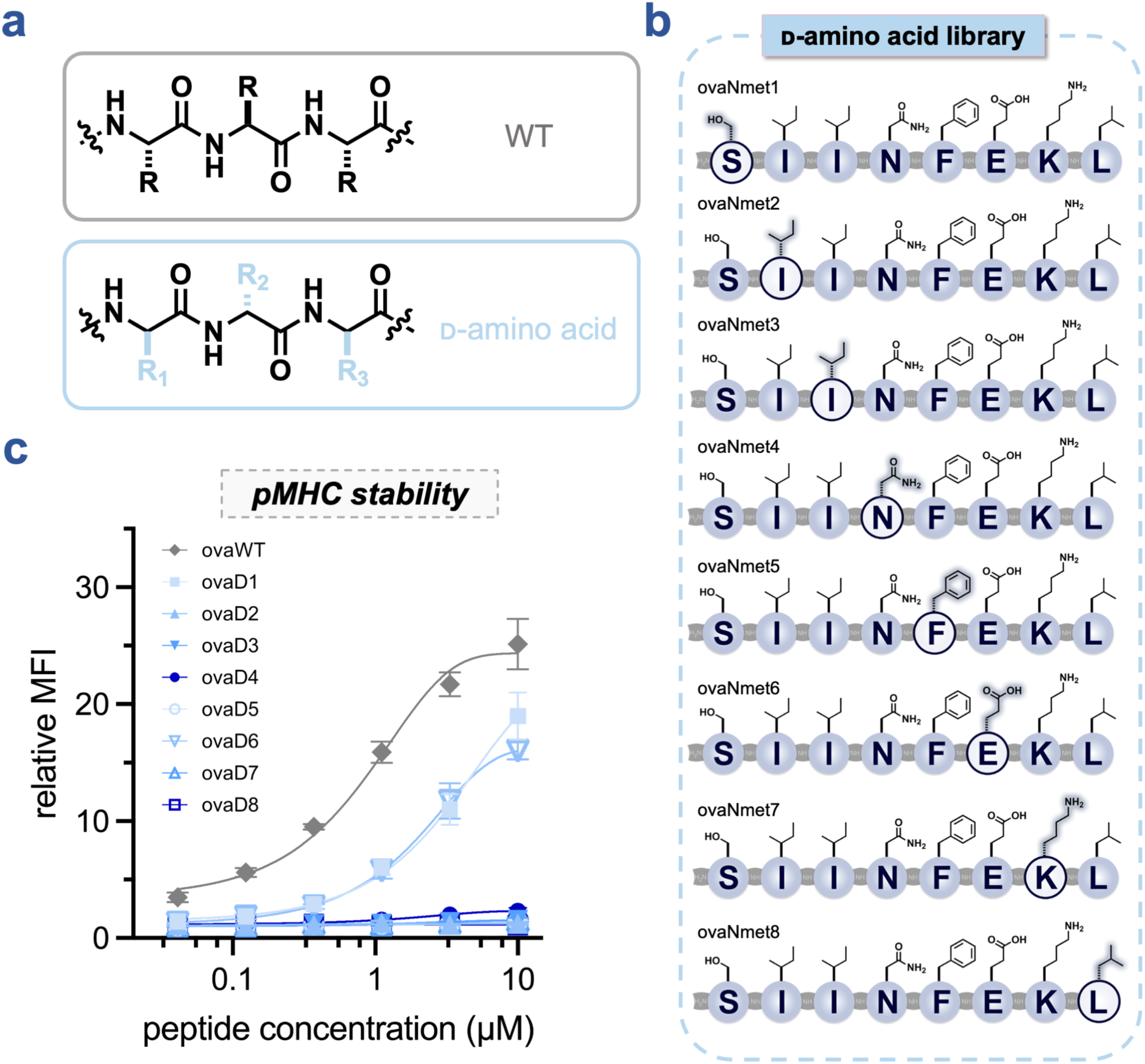
**(a)** Schematic depicting generic tripeptide with ᴅ-amino acid modifications. **(b)** Chemical structures of the singly substituted ᴅ-amino acid library. **(c)** Dose-response curves from flow cytometry analysis of the RMA-S stabilization assay. RMA-S cells were incubated with the indicated concentration of **ovaWT** and peptidomimetics from singly-substituted ᴅ-amino acid library. H-2K^b^ expression was analyzed via flow cytometry by APC anti-mouse H-2K^b^ antibody. MFI is the mean fluorescence intensity of the level of fluorescence relative to the DMSO control. Data are represented as mean ± SD (*n* = 3), and Boltzmann sigmoidal curves were fitted to the data using GraphPad Prism.

Although D-amino acid-containing peptides are often presumed to lack cell-based immunogenicity, this generalization warrants empirical validation. In the RMA-S system, stereoinversion at most positions resulted in complete loss of pMHC stability (**Fig. 5c**). Only the serine (**ovaD1**) and glutamic acid (**ovaD6**) variants retained partial activity, indicating limited positional tolerance to chirality reversal within the MHC-binding framework. However, even for stable complexes, the immunological outcome of stereoinversion is nuanced. It was previously demonstrated that while single D-substitutions typically abrogate recognition by T cell clones specific for the wild-type epitope, they can give rise to de novo CTL responses directed against D-isomer topology.^84^ Moreover, although peptides composed entirely of ᴅ-amino acids generally exhibit reduced MHC binding or diminished relative to their native L-counterparts, TCR cross-reactivity has enabled the identification of structurally unrelated all-D peptides that remain immunogenic.^85^ Thus, while direct stereochemical inversion of a known epitope may not preserve immunogenicity, the incorporation of ᴅ-amino acid building blocks represents a strategically valuable approach for the design of novel immunogenic peptides with improved stability and recognition properties.

### Effect of Peptidomimetic Substitutions on T Cell Activation

We then aimed to evaluate how peptidomimetic modifications to **ovaWT** influence TCR engagement. To do so, we employed the B3Z T cell hybridoma cell line, which expresses an OVA-specific TCR and a NFAT-LacZ reporter gene encoding for β-galactosidase under an IL-2-inducible promoter.^86^ When B3Z cells detect the SIINFEKL pMHC-I complex presented by RMA-S cells, IL-2 signaling induces the expression of β-galactosidase (**Fig. 6a**). This enzyme cleaves chlorophenol red-β-galactopyranoside (CPRG), producing a colorimetric shift that can be quantified spectrophotometrically. Because β-galactosidase expression is directly coupled to TCR signaling strength, the magnitude of the colorimetric response provides a quantitative measure of functional T cell activation, enabling the generation of dose-response curves for each peptidomimetic analog. By comparing these profiles to the native **ovaWT** baseline, we identified specific structural modifications that decouple pMHC stability from functional TCR engagement, thereby revealing positions where MHC binding and T cell signaling can be differentially modulated.

**Figure 6.**
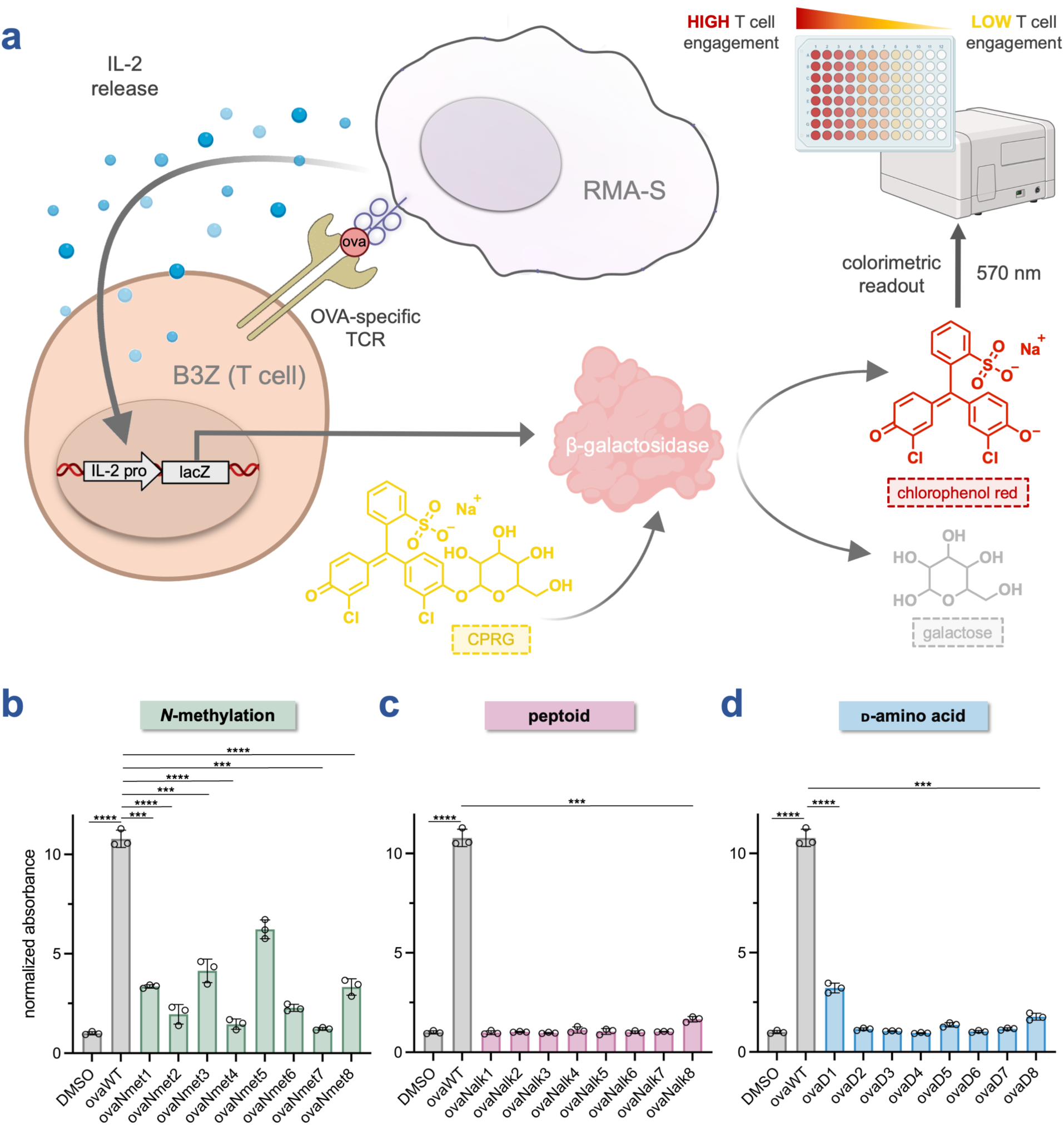
**(a)** Schematic illustrating the B3Z T cell activation assay. **(b – d)** RMA-S cells were incubated with 100 nM of **ovaWT** and peptidomimetics from singly-substituted **(b)** *N*-methylation, **(c)** peptoid, or **(d)** ᴅ-amino acid libraries for 1 h at 26°C. RMA-S cells were subsequently co-incubated with B3Z T cells for 6 h at 37°C. β-galactosidase expression was then measured via the conversion of the colorimetric reagent chlorophenol red-β-galactopyranoside (CPRG) on a plate reader at 570 nm. The data presented has been normalized to the absorbance of the DMSO control. Data are represented as mean ± SD (*n* = 3). *P*-values were determined by a two-tailed *t*-test (*** *p* < 0.001, **** *p* < 0.0001).

Our data revealed that, consistent with trends observed for pMHC stability, several *N*-methylated variants were still capable of activating SIINFEKL-specific T cells at a potent concentration of 100 nM (**Fig. 6b** and **S1**). Notably, **ovaNmet5**, *N*-methylated at the phenylalanine residue, retained appreciable TCR engagement. *N*-methylation at serine (**ovaNmet1**), the second isoleucine (**ovaNmet3**), and leucine (**ovaNmet8**) also produced moderate levels of T cell recognition. Collectively, these findings validate backbone N-methylation as a robust strategy for peptide engineering, identifying a permissible structural window where even anchor-modified variants such as **ovaNmet5** can maintain the precise molecular geometry required for potent T cell activation. In contrast, *N*-methylation at most other positions, particularly lysine (**ovaNmet7**) and asparagine (**ovaNmet4**), was highly detrimental to TCR activation. Consistent with these findings, previous work in our lab has demonstrated that lysine modifications in **ovaWT**,^87^ including enzymatic lysine methylation,^88^ disrupt TCR recognition, likely due to altered charge distribution and hydrophobicity. As indicated by the SIINFEKL-H-2K^b^ crystal structure, these solvent-exposed residues appear especially sensitive to modification, supporting the structural relevance of side chain accessibility in TCR binding. Taken together, these data delineate a clear structural boundary for peptidomimetic design: while the TCR interface can surprisingly accommodate backbone modifications at specific anchor sites, it remains exquisitely sensitive to perturbations at solvent-exposed residues essential for molecular recognition.

Since *N*-methylation demonstrated that the pMHC interface can accommodate backbone alkylation at specific sites, we projected that peptoid substitutions might offer a similar opportunity for modulation. Although peptoids introduce a more significant structural alteration by relocating the side chain to the nitrogen atom, they share the core feature of *N*-alkylation. We therefore tested the corresponding library of peptoid analogs to determine if the permissive structural windows identified for *N*-methylated peptides are conserved when the side chain itself is shifted to the backbone. When evaluating the peptoid-modified series in T cell engagement, the results were striking: none of the variants exhibited measurable TCR activation at any concentration tested, with the sole exception of **ovaNalk8**, which elicited weak response only at the highest concentration (1 µM) (**Fig. 6c** and **S2**). These findings suggest that peptoid backbones are unable to sufficiently replicate the native peptide conformation within the MHC binding groove to permit productive TCR engagement. While peptoids preserve the chemical identity of the side chain, our findings indicate that the shift in backbone-to-side chain registration could potentially disrupts the precise geometry required for TCR recognition. Furthermore, the increased conformational mobility of the *N*-substituted side chain may impose an entropic penalty that destabilizes the induced-fit complex required for signaling.

Finally, we sought to determine whether the unique protease-resistance profile of stereochemically inverted peptides could be translated into functional immunogenicity. We therefore examined the effect of stereochemical inversion on T cell activation (**Fig. 6d** and **S3**). Similar to the peptoid series, conversion of each residue from the L- to the D-stereoisomer almost completely abolished TCR engagement for every diastereomeric analog across all concentrations. Exceptions were rare and weak: only **ovaD1** (D-Ser) retained minor activity, while the **ovaD5** (D-Phe) and **ovaD8** (D-Leu) variants exhibited low-level recognition only at the highest concentration (1 µM). This nearly universal loss of function highlights the strict geometric constraints of the T cell engagement for specifically evolved TCR pairs. Even when specific side chains like phenylalanine (P5) or leucine (P8) are critical for binding, their inverted presentation likely misaligns the antigenic surface, preventing the precise induced-fit mechanism required for robust TCR triggering. Overall, these results indicate that TCR recognition is predominantly dictated by interactions with the peptide side chains, while limited tolerance exists for backbone modifications. The data support a model in which TCR recognition depends on the native peptide backbone conformation and precise side chain orientation within the pMHC complex. Even subtle alterations in backbone geometry can disrupt the integrity of the pMHC-TCR interface. In this sense, the immunological synapse is critical, and the TCR engages the peptide-MHC complex as a single, integrated structural unit, rather than discrete molecular components.^2^

Recognizing that an all-ᴅ peptide would be unlikely to retain the immunogenic properties of its native L-counterpart owing to the inversion of the peptide backbone orientation, we next investigated a retro-inverso variant of SIINFEKL (**ovaRI**). In the retro-inverso strategy, the amino acid sequence is reversed and the chirality of each residue inverted. This design is intended to preserve the approximate spatial projection of the side chains while reversing the directionality of the peptide backbone. Although this maneuver can reproduce aspects of the parent side chain topology, it inverts the orientation of the backbone amide hydrogen bond donors and carbonyl acceptors relative to the native epitope. Retro-inverso analogs of antigenic peptides have previously been explored in both MHC-I and MHC-II systems, with mixed outcomes reported.^44–48^ In principle, if peptide-MHC binding were governed predominantly by side-chain interactions, preservation of side-chain orientation might allow retention of pMHC stability and downstream TCR activation. However, if productive binding requires coordinated contributions from both side chains and the native backbone hydrogen bonding geometry, retro-inversion would be predicted to impair complex formation. To interrogate this mechanistic distinction, we evaluated **ovaRI** in both assays. In the RMA-S stabilization assay, **ovaRI** failed to generate detectable signal relative to **ovaWT**, indicating an absence of stable pMHC complex formation (**Fig. S4a**). Consistent with this finding, the B3Z assay demonstrated only a minimal increase in activation (**Fig. S4b**). Taken together, these data indicate that preservation of side-chain topology alone is insufficient to sustain MHC binding in the context of SIINFEKL. Instead, stable complex formation appears to require the cooperative alignment of both side-chain positioning and native backbone orientation, including proper hydrogen bonding geometry and peptide register within the MHC groove. The failure of the retro-inverso variant therefore supports a model in which MHC binding is jointly backbone- and side chain-dependent, rather than dominated by either component in isolation.

Interestingly, certain discrepancies between the RMA-S and B3Z assays warrant further consideration. In several cases, peptides that appeared incapable of stabilizing MHC in the RMA-S assay nonetheless elicited measurable TCR responses in the B3Z assay. At first glance, this observation seems counterintuitive, since MHC binding is the prerequisite step for TCR recognition and subsequent T cell activation. The most likely explanation lies in the differing sensitivities and readouts of the two assays. The RMA-S assay primarily reflects the abundance and overall stability of pMHC complexes presented on the cell surface, whereas the B3Z assay reports on the functional quality of those complexes in engaging the TCR. It is therefore possible that certain modified peptides form a limited number of conformationally favorable complexes that remain below the detection threshold of the RMA-S assay yet are sufficient to activate T cells bearing high-affinity TCRs. This apparent discrepancy highlights that immune recognition depends not solely on the quantity of pMHC complexes, but also on the structural and kinetic quality of the pMHC-TCR interaction.

### Effect of Peptidomimetic Substitutions on Permeability

Building on this framework, we aimed to measure the intracellular accumulation of modified peptides that have the potential to function as peptide vaccines. By quantitatively assessing cytosolic delivery in parallel with MHC presentation, we will define how specific peptidomimetic modifications influence not only cellular permeability but also the abundance of antigenic peptides available for immune recognition. This integrated analysis can potentially start the path towards establishing design principles for optimizing peptide stability, intracellular access, and presentation efficiency, thereby advancing the rational development of next-generation peptide-based vaccines. To the best of our knowledge, this represents the first integration of a definitive cytosolic demarcation strategy with direct evaluation of MHC presentation. For this approach, we employed the Chloroalkane HaloTag Azide-based Membrane Penetration (CHAMP) assay, a method developed in our laboratory which we have optimized for use first in bacterial cells^89^ and, more recently, in mammalian systems.^71^ CHAMP leverages HaloTag-expressing cells in combination with strain-promoted azide-alkyne cycloaddition (SPAAC) chemistry to quantify the presence of azide-tagged compounds within the cytosol (**Fig. 7a**). Specifically, intracellular chloroalkane-linked dibenzoazacyclooctyne (DBCO) landmarks react covalently with azide-bearing molecules that successfully reach the cytosol, enabling assessment of accumulation. Unlike traditional fluorophore-based tracking approaches, which often conflate general cell association (such as membrane binding or endosomal entrapment) with internalization, CHAMP utilizes intracellular DBCO landmarks that exclusively react with molecules that have genuinely accessed the cytosol.

**Figure 7.**
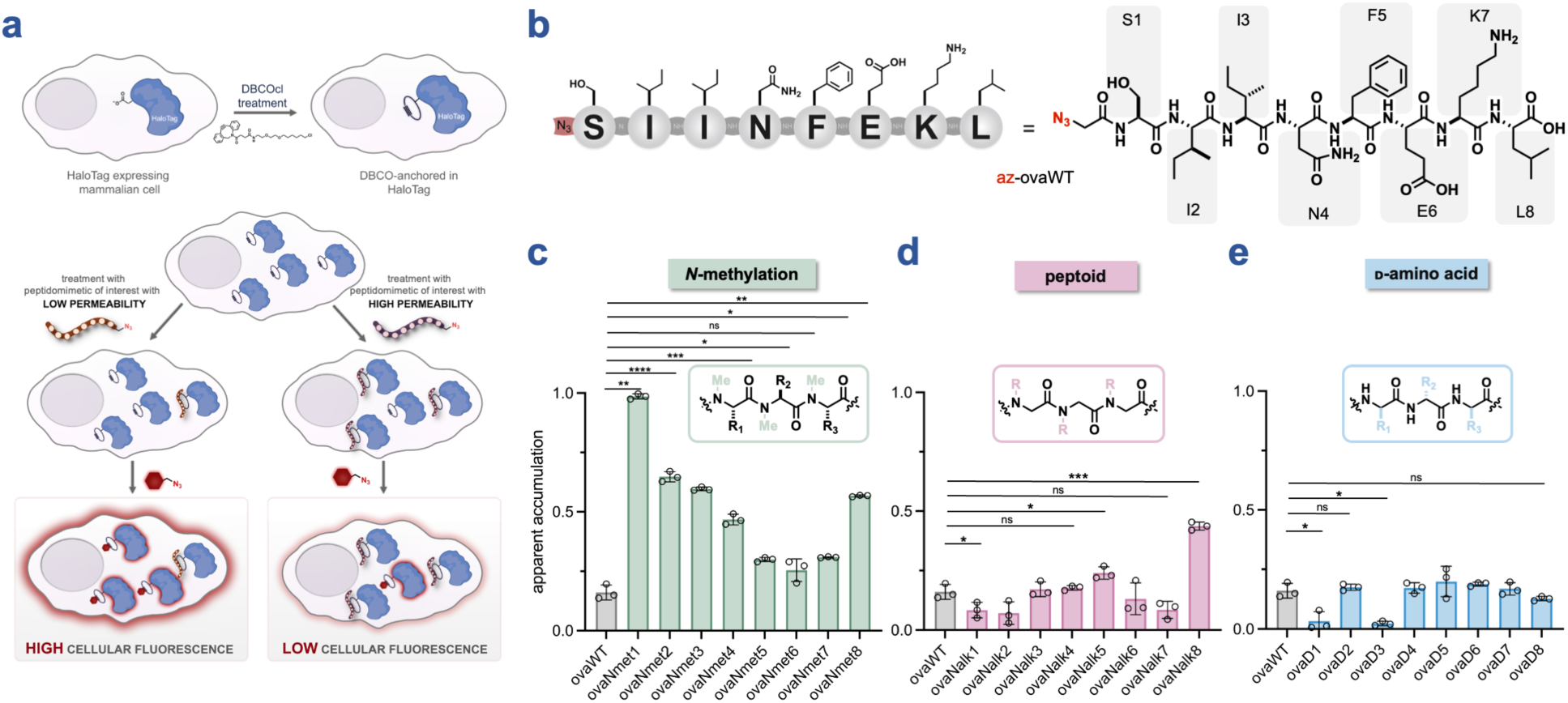
**(a)** Schematic illustrating the CHAMP assay in mammalian cells. Cells expressing HaloTag in the cytosol are first treated with a chloroalkane-modified strained alkyne, enabling covalent installation of DBCO landmarks within the cytosol. Subsequently, an azide-tagged molecule is introduced in a “pulse” step, reacting with the strained alkyne via a SPAAC reaction. Molecules exhibiting high cytosolic entry consume more DBCO sites, resulting in fewer available reactive handles during the subsequent fluorescent azide “chase” step. Thus, molecules with greater cytosolic accumulation yield lower cellular fluorescence. **(b)** Chemical structure of **az-ovaWT. (c – e)** Flow cytometry analysis of the CHAMP assay. HaloTag-expressing HeLa cells were pulsed with 50 µM of **az-ovaWT** or peptidomimetics from azide-tagged singly-substituted **(c)** *N*-methylation, **(d)** peptoid, or **(e)** ᴅ-amino acid libraries for 24 h, and chased with 50 µM of TMRaz. The data are presented such that higher fold change is indicative of higher relative accumulation. Data are represented as mean ± SD (*n* = 3). *P*-values were determined by a two-tailed *t*-test (ns = not significant, * *p* < 0.05, ** *p* < 0.01, *** *p* < 0.001, **** *p* < 0.0001).

To enable CHAMP-based evaluation across our libraries of **ovaWT** analogs, we installed an *N*-terminal azide handle into the parent peptides (**az-ovaWT**, **Fig. 7b**) as well as into each corresponding analog from the three-modification series (**az-ovaNmet1-8**, **az-ovaNalk1-8**, **az-ovaD1-8**, **Fig. S5**). This uniform modification allowed direct comparison of cytosolic accumulation across all variants under identical experimental conditions. Following a 24-hour incubation with each peptidomimetic analog, most variants exhibited little to no statistically significant difference in fluorescence signal relative to **az-ovaWT**, with uniformly low apparent accumulation observed across these samples (including **az-ovaWT** itself). Notably, however, several members of the *N-*methylation library displayed enhanced apparent accumulation (**Fig. 7c**). In particular, **az-ovaNmet1**, corresponding to backbone *N*-methylation at the *N*-terminal serine, exhibited a robust increase in accumulation after the incubation period. More broadly, sequentially moving the *N*-methyl group along the peptide backbone from the *N*-terminus toward the *C*-terminus resulted in progressively decreased accumulation, reaching non-statistically significant levels by **az-ovaNmet6**, which contains *N*-methylation at the phenylalanine residue. One exception to this trend was **az-ovaNmet8**, corresponding to modification at the *C-*terminal leucine, which again displayed increased apparent accumulation. These findings are consistent with prior reports demonstrating that backbone *N-*methylation can enhance membrane permeability in mammalian systems.^67–69^ As a result, the peptide becomes less hydrophilic and more readily sheds its solvation shell, a step that favors passive diffusion through nonpolar environments.

Compared to the *N*-methylated peptides, the peptoid library exhibited substantially lower overall apparent accumulation (**Fig. 7d**). However, **az-ovaNalk8**, corresponding to the *N*-substituted glycine unit at the *C-*terminal leucine position, showed a moderate increase in apparent accumulation (**Fig. 7d**). Notably, was also the only member of the peptoid library that elicited an above-background signal in the B3Z T cell activation assay, although as previously mentioned, the response remained very weak and was observed only at the highest concentration tested (**Fig. S2**). Consistent with their chemical properties, peptoids (like *N-*methylated peptides) lack the backbone amide hydrogen bond donor, which may improve their permeability relative to canonical peptides.^89–92^ In contrast, all members of the ᴅ-amino acid library exhibited near-background levels of accumulation, comparable to (and in some cases lower than **az-ovaWT**. Collectively, these results suggest that backbone *N-*methylation, and to a much lesser extent peptoid substitution under specific circumstances, can enhance cellular permeability of antigenic peptides when introduced at particular positions. Importantly, many of the *N-*methylated variants also retained appreciable T cell activation, as demonstrated earlier, indicating that permeability and immunogenic function can be tuned simultaneously within this scaffold.

### Design and Analysis of Multiply Substituted SIINFEKL Derivatives

Reasoning that single-residue substitutions might yield only incremental pharmacokinetic improvements, we next engineered multiply substituted SIINFEKL variants with the goal of maximizing MHC-I binding stability and minimal loss of immunogenicity. Candidate modifications were strictly filtered for their ability to retain dual functionality: robust MHC-I stabilization (RMA-S assay) and potent T cell activation (B3Z assay). This rational design strategy yielded two analogs: **ovaDiMod**, featuring an inverted stereocenter at serine and backbone *N*-methylation at phenylalanine; and **ovaTriMod**, which appends an additional stereochemical inversion at the *C*-terminal leucine (**Fig. 8a**). Notably, the triply modified scaffold spatially distributes alterations across the peptide backbone, allowing us to probe the cumulative impact of simultaneous *N*-terminal, core, and *C*-terminal modifications. We incorporated the *C*-terminal inversion despite the reduced potency of the single variant (**ovaD8**) at 100 nM, as dose–response profiling confirmed significant TCR engagement at 1 µM (**Fig. S3**). This ensured that our most heavily modified variant balanced maximal proteolytic resistance with preserved antigenicity.

**Figure 8.**
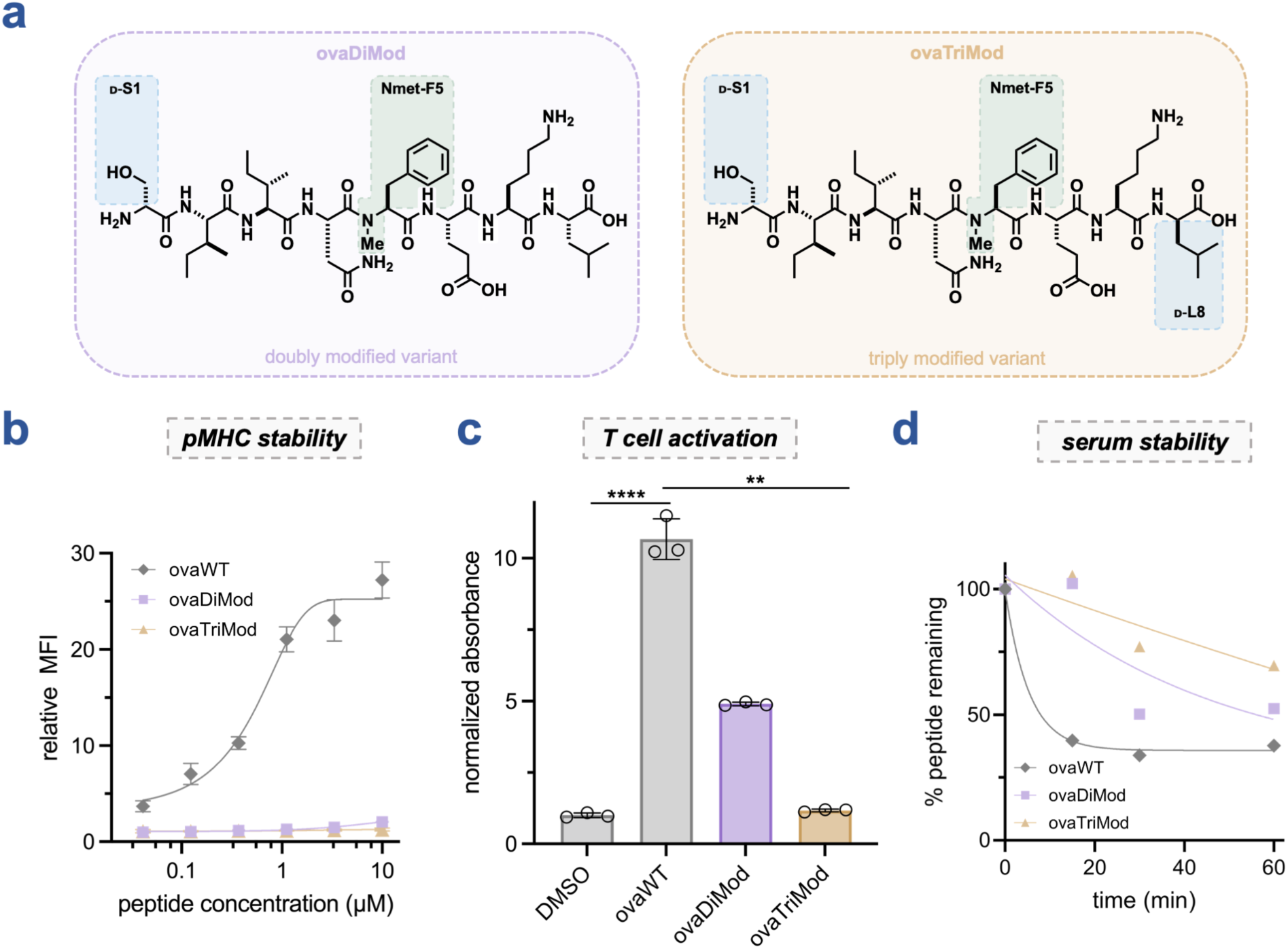
**(a)** Chemical structures of **ovaDiMod** and **ovaTriMod** with their structural modifications highlighted. **(b)** Dose-response curves from flow cytometry analysis of the RMA-S stabilization assay. RMA-S cells were incubated with the indicated concentration of **ovaWT**, **ovaDiMod**, or **ovaTriMod**. H-2K^b^ expression was analyzed via flow cytometry by APC anti-mouse H-2K^b^ antibody. MFI is the mean fluorescence intensity of the level of fluorescence relative to the DMSO control. Data are represented as mean ± SD (*n* = 3), and Boltzmann sigmoidal curves were fitted to the data using GraphPad Prism. **(c)** RMA-S cells were incubated with 100 nM **ovaWT**, **ovaDiMod**, or **ovaTriMod** for 1 h at 26°C. RMA-S cells were subsequently co-incubated with B3Z T cells for 6 h at 37°C. β-galactosidase expression was then measured via the conversion of the colorimetric reagent chlorophenol red-β-galactopyranoside (CPRG) on a plate reader at 570 nm. The data presented has been normalized to the absorbance of the DMSO control. Data are represented as mean ± SD (*n* = 3). *P*-values were determined by a two-tailed *t*-test (** *p* < 0.01, **** *p* < 0.0001). **(d)** Comparison of serum stability of **ovaWT**, **ovaDiMod**, and **ovaTriMod** incubated in mouse serum at 37°C over 1 h. One-phase decay curves were fitted to the data using GraphPad Prism.

With these two multiply modified variants in hand, we performed RMA-S and B3Z assays to assess their effects on pMHC stability and T cell activation, respectively. In the RMA-S assay, both the doubly modified (**ovaDiMod**) and triply modified (**ovaTriMod**) peptides exhibited a complete loss of detectable pMHC stability across all concentrations tested (**Fig. 8b**). This result was notable given that the two single substitutions comprising **ovaDiMod** (backbone *N*-methylation at Phe5 as **ovaNmet5**, and stereochemical inversion at Ser1 as **ovaD1**) each retained appreciable pMHC stability when evaluated individually (**Fig. 3c and 5c**). Incorporation of the additional stereochemical inversion at the *C*-terminal leucine in **ovaTriMod** neither rescued nor further diminished this response, although this was not entirely unexpected, consistent with the poor performance of the corresponding single variant (**ovaD8**) in the initial RMA-S analysis.

Despite the loss of measurable pMHC stability in these multiply modified variants, T cell activation remained a primary metric of interest, as it is more representative of productive immune recognition even when overall pMHC abundance may be low or falls below the detection threshold of the RMA-S stabilization assay. Encouragingly, in the B3Z assay, **ovaDiMod** elicited a ∼5-fold increase in T cell activation relative to background at 100 nM (**Fig. 8c**), an intermediate response between those of its constituent single variants (**ovaNmet5** and **ovaD1**, which produced ∼6-fold and ∼4-fold increases, respectively) (**Fig. 6b and 6d**). Moreover, at the highest concentration tested (1 µM), **ovaDiMod** approached the level of activation observed for **ovaWT**, and notably exceeded the responses of either single variant at this concentration (**Fig. S6**). In contrast, the addition of the third modification in **ovaTriMod** abolished T cell activation, mirroring its lack of pMHC stability. These data identify the *C*-terminal anchor as a site of exquisite stereochemical sensitivity, where modification disrupts both MHC binding and productive TCR engagement. Critically, the divergent fates of these variants underscore the non-additive nature of peptidomimetic design: the successful combination of modifications in **ovaDiMod**, versus the functional loss of **ovaTriMod**, demonstrates that individually tolerated substitutions do not guarantee preservation of function when multiplexed. Consequently, **ovaDiMod** was selected as the optimal lead candidate for further physiological characterization.

Finally, we interrogated how the peptidomimetic substitutions with **ovaWT** impacted their resistance to serum proteases. In this context, we examined only our derivatives containing multiple modifications, as a substitution at any given single residue is unlikely to offer significantly improved protection from proteolysis across the entire length of the peptide. Additionally, considering the activity of both *N*- and *C*-terminal exopeptidases,^93^ we felt that our multiply modified variants bearing ᴅ-amino acids at the termini would be promising for investigating this impact. Assessment of metabolic stability was performed by treatment with mouse serum and it revealed rapid degradation of **ovaWT**, with only 40% of the initial peptide remaining after 15 minutes of serum incubation and no further decline over the remainder of the hour (**Fig. 8d**). In contrast, **ovaDiMod** remained stable during the first 15 minutes, followed by a sharp drop to ∼50% remaining after 30 minutes, plateauing thereafter. **ovaTriMod** exhibited the greatest stability, retaining nearly 100% of the peptide after 15 minutes, 77% after 30 minutes, and 70% after one hour. These results align with prior findings that backbone *N-*methylation markedly enhances serum stability.^94, 95^ In **ovaDiMod** and **ovaTriMod**, backbone *N-*methylation at the phenylalanine at position 5 likely imparts steric hindrance that restricts protease access to the cleavage site and simultaneously removes a backbone amide hydrogen essential for protease recognition through hydrogen bonding.^96^ Furthermore, because proteases display strict stereospecificity, inversion of terminal stereocenters is a well-established strategy to impede proteolysis.^97, 98^ Protection of the *N*-terminus in **ovaDiMod** likely reduces susceptibility to aminopeptidases, whereas protection of both termini in **ovaTriMod** additionally limits carboxypeptidase activity. Collectively, these data suggest that the combined effects of backbone *N*-methylation and terminal stereochemical inversion confer multi-level resistance to serum proteases, leading to the progressive increase in stability observed from **ovaWT** to **ovaTriMod**.

## CONCLUSIONS

In this study, we systematically interrogated how distinct peptidomimetic modifications influence the key molecular and cellular determinants of antigen presentation using the well-defined SIINFEKL-H-2K^b^ model system. By integrating assays for MHC-I stabilization, TCR activation, cellular permeability, and serum stability, we delineate how backbone chemistry, stereochemistry, and residue positioning collectively shape peptide immunogenicity and pharmacokinetic behavior. Our findings demonstrate that tolerance to peptidomimetic modification is highly position dependent. Certain backbone *N*-methylations were compatible with MHC binding and TCR recognition and were associated with enhanced cellular accumulation, whereas peptoid substitutions and stereochemical inversion were generally less well tolerated for productive TCR engagement and intracellular accumulation. Importantly, discrepancies between MHC stabilization and T cell activation highlight that functional immune recognition depends not only on the abundance of pMHC complexes, but also on their conformational and kinetic properties.

Extension of these principles to multiply modified peptides revealed that combinatorial effects are non-additive. While a doubly modified peptide retained substantial T cell activation alongside improved serum stability, the introduction of an additional modification abolished immune recognition despite further gains in protease resistance. These results underscore the inherent trade-offs between enhancing pharmacokinetic properties and preserving immunogenic function. Collectively, this study establishes principles for peptidomimetic antigen design, emphasizing the need to balance stability and permeability with the stringent structural requirements of MHC-I presentation and productive TCR engagement. More broadly, our findings provide a framework for rationally tuning peptide-based vaccines while maintaining the molecular features necessary for effective immune recognition.

## ACKNOWLEDGEMENT

This study was supported by the NIH grant R35GM124893 (M.M.P.)

## SUPPORTING INFORMATION

Additional figures, tables, and materials/methods are included in the supporting information file.

