## Supporting Information for "Mapping the Modification Landscape of MHC-I Epitopes: A Framework for Immunogenic Peptidomimetic Antigen Design"

### TABLE OF CONTENTS


| | |
|---|---|
| **SUPPLEMENTARY FIGURES** | **S3–S8** |
| **Figure S1.** Dose-response curves from B3Z T cell activation assay for *N*-methylation library. | S3 |
| **Figure S2.** Dose-response curves from B3Z T cell activation assay for peptoid library. | S4 |
| **Figure S3.** Dose-response curves from B3Z T cell activation assay for D-amino acid library. | S5 |
| **Figure S4.** RMA-S and B3Z assay for **ovaRI** | S6 |
| **Figure S5.** Chemical structures of azide-tagged peptidomimetic libraries | S7 |
| **Figure S6.** Dose-response curves from B3Z T cell activation assay for multiply-substituted variants **ovaDiMod** and **ovaTriMod** | S8 |
| **MATERIALS AND METHODS** | **S9–S12** |
| **Materials** | **S9–S10** |
| **Experimental Methods** | **S11–S12** |
| Mammalian Cell Culture | S11 |
| RMA-S Stabilization Assay | S11 |
| B3Z T Cell Activation Assay | S11 |
| Chloroalkane HaloTag Azide-based Membrane Penetration (CHAMP) Assay | S12 |
| Serum Stability Assay | S12 |
| **SYNTHESIS AND CHARACTERIZATION OF PEPTIDOMIMETICS** | **S13–S68** |
| General procedure for the solid-phase synthesis of peptidomimetics | S13 |
| Solid-phase peptide synthesis of *N*-methylation and D-amino acid libraries | S13 |
| Solid-phase peptoid synthesis of *N*-alkylation library | S14 |
| **Characterization of Peptidomimetics** | **S15–S69** |
| **ovaWT** (SIINFEKL) | S16 |
| *N*-methylation library (**ovaNmet1** – **ovaNmet8**) | S17–S24 |
| Peptoid library (**ovaNalk1** – **ovaNalk8**) | S25–S32 |
| D-amino acid library (**ovaD1** – **ovaD8**) | S33–S40 |
| **ovaRI** (retro-inverso SIINFEKL) | S41 |
| Multiply substituted variants (**ovaDiMod** and **ovaTriMod**) | S42–S43 |
| **az-ovaWT** (azide-modified SIINFEKL) | S44 |
| Azide-modified *N*-methylation library (**az-ovaNmet1** – **az-ovaNmet8**) | S45–S52 |
| Azide-modified peptoid library (**az-ovaNalk1** – **az-ovaNalk8**) | S53–S60 |
| Azide-modified D-amino acid library (**az-ovaD1** – **az-ovaD8**) | S61–S68 |
| **REFERENCES** | **S69** |

### SUPPLEMENTARY FIGURES

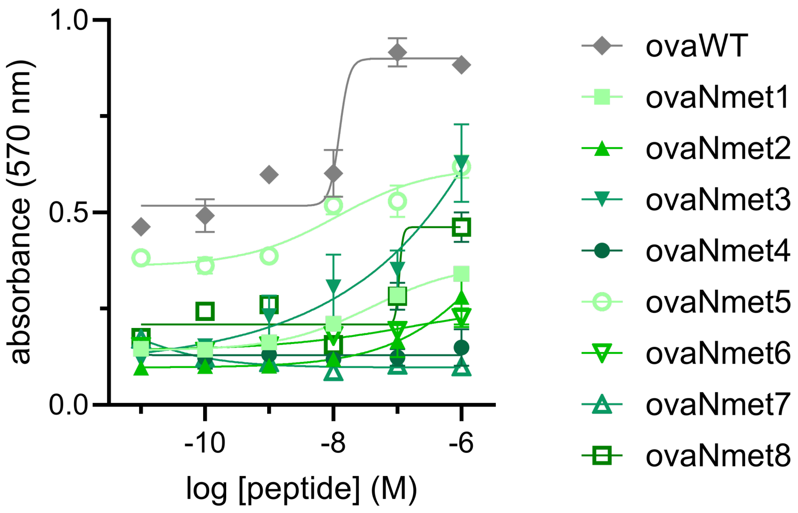


**Figure S1.** Dose-response curves from B3Z T cell activation assay for *N*-methylation library (full concentration scan from which the 100 nM data in **Fig. 6b** was obtained). RMA-S cells were incubated with the indicated concentrations of **ovaWT** and peptidomimetics from the singly-substituted *N*-methylation library for 1 h at 26°C. RMA-S cells were subsequently co-incubated with B3Z T cells for 6 h at 37°C. β-galactosidase expression was then measured via the conversion of the colorimetric reagent chlorophenol red-β-galactopyranoside (CPRG) on a plate reader at 570 nm. The data presented has been normalized to the absorbance of the DMSO control. Data are represented as mean ± SD (*n* = 3), and Boltzmann sigmoidal curves were fitted to the data using GraphPad Prism.

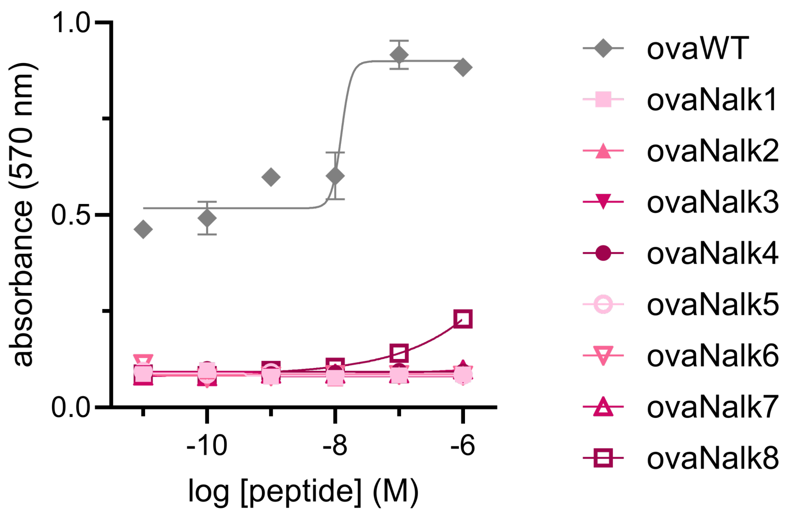


**Figure S2.** Dose-response curves from B3Z T cell activation assay for peptoid library (full concentration scan from which the 100 nM data in **Fig. 6c** was obtained). RMA-S cells were incubated with the indicated concentrations of **ovaWT** and peptidomimetics from the singly-substituted peptoid library for 1 h at 26°C. RMA-S cells were subsequently co-incubated with B3Z T cells for 6 h at 37°C. β-galactosidase expression was then measured via the conversion of the colorimetric reagent chlorophenol red-β-galactopyranoside (CPRG) on a plate reader at 570 nm. The data presented has been normalized to the absorbance of the DMSO control. Data are represented as mean ± SD ($n$ = 3), and Boltzmann sigmoidal curves were fitted to the data using GraphPad Prism.

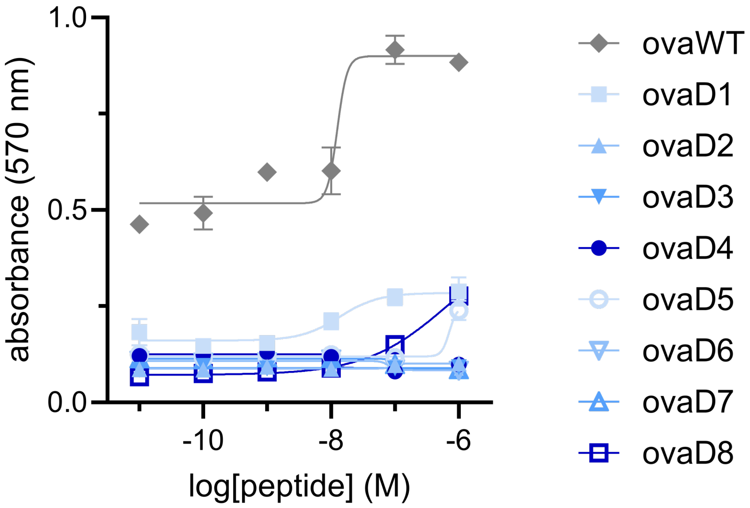


**Figure S3.** Dose-response curves from B3Z T cell activation assay for D-amino acid library (full concentration scan from which the 100 nM data in **Fig. 6d** was obtained). RMA-S cells were incubated with the indicated concentrations of **ovaWT** and peptidomimetics from the singly-substituted D-amino acid library for 1 h at 26°C. RMA-S cells were subsequently co-incubated with B3Z T cells for 6 h at 37°C. β-galactosidase expression was then measured via the conversion of the colorimetric reagent chlorophenol red-β-galactopyranoside (CPRG) on a plate reader at 570 nm. The data presented has been normalized to the absorbance of the DMSO control. Data are represented as mean ± SD (*n* = 3), and Boltzmann sigmoidal curves were fitted to the data using GraphPad Prism.

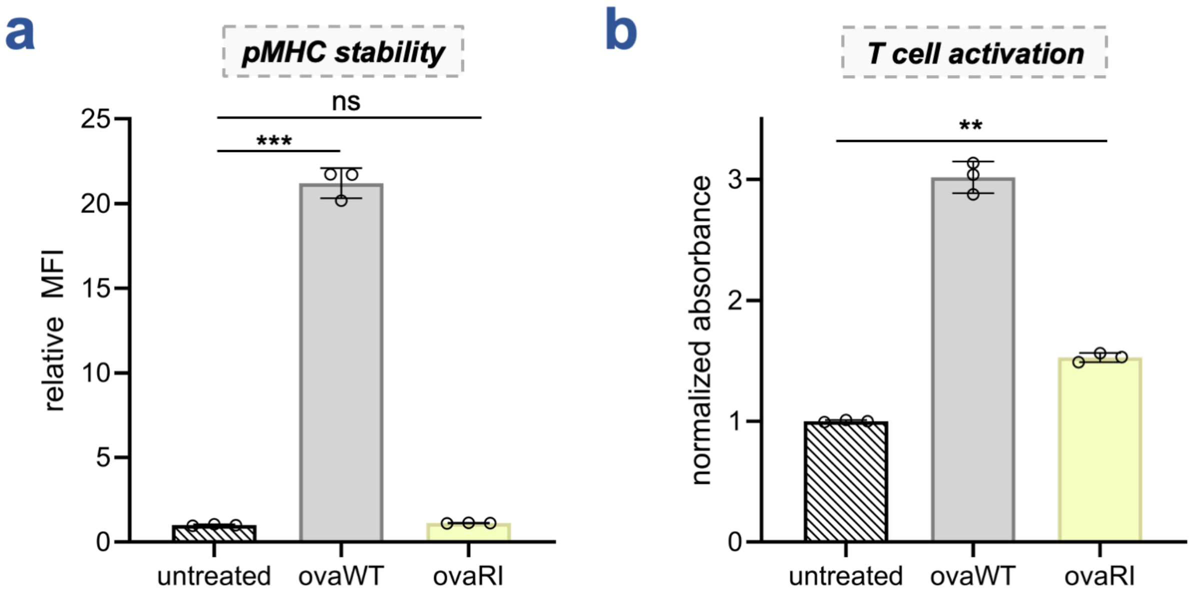


**Figure S4. (a)** Flow cytometry analysis of the RMA-S stabilization assay. RMA-S cells were incubated with 20 μM of **ovaWT** and **ovaRI**. H-2K$^b$ expression was analyzed via flow cytometry by APC anti-mouse H-2K$^b$ antibody. MFI is the mean fluorescence intensity of the level of fluorescence relative to the DMSO control. Data are represented as mean ± SD (*n* = 3). **(b)** RMA-S cells were incubated with 20 μM of **ovaWT** and **ovaRI** for 1 h at 26°C. RMA-S cells were subsequently co-incubated with B3Z T cells for 6 h at 37°C. β-galactosidase expression was then measured via the conversion of the colorimetric reagent chlorophenol red-β-galactopyranoside (CPRG) on a plate reader at 570 nm. The data presented has been normalized to the absorbance of the DMSO control. Data are represented as mean ± SD (*n* = 3).

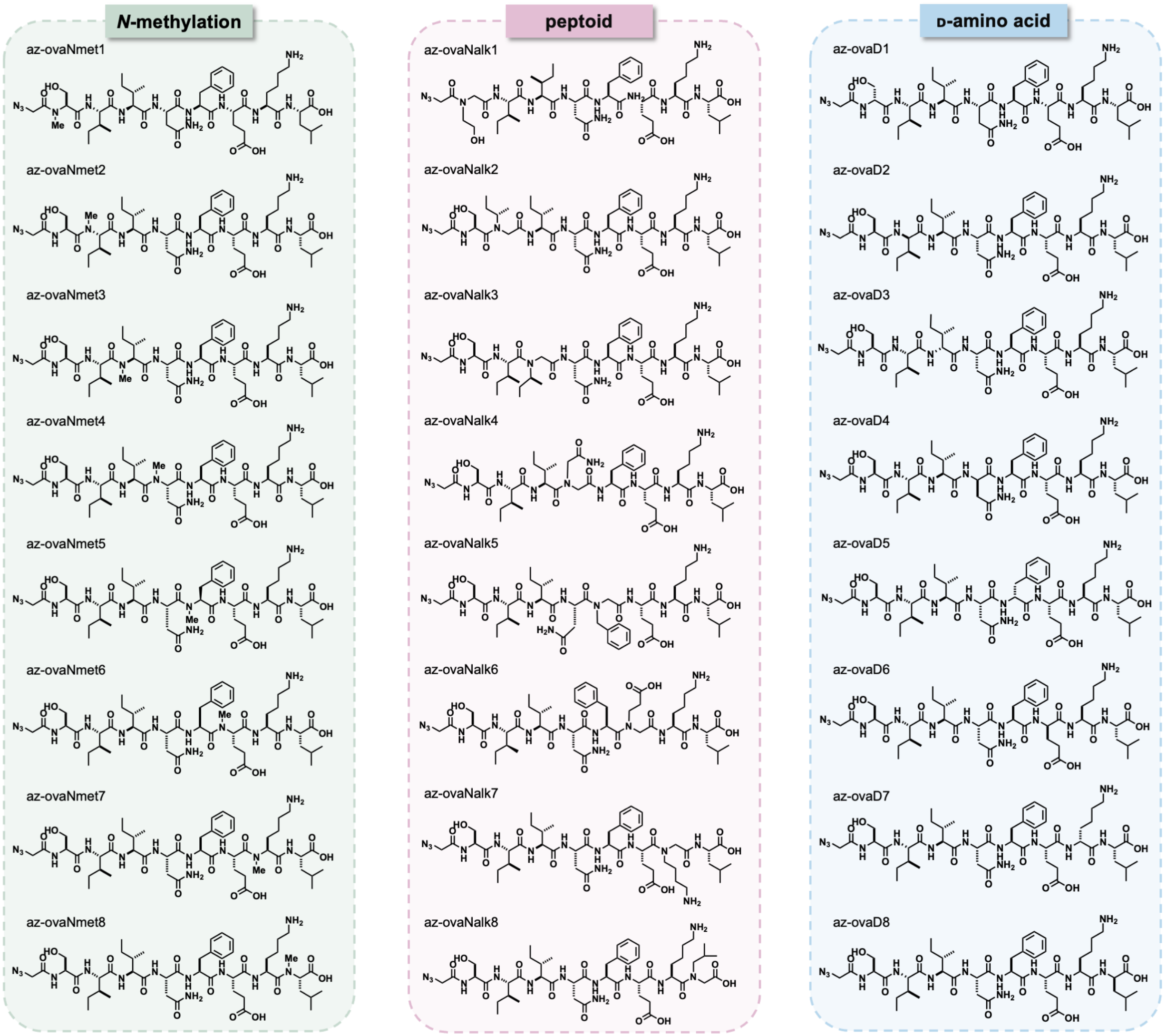


**Figure S5.** Chemical structures of azide-tagged peptidomimetic libraries.

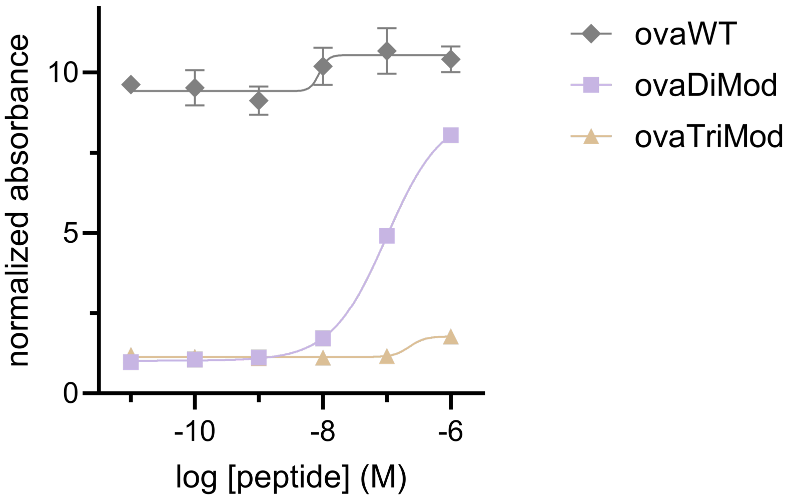


**Figure S6.** Dose-response curves from B3Z T cell activation assay for multiply-substituted variants **ovaDiMod** and **ovaTriMod** (full concentration scan from which the 100 nM data in **Fig. 8c** was obtained). RMA-S cells were incubated with the indicated concentrations of **ovaWT**, **ovaDiMod**, and **ovaTriMod** for 1 h at 26°C. RMA-S cells were subsequently co-incubated with B3Z T cells for 6 h at 37°C. β-galactosidase expression was then measured via the conversion of the colorimetric reagent chlorophenol red-β-galactopyranoside (CPRG) on a plate reader at 570 nm. The data presented has been normalized to the absorbance of the DMSO control. Data are represented as mean ± SD (*n* = 3), and Boltzmann sigmoidal curves were fitted to the data using GraphPad Prism.

### MATERIALS AND METHODS

#### Materials

| REAGENTS | VENDOR SOURCE | CATALOG # |
|---|---|---|
| **Reagents for biological methods** | | |
| RPMI 1640 Medium | Sigma-Aldrich | R8758 |
| DMEM, high glucose, pyruvate | Sigma-Aldrich | 11995065 |
| Fetal Bovine Serum | Sigma-Aldrich | A52568-01 |
| Penicillin-Streptomycin | Sigma-Aldrich | P4333 |
| Puromycin Dihydrochloride | Sigma-Aldrich | P4512 |
| TrypLE™ Express Enzyme (1X), phenol red | Fisher Scientific | 12605028 |
| APC anti-mouse H-2K$^{b}$ antibody | BioLegend | 116518 |
| TAMRA azide, 6-isomer (TMRaz) | Lumiprobe | D8130 |
| Formaldehyde solution | Sigma-Aldrich | 252549 |
| Saponin | Thermo Scientific | A18820.14 |
| Chlorophenol Red β-D-Galactopyranoside (CPRG) | Cayman Chemical | 29707 |
| 2-Mercaptoethanol | Sigma-Aldrich | M3148 |
| Magnesium chloride | Sigma-Aldrich | 208337 |
| Phosphate buffered saline | Sigma-Aldrich | P3813 |
| Mouse Serum | Novus Biologicals | S18193 |
| **Reagents for the synthesis and characterization of peptidomimetics** | | |
| Fmoc-L-leucine 4-alkoxybenzyl alcohol resin (100-200 mesh, 0.3-0.8 meq/g) | ChemImpex | 01911 |
| 2-Chlorotrityl chloride resin (1.0-2.0 meq/g, 100-200 mesh | Chem Impex | 03498 |
| α-*N*-Fmoc-amino acids | Chem Impex | Various |
| *N*-methylated amino acids | AmBeed | Various |
| *N*-alkyl substituted glycine building blocks | AK Scientific | Various |
| 2-Azidoacetic acid | AK Scientific | 1050AQ |
| N, N'-Diisopropylcarbodiimide (DIC) | Chem Impex | 00100 |
| Ethyl Cyano(hydroxyimino)acetate (Oxyma) | TCI Chemicals | E0847 |
| N, N-Dimethylformamide (DMF) | Sigma-Aldrich | 319937 |
| Dichloromethane (DCM) ACS-grade | Sigma-Aldrich | D65100 |
| Methanol ACS-grade | Sigma-Aldrich | 179337 |

| | | |
|---|---|---|
| Acetonitrile HPLC-grade | Sigma-Aldrich | 34851 |
| NMP | Sigma-Aldrich | 443778 |
| Piperidine | Sigma-Aldrich | 104094 |
| Trifluoroacetic acid (TFA) reagent-grade | ChemImpex | 00289 |
| Trifluoroacetic acid (TFA) HPLC-grade | Sigma-Aldrich | 302031 |
| Triisopropylsilane | ChemImpex | 01966 |
| **General materials and equipment** | | |
| Greiner Bio-One CELLSTAR TC Treated Cell Culture Flasks | Fisher Scientific | 07-000-225 |
| Corning™ Costar™ 96-Well, Cell Culture-Treated, Flat-Bottom Microplate | Fisher Scientific | 09-761-145 |
| Greiner Bio-One 96-well Non-treated Polystyrene Microplates | Fisher Scientific | 07-000-124 |
| MilliporeSigma™ Microcon-30 kDa Centrifugal Filter Unit with Ultracel-30 Membrane | Fisher Scientific | MRCF0R030 |

### Experimental Methods

#### Mammalian Cell Culture

RMA-S cells were a kind gift from Dr. John Sampson. B3Z cells were kindly provided by Dr. Aaron Esser-Kahn. HeLa cells expressing a GFP-HaloTag fusion protein (HT HeLa) were generously provided by the Chenoweth Lab at the University of Pennsylvania. RMA-S and B3Z cells were maintained in RPMI 1640 media supplemented with 10% fetal bovine serum, 50 IU/mL penicillin, and 50 μg/mL streptomycin. HT HeLa cells were cultured in Dulbecco's Modified Eagle Medium (DMEM) supplemented with 10% fetal bovine serum (FBS), 50 IU/mL penicillin, 50 μg/mL streptomycin, and 1 μg/mL puromycin. All cells were cultured in T75 flasks and maintained in a humidified atmosphere of 5% $CO_2$ at 37°C.

#### RMA-S Stabilization Assay

$10^5$ RMA-S cells were seeded in a treated 96 well plate at 37°C overnight. The next day, RMA-S cells were moved to a 26°C incubator for 24 hours. Following the incubation period, cells were incubated with peptides in culture media at indicated concentrations for 1 hour at 26°C before being moved to the 37°C incubator for 6 hours. Cells were removed from the well plate by vigorous pipetting and transferred to a round-bottom 96-well plate. Transferred cells were centrifuged (1100 x g, 5 min) in a Thermo Scientific Jouan C4i centrifuge and the supernatant was removed. Cell pellets were resuspended in a 1:100 dilution of APC-labeled anti-mouse H-2K$^b$ antibody in culture media for 1 hour at 4°C. Following centrifugation and removal of the supernatant, cells were then fixed with 4% formaldehyde solution, and analyzed using the Attune NxT Flow Cytometer (Thermo Fischer) equipped with a 637 nm laser with 670/14 nm bandpass filter.

#### B3Z T Cell Activation Assay

$10^5$ RMA-S cells were seeded in a treated 96 well plate at 37°C overnight. The next day, RMA-S cells were moved to a 26°C incubator for 24 hours. Following the incubation period, cells were incubated with peptides in culture media at indicated concentrations for 1 hour at 26°C. After this time, 1.5 x $10^4$ B3Z cells in culture media were co-incubated with the RMA-S cells for 6 hours. Cells were centrifuged (1100 x g, 5 min) in a Thermo Scientific Jouan C4i centrifuge and the supernatant was removed. Lysis buffer containing 0.2% saponin, 500 mM CPRG reagent (Chlorophenol Red β-D-Galactopyranoside), 20 mM $MgCl_2$, and 100 mM β-mercaptoethanol in 1X PBS was added to each well. After 1 hour, absorbance at 570 nm was recorded using a BioTek Synergy H1 Microplate Reader.

**Chloroalkane HaloTag Azide-based Membrane Penetration (CHAMP Assay)**

5 x $10^4$ HT HeLa cells were seeded in a treated 96 well plate at 37°C overnight, reaching 80 – 90% confluency. The next day, cells were washed three times with 1X PBS and treated with 10 μM DBCOcl for 15 minutes. Cells underwent a "pulse" step with the azide-tagged peptidomimetics at 50 μM for 24 hours. After "pulse" time, cells were "chased" with 50 μM azido-tetramethylrhodamine (TMRaz) for 15 minutes. Cells were washed three times with 1X PBS between each step. After dye treatment, cells were removed using TrypLE™ Express Enzyme and transferred to a round-bottom 96-well plate. Transferred cells were centrifuged (1100 x g, 5 min) in a Thermo Scientific Jouan C4i centrifuge, and the cell pellets were resuspended and fixed in 4% formaldehyde solution for 20 minutes. Cells were analyzed using an Attune NxT Flow Cytometer (Thermo Fisher) equipped with a 561 nm laser with 585/15 nm bandpass filter.

**Serum Stability Assay**

Time course degradation in mouse serum was evaluated for peptides **ovaWT**, **ovaDiMod**, and **ovaTriMod**. Compounds were incubated in mouse serum at a concentration of 5.0 mM to a total of 400 μL at 37 °C. At each time point (0, 15, 30, 60 min), 80 μL of the solution was aliquoted and diluted in a 1:1 solution of $H_2O/CH_3CN$ (202 μL) containing trichloroacetic acid (18 μL) and 1 mM of internal standard Fmoc-ʟ-phenylalanine. This mixture was incubated in ice for 5 min to precipitate serum proteins. The samples were then centrifuged at 21,000 x g for 5 min. The supernatant was then transferred to a filter tube and centrifuged again at 21,000 x g for 5 min to filter. 50 μL of filtrate was then analyzed using Phenomenex Luna 5 μm C8(2) on RP-HPLC; gradient elution in $H_2O/CH_3CN$ with 0.1% TFA at 1 mL/min. Peptide identity was confirmed via peak collection and subsequent matrix-assisted laser desorption ionization time-of-flight (MALDI-TOF) mass spectroscopy (Shimadzu 8020). To eliminate variations in the overall signal intensity, area under the curve (AUC) for the peptides of interest at each time point was normalized to the AUC of the internal standard Fmoc-ʟ-phenylalanine, the concentration of which was predetermined. These normalized AUC values were then further normalized to the "0 min" time point which was set to 100%. The percentage of peptide remaining was then plotted as a function of time.

### SYNTHESIS AND CHARACTERIZATION OF PEPTIDOMIMETICS

#### General procedure for the solid-phase synthesis of peptidomimetics

All peptides were prepared by standard Fmoc-based solid-phase chemistry using the appropriate resins. Briefly, to a 25 mL peptide synthesis vessel, an appropriate amount of resin was added, followed by 20% piperidine in N, N-Dimethylformamide (DMF, 15 mL) (if resin was Fmoc-protected). This was followed by shaking at room temperature for 30 min. The resin was then washed with methanol ($CH_3OH$) and dichloromethane (DCM) three times. After the last wash, 4 equiv. of amino acid was added along with 4 equiv. of ethyl cyanohydroxyiminoacetate (Oxyma) and 4 equiv. of N, N'-Diisopropylcarbodiimide (DIC). The resin was shaken at room temperature for 2 h, then washed with $CH_3OH$/DCM. The remainder of the amino acids were coupled in the same manner. Peptides were cleaved from the resin using a TFA/TIPS/$H_2O$ mixture (95:2.5:2.5, v/v/v) shaking at room temperature for 2 h. The solution was filtered and concentrated prior to precipitation by the addition of cold diethyl ether to yield crude peptide. Crude peptides were purified by reverse-phased preparative high-performance liquid chromatography (RP-HPLC) equipped with Waters 1525 with 2489 UV/Visible Detector on a Phenomenex Luna 10 μm C8(2) 100 Å (250 x 21.2 mm) column using gradient elution with $H_2O$/$CH_3OH$ with 0.1% TFA at 10 mL/min. The HPLC fractions of the desired purified compounds were first concentrated under reduced pressure using a rotary evaporator. The final concentrated aqueous solutions were lyophilized to dryness using Labconco Freezone 4.5 L (-84°C) lyophilizer. The peptides were analyzed for purity using Phenomenex Luna 5 μm C8(2) on the same RP-HPLC; gradient elution in $H_2O$/$CH_3OH$ with 0.1% TFA at 1 mL/min. Peptide identities were analyzed via matrix-assisted laser desorption ionization time-of-flight (MALDI-TOF) mass spectrometry (Shimadzu 8020).

#### Solid-phase peptide synthesis of *N*-methylation and D-amino acid libraries

To prepare peptides in this library, Fmoc-L-leucine 4-alkoxybenzyl alcohol resin (0.608 mmol/g of loading capacity) was used such that the *C*-terminus would remain as a carboxylic acid upon cleavage. For **ovaNmet8** and **ovaD8** (and their corresponding *N*-terminal azide peptides), 2-Chlorotrityl chloride resin (1.42 mmol/g of loading capacity) was used instead, since the modifications were located at the leucine residue, but would still allow the *C*-terminus to remain as a carboxylic acid upon cleavage. Synthesis was performed in accordance with established procedures for the resin. Commercially available *N*-methyl amino acids and D-amino acids were used for the *N*-methyl and D-amino acid library analogs, respectively. For the corresponding *N*-terminal azide series of these libraries, 2-azido-acetic acid was coupled on the *N*-terminus of each peptide on resin. Cleavage and purification were carried out as described before. All peptides in this library were characterized using UV-Vis absorbance of the phenylalanine residue in their sequence at 257 nm ($\varepsilon$ = 195 $cm^{-1}M^{-1}$).

**Solid-phase peptoid synthesis of *N*-alkylation library**

As before, Fmoc-ʟ-leucine 4-alkoxybenzyl alcohol resin (0.608 mmol/g of loading capacity) was used such that the *C*-terminus would remain as a carboxylic acid upon cleavage. For **ovaNalk8** (and its corresponding *N*-terminal azide peptidomimetic), 2-Chlorotrityl chloride resin (1.42 mmol/g of loading capacity) was used instead. For the peptoid library, the *N*-alkyl substituted glycine residues in the peptoids were added as per reported protocols.[1] Briefly, a 25 mL peptide synthesis vessel containing the resin, conjugated with or without peptide residues, was washed with $CH_3OH$/DCM, following Fmoc deprotection of the most recently added amino acid using 20% piperidine in DMF as described above. After the last wash, 0.8 M 2-bromo acetic acid was diluted in 5 mL of DMF, followed by addition of 0.8 M DIC, and this solution was added to the resin. The resin was shaken at room temperature for 20 min, then washed with DMF 5 times. 1.5 M primary amine building blocks were then diluted in 5 mL N-methyl-2-pyrrolidone (NMP), and this solution was added to the resin and shaken at room temperature for 2 hours.* The rest of the amino acids were coupled by standard Fmoc-based solid-phase chemistry according to the sequence of the peptide. As before, for the corresponding *N*-terminal azide series of this library, 2-azido-acetic acid was coupled on the *N*-terminus of each peptoid on resin. Cleavage and purification were carried out as described before. All peptoids in this library were characterized using UV-Vis absorbance of the phenylalanine residue in their sequence at 257 nm ($\varepsilon$ = 195 $cm^{-1}M^{-1}$).

***** *For primary amine building blocks that were commercially available as HCl salts, a free basing protocol was followed. For this, a 1 M solution of the primary amine building block was made in 12.5 mL DMF. Separately, 0.36 g of KOH was dissolved in 0.72 mL of water. The solution of KOH was quickly added to the 1 M amine solution, vortexed, and centrifuged at 4,000 x g for 10 min. The supernatant was then added to the vessel for the displacement reaction and allowed to shake at room temperature for 2 hours.*

### Characterization of Peptidomimetics

The peptidomimetics were analyzed for purity using Phenomenex Luna 5 μm C8(2) on the same RP-HPLC; gradient elution in $H_2O/CH_3CN$ with 0.1% TFA at 1 mL/min. Peptides were analyzed via matrix-assisted laser desorption ionization time-of-flight (MALDI-TOF) mass spectrometry (Shimadzu 8020). All peptides exhibited a purity greater than 93%. All peptidomimetics were characterized using UV-Vis absorbance of the phenylalanine residue in their sequence at 257 nm ($\varepsilon$ = 195 $cm^{-1}M^{-1}$). Characterization data are provided below.

**ovaWT**

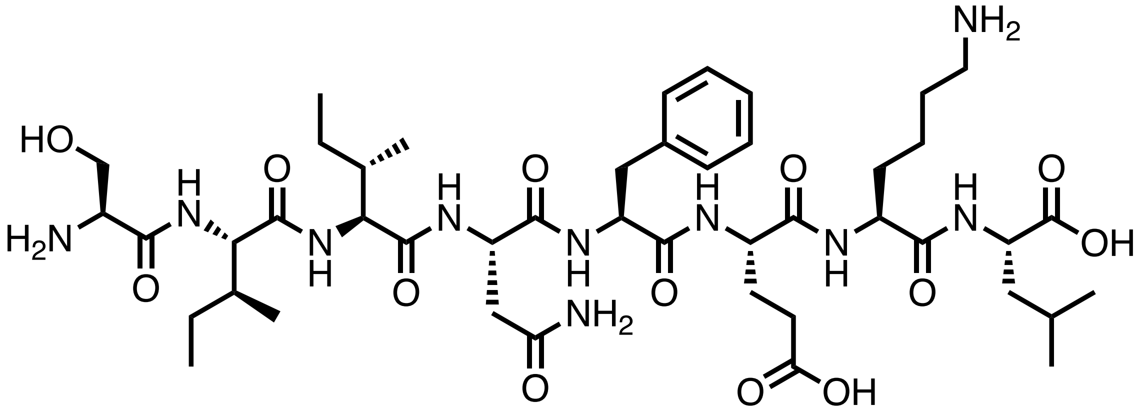


Chemical structure of **ovaWT**.

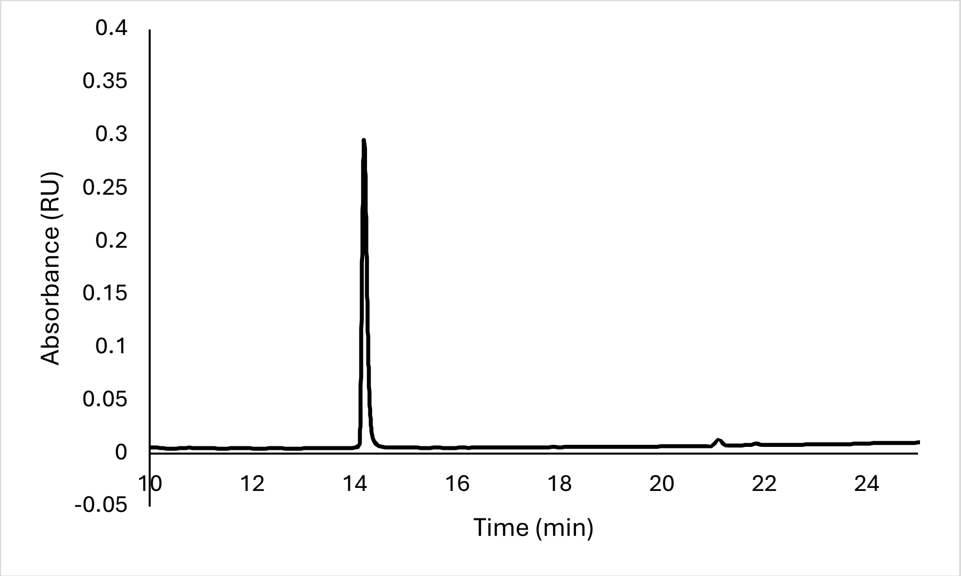


Analytical HPLC Chromatogram of **ovaWT**.

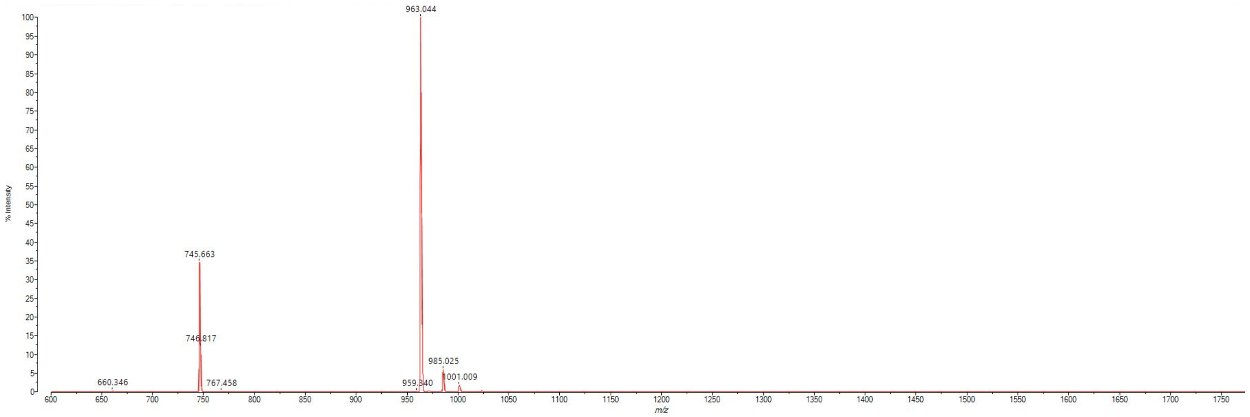


MALDI-TOF mass spectrum for **ovaWT** (m/z 963.551 for $[M+H]^+$, found 963.044).

**ovaNmet1**

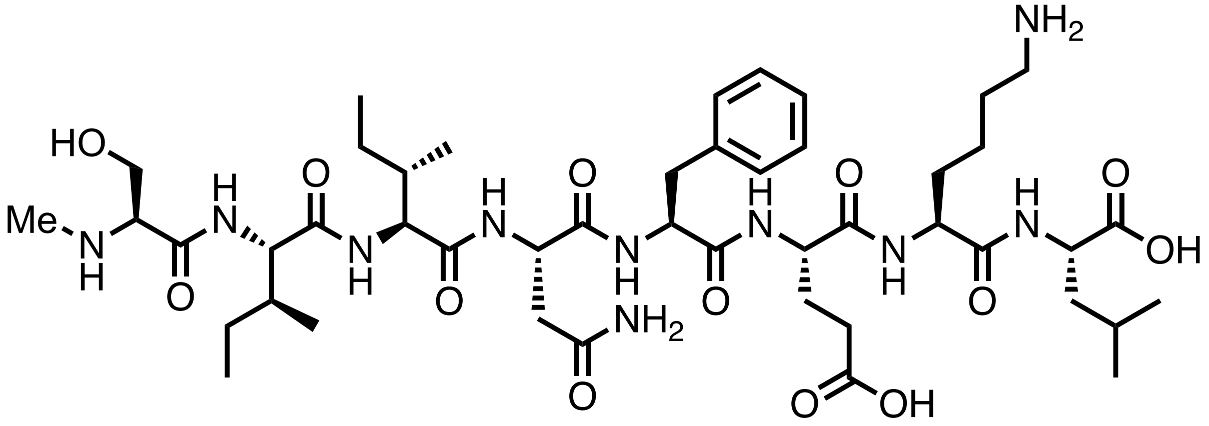

Chemical structure of **ovaNmet1**.

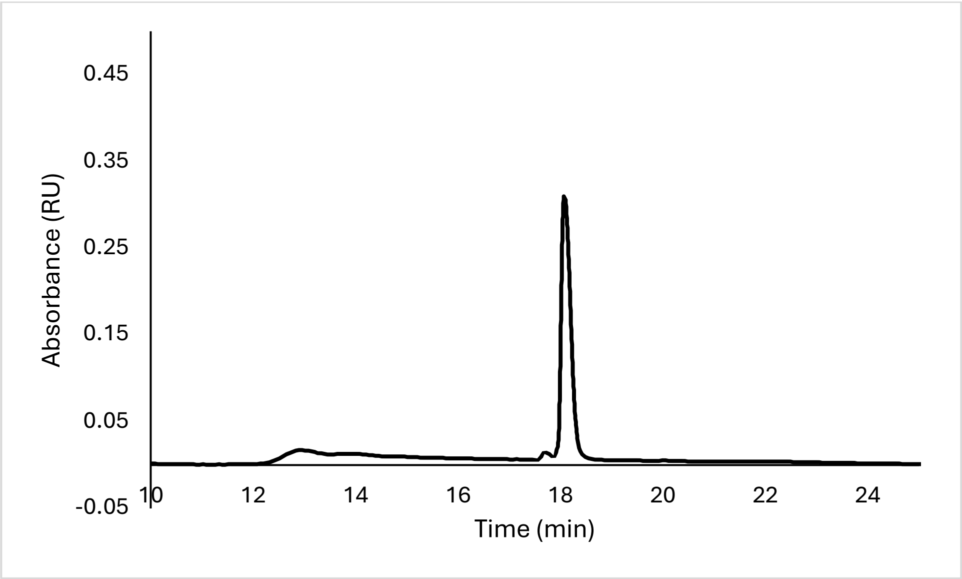


Analytical HPLC Chromatogram of **ovaNmet1**.

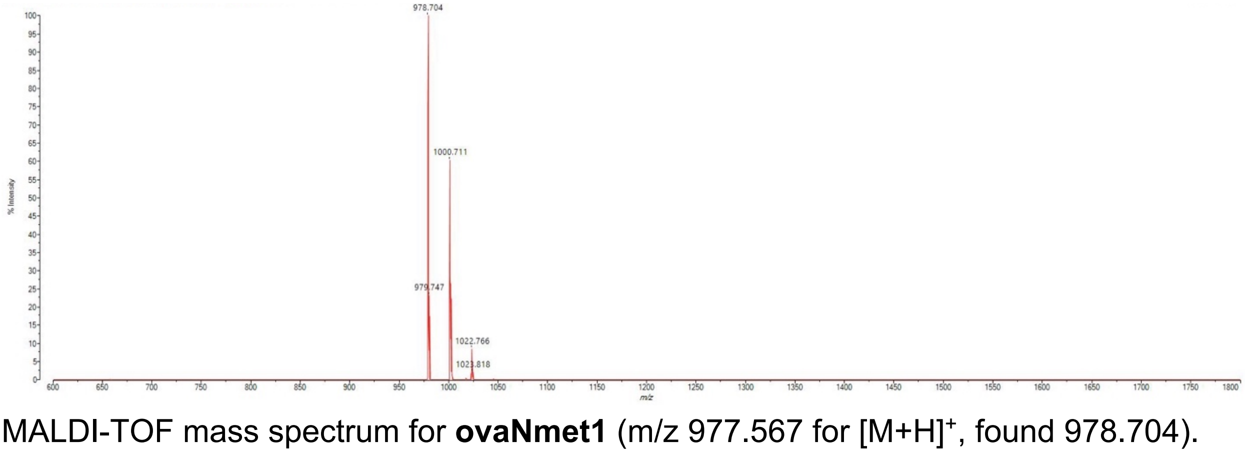


MALDI-TOF mass spectrum for **ovaNmet1** (m/z 977.567 for $[M+H]^+$, found 978.704).

**ovaNmet2**

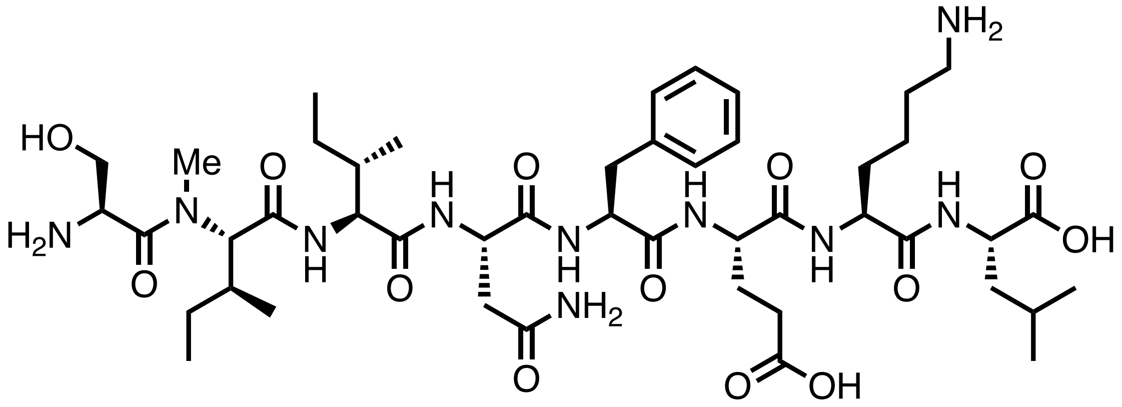


Chemical structure of **ovaNmet2**.

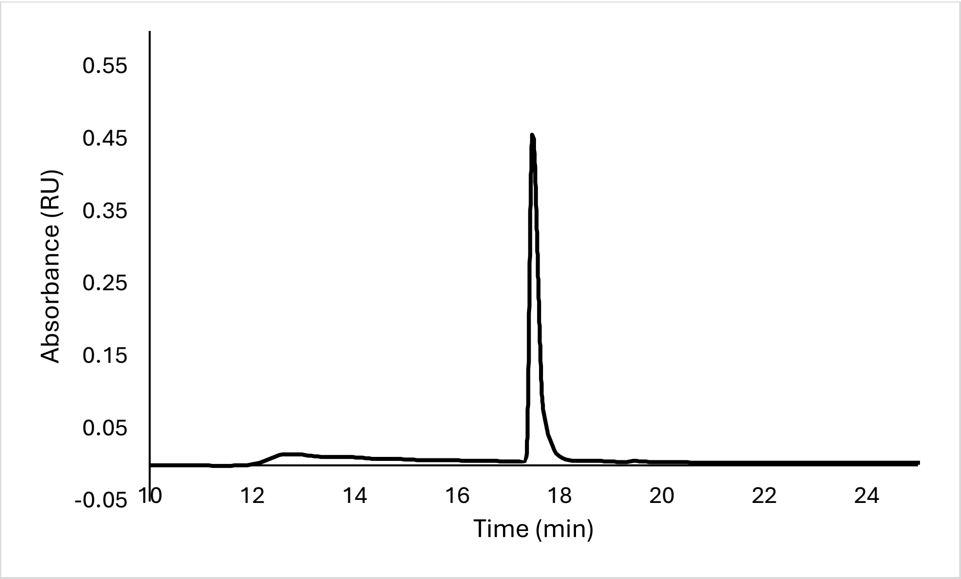


Analytical HPLC Chromatogram of **ovaNmet2**.

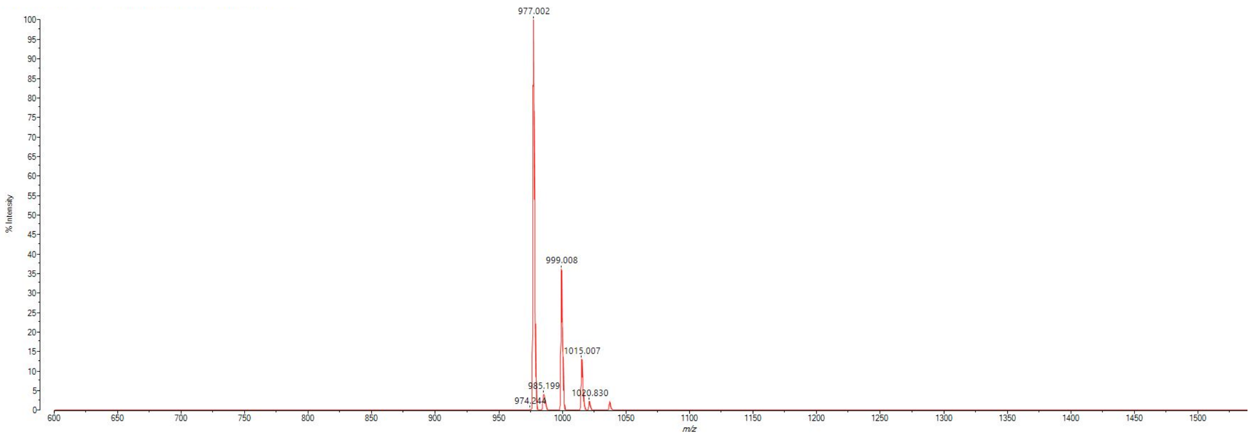


MALDI-TOF mass spectrum for **ovaNmet2** (m/z 977.567 for $[M+H]^+$, found 977.002).

**ovaNmet3**

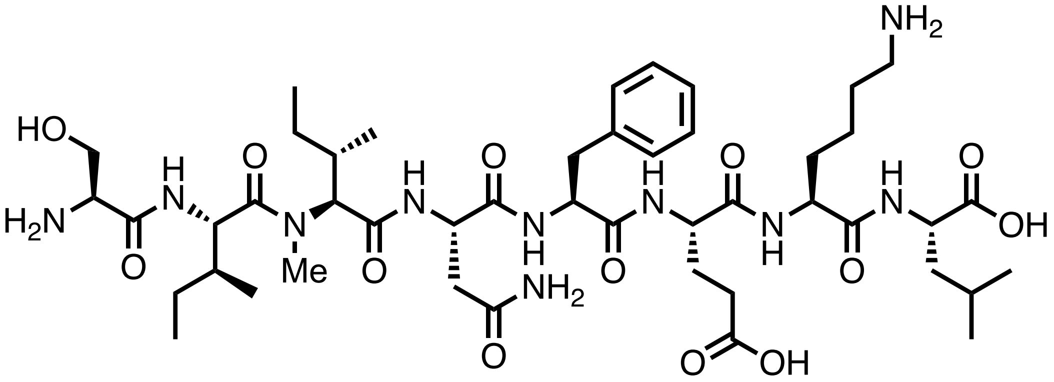


Chemical structure of **ovaNmet3**.

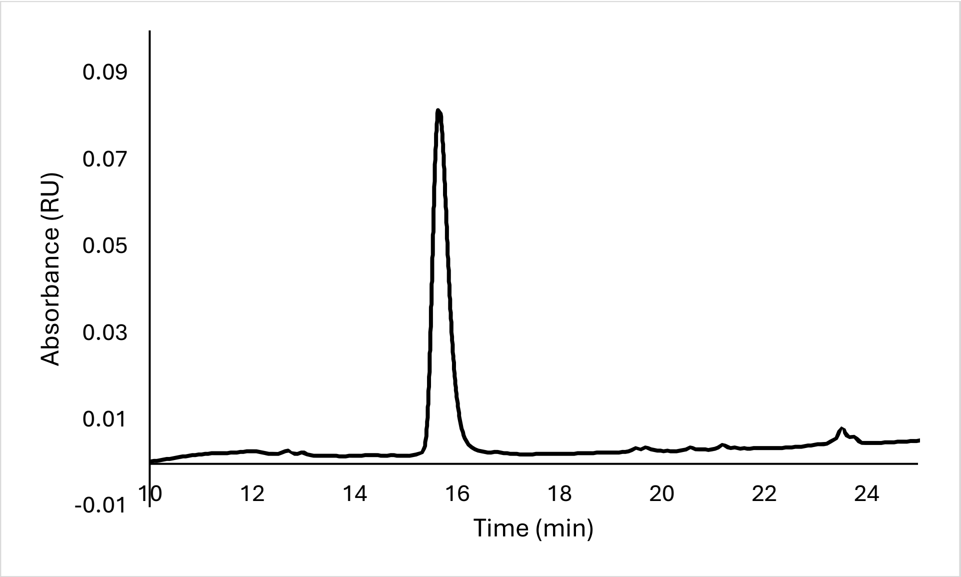


Analytical HPLC Chromatogram of **ovaNmet3**.

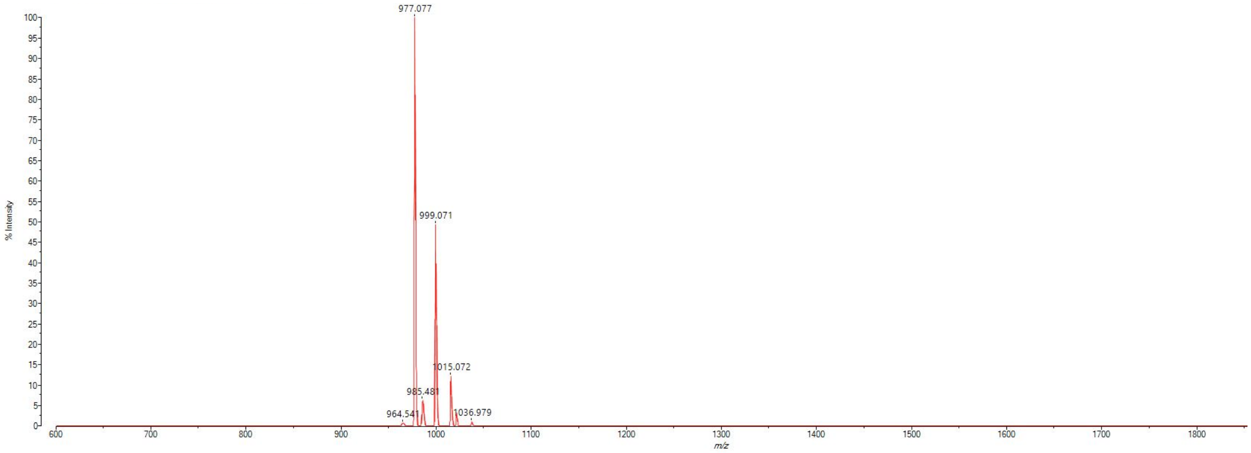


MALDI-TOF mass spectrum for **ovaNmet3** (m/z 977.567 for $[M+H]^+$, found 977.077)

**ovaNmet4**

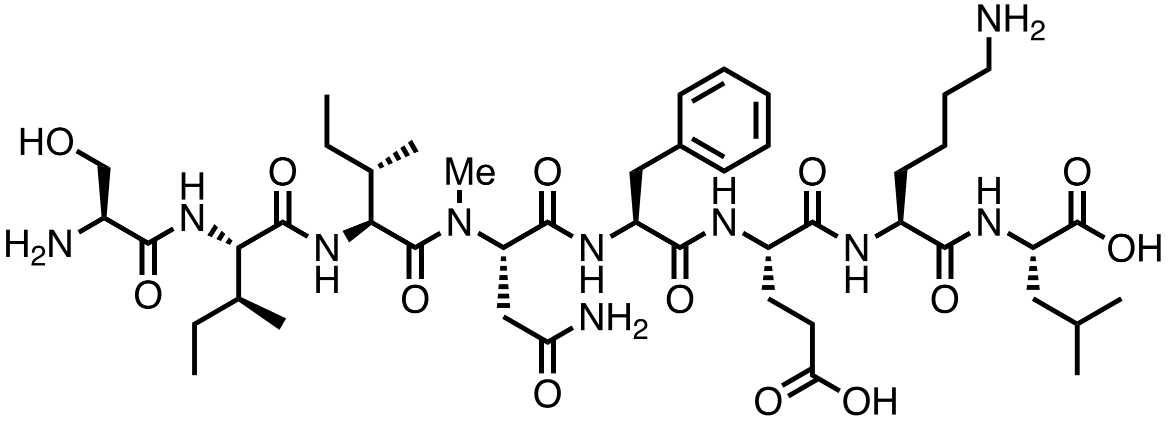

Chemical structure of **ovaNmet4**.

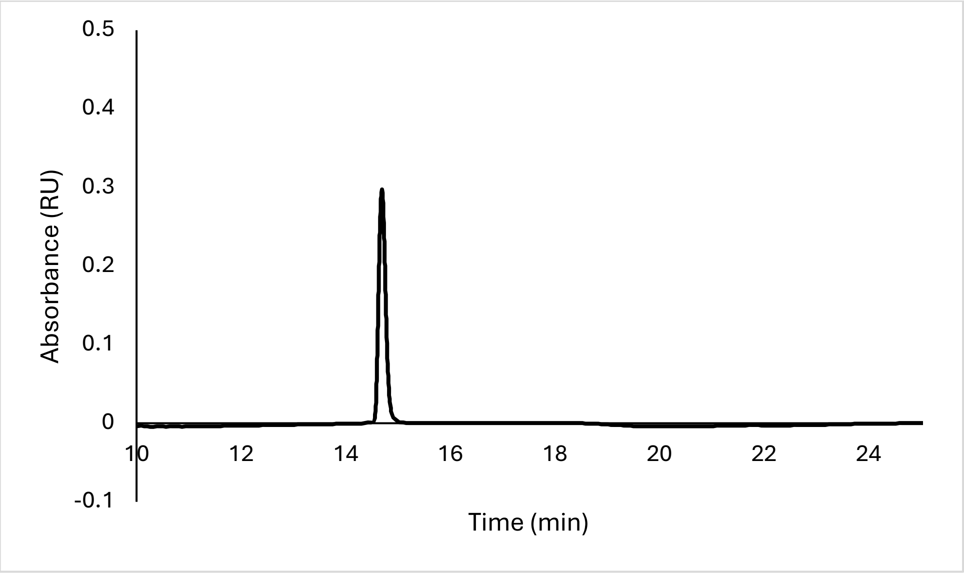


Analytical HPLC Chromatogram of **ovaNmet4**.

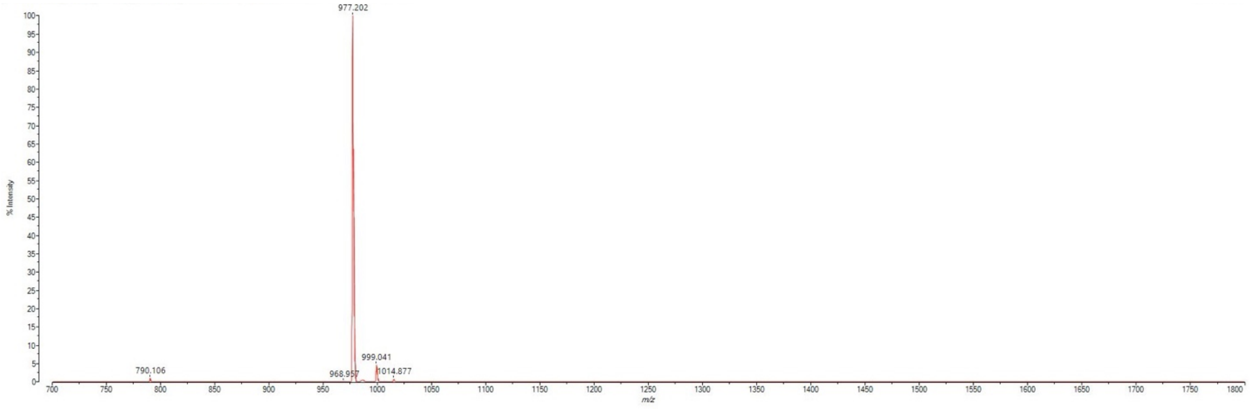


MALDI-TOF mass spectrum for **ovaNmet4** (m/z 977.567 for $[M+H]^+$, found 977.202).

**ovaNmet5**

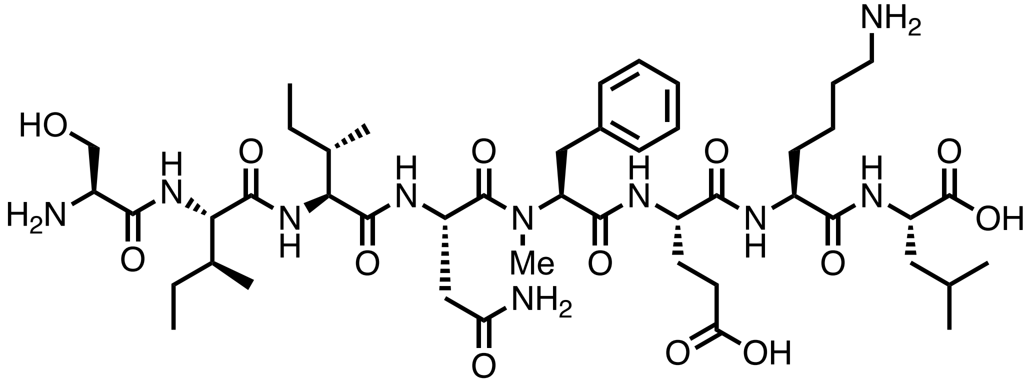


Chemical structure of **ovaNmet5**.

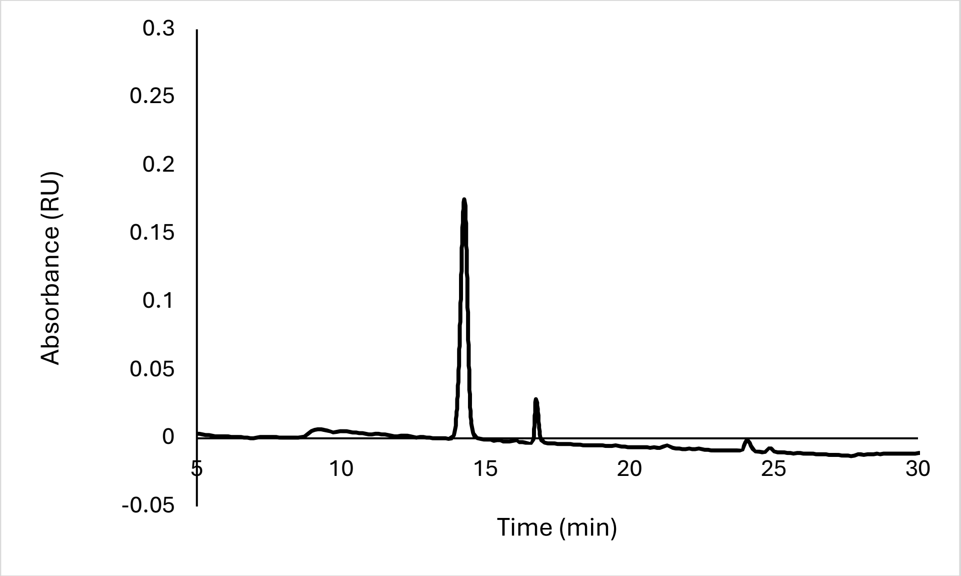


Analytical HPLC Chromatogram of **ovaNmet5**.

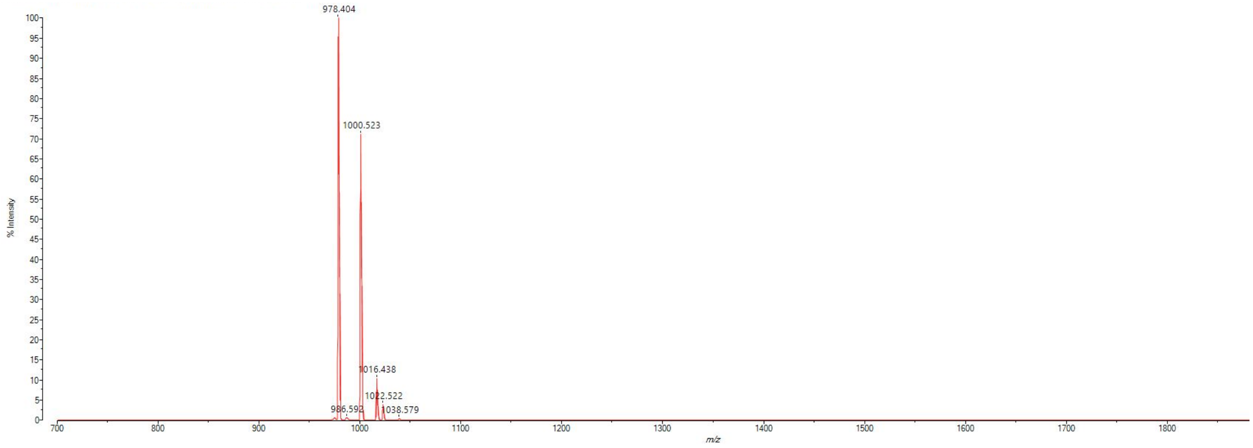


MALDI-TOF mass spectrum for **ovaNmet5** (m/z 977.567 for $[M+H]^+$, found 978.404).

**ovaNmet6**

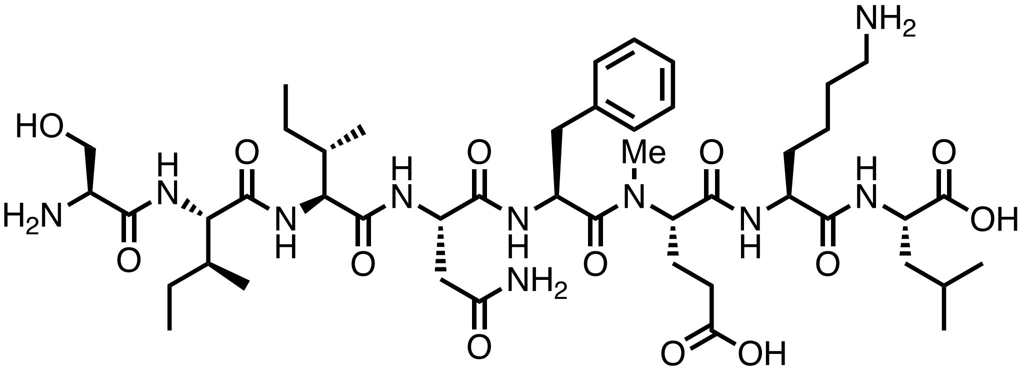

Chemical structure of **ovaNmet6**.

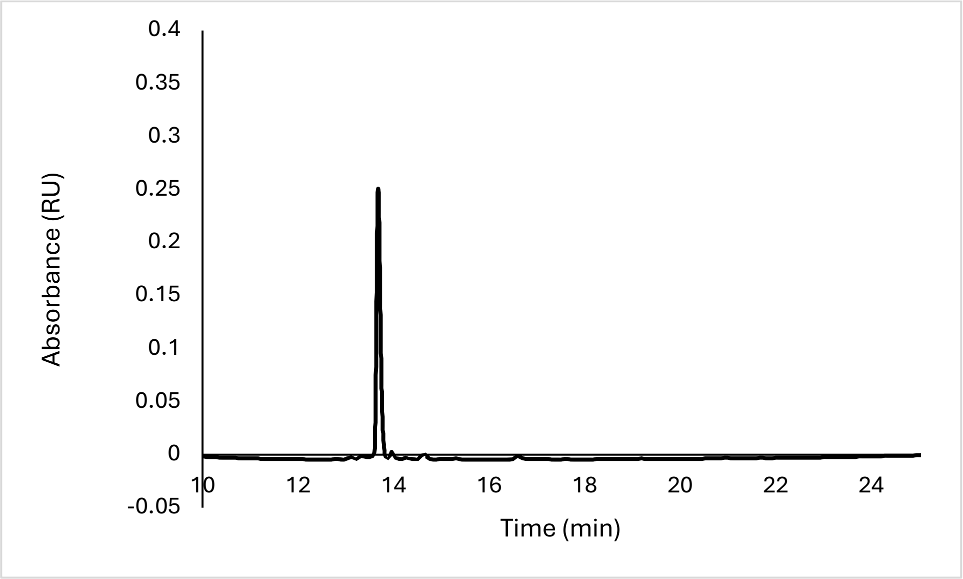


Analytical HPLC Chromatogram of **ovaNmet6**.

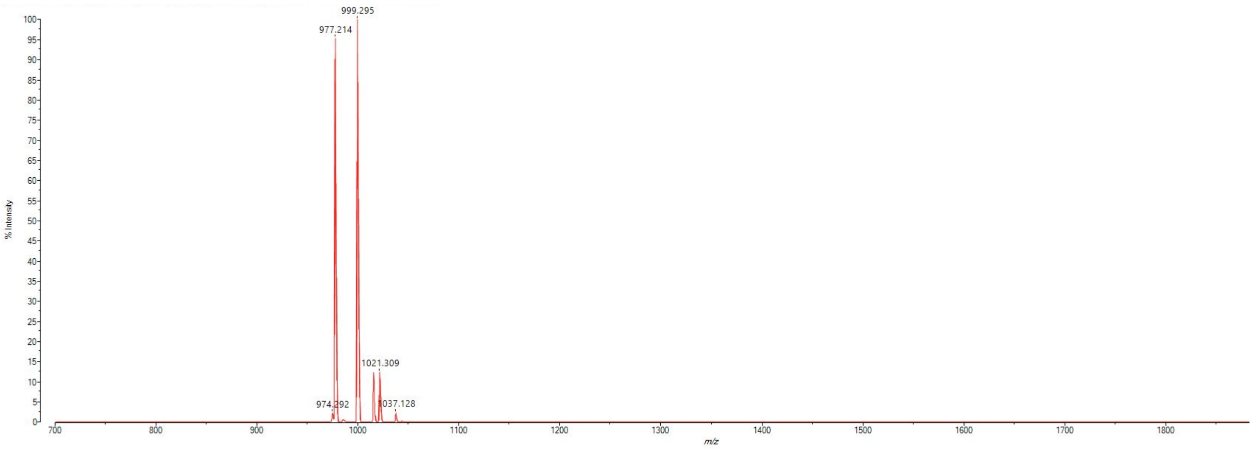


MALDI-TOF mass spectrum for **ovaNmet6** (m/z 977.567 for $[M+H]^+$, found 977.214).

**ovaNmet7**

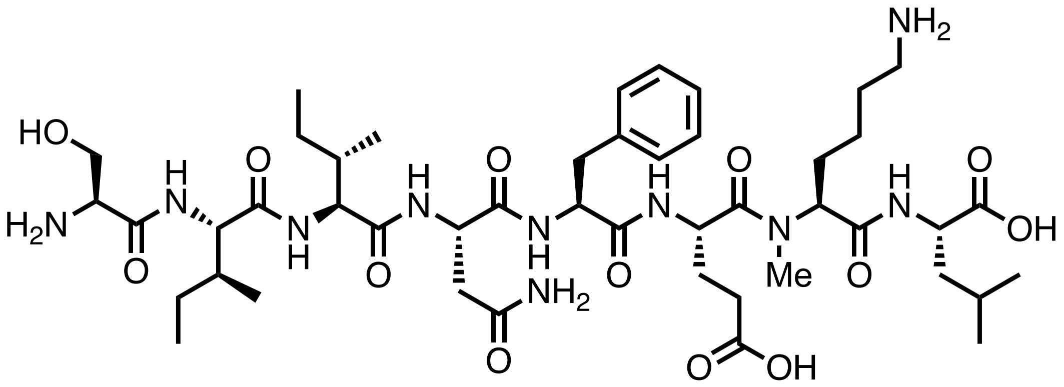


Chemical structure of **ovaNmet7**.

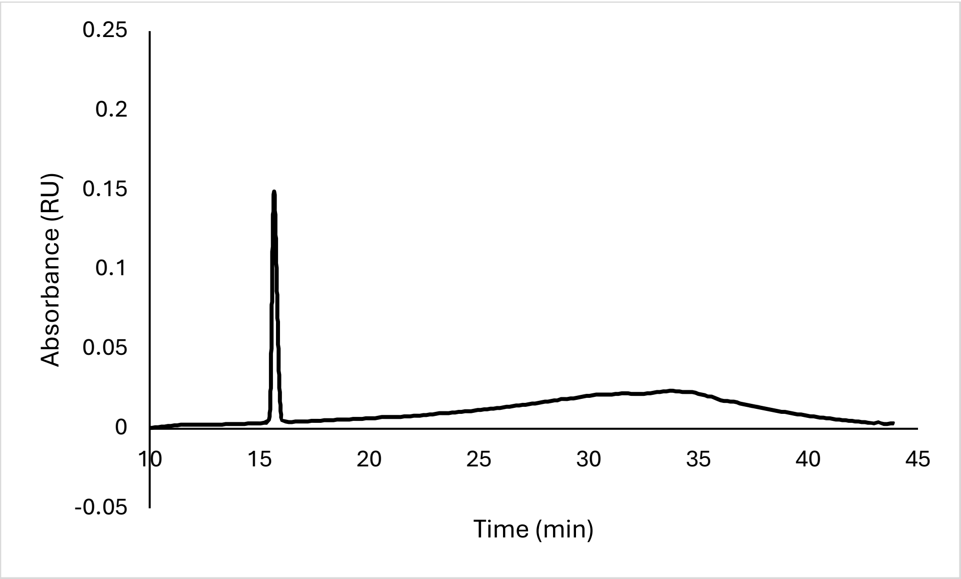


Analytical HPLC Chromatogram of **ovaNmet7**.

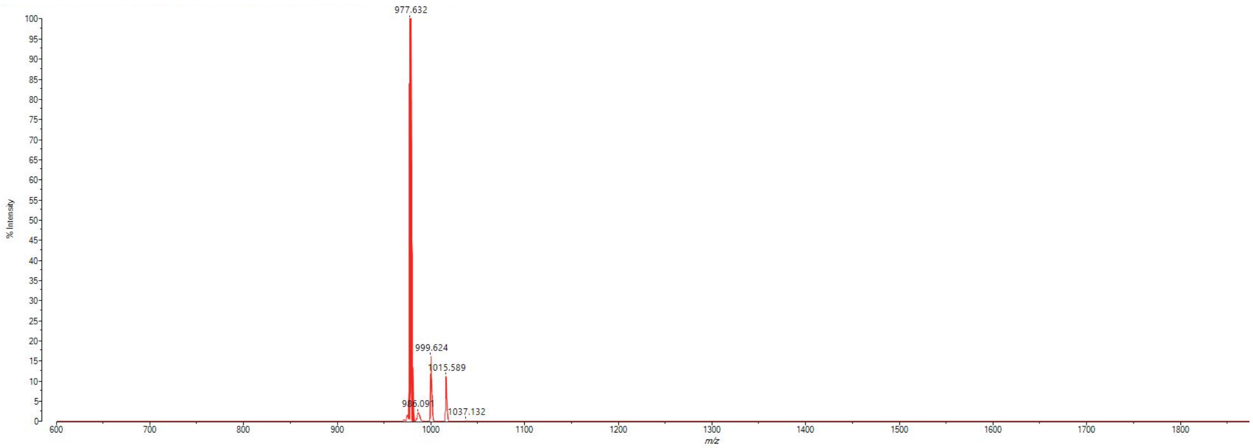


MALDI-TOF mass spectrum for **ovaNmet7** (m/z 977.567 for $[M+H]^+$, found 977.632).

**ovaNmet8**

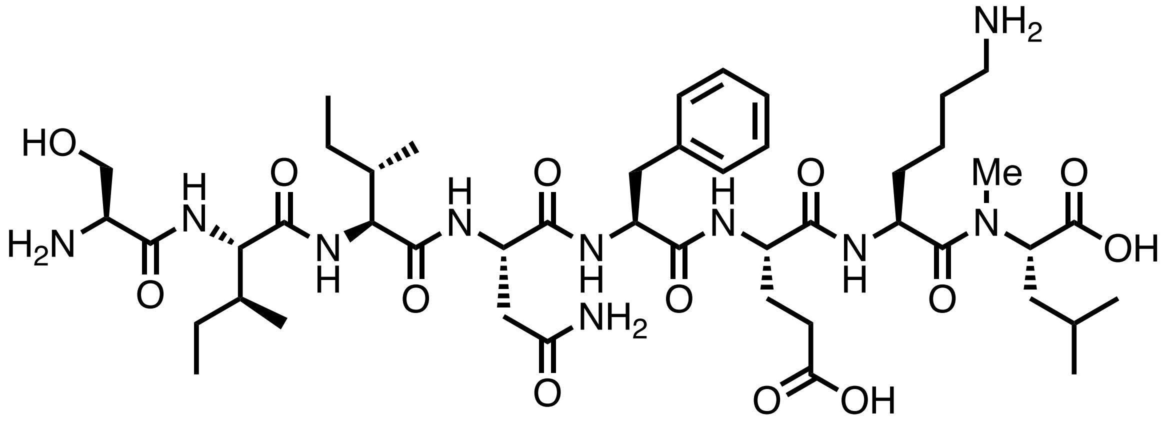


Chemical structure of **ovaNmet8**.

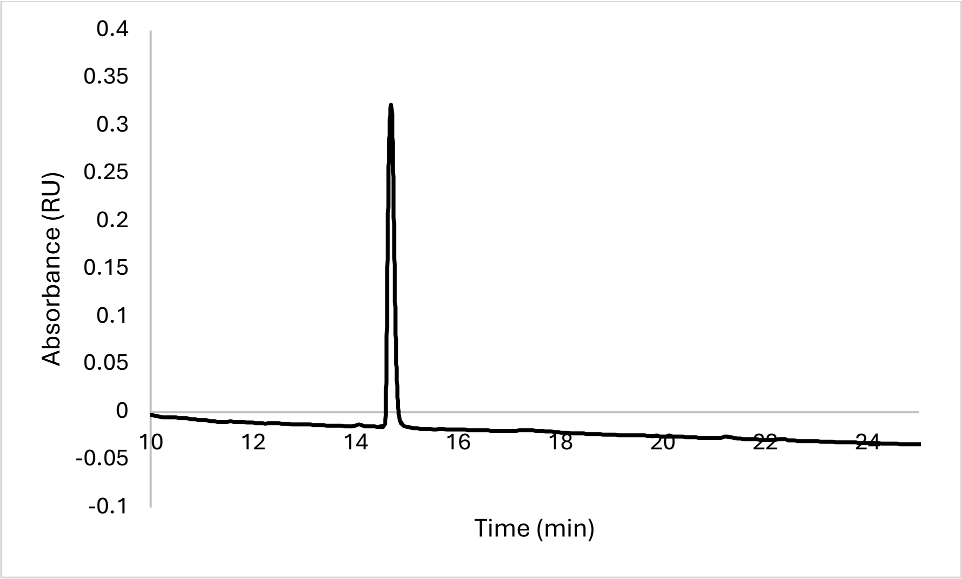


Analytical HPLC Chromatogram of **ovaNmet8**.

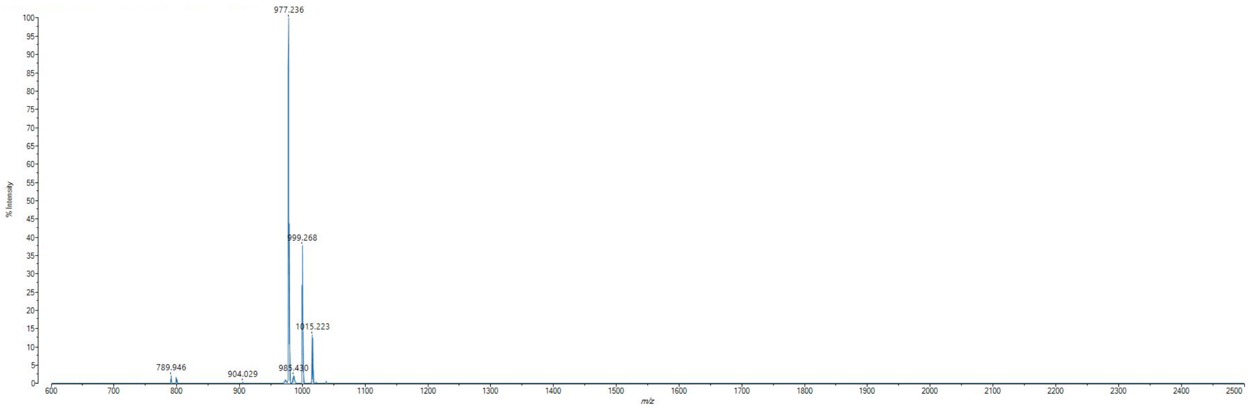


MALDI-TOF mass spectrum for **ovaNmet8** (m/z 977.567 for $[M+H]^+$, found 977.236).

**ovaNalk1**

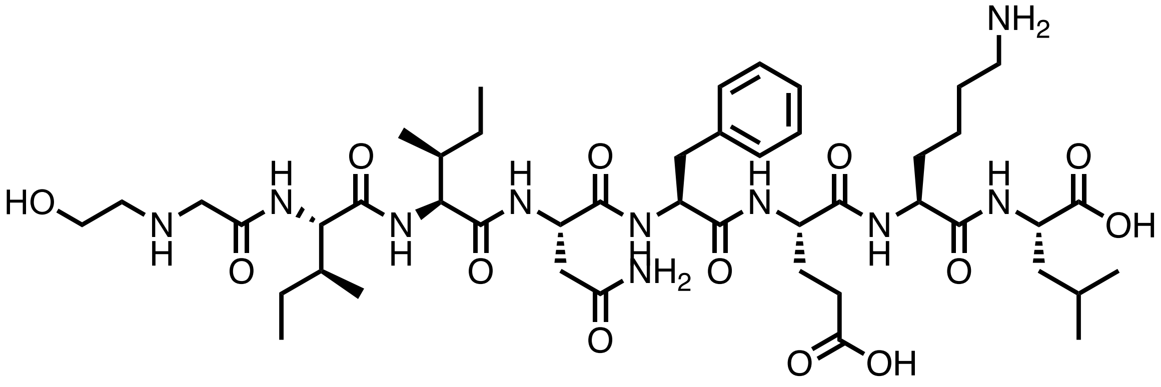

Chemical structure of **ovaNalk1**.

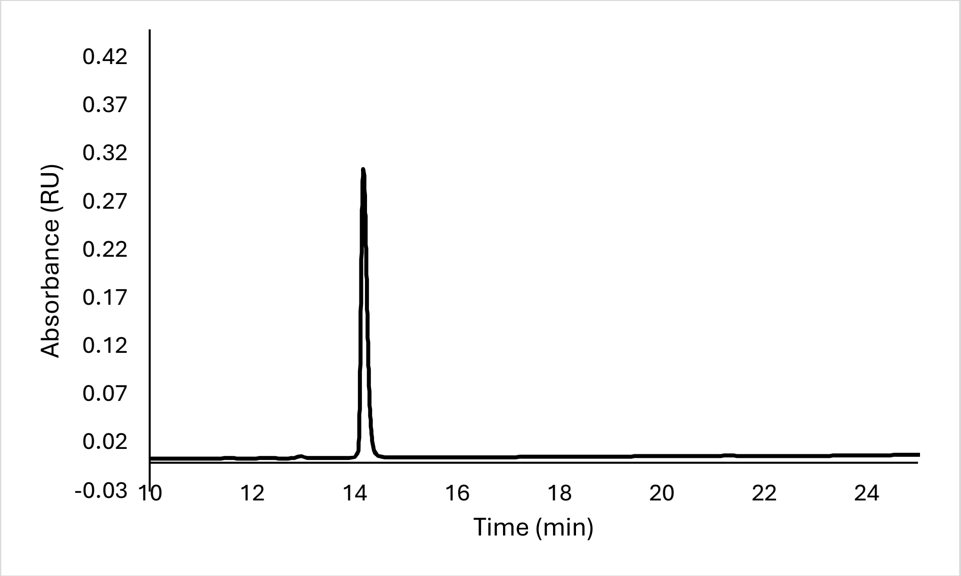


Analytical HPLC Chromatogram of **ovaNalk1**.

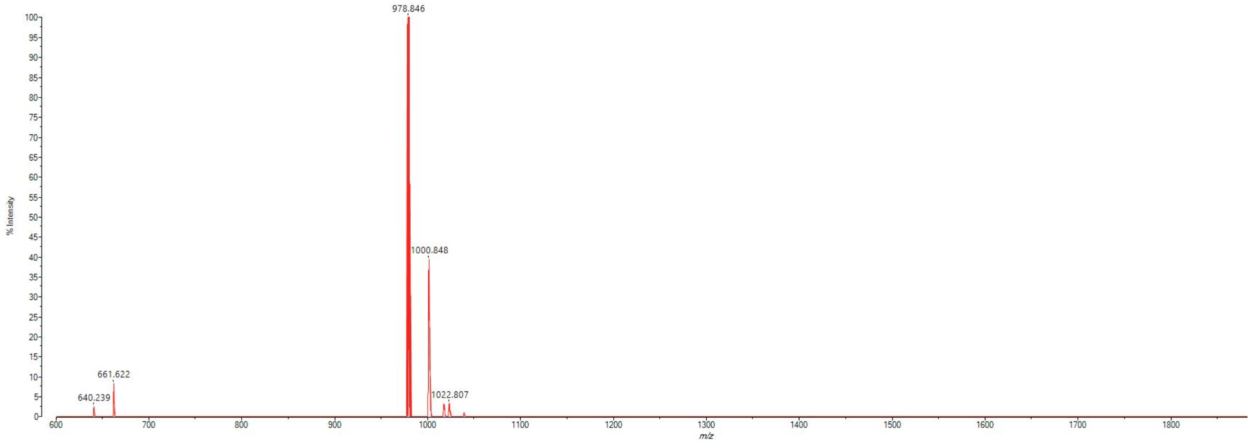


MALDI-TOF mass spectrum for **ovaNalk1** (m/z 977.567 for $[M+H]^+$, found 978.846).

**ovaNalk2**

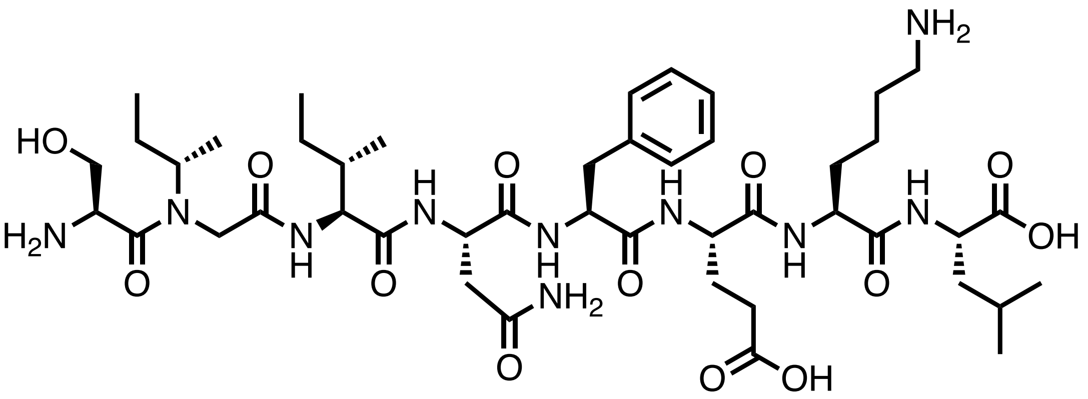

Chemical structure of **ovaNalk2**.

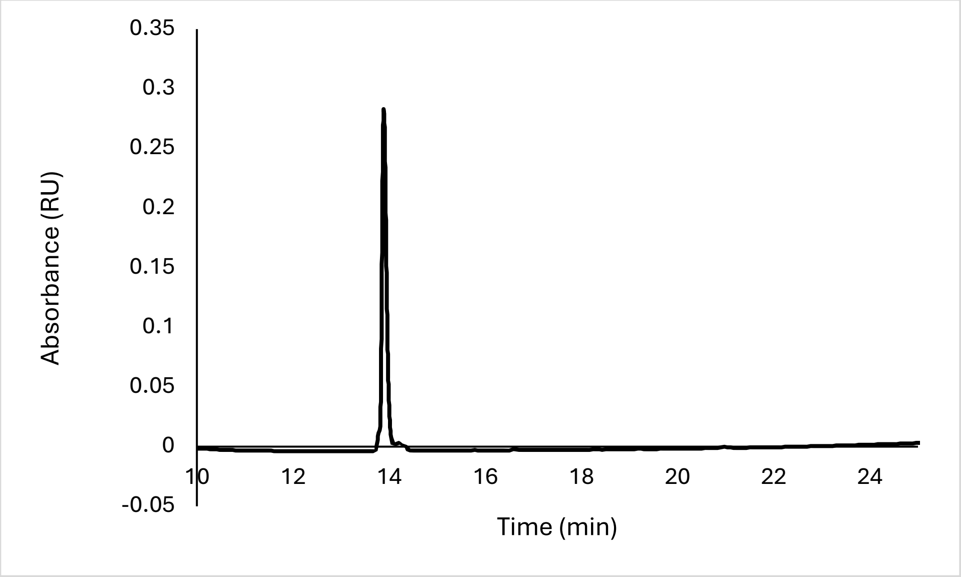


Analytical HPLC Chromatogram of **ovaNalk2**.

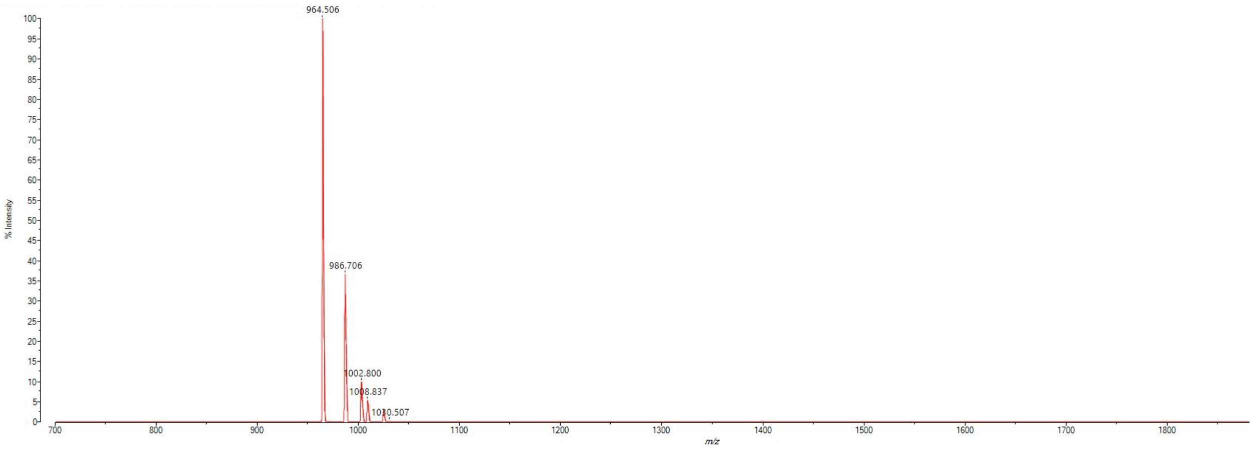


MALDI-TOF mass spectrum for **ovaNalk2** (m/z 963.551 for $[M+H]^+$, found 964.506).

**ovaNalk3**

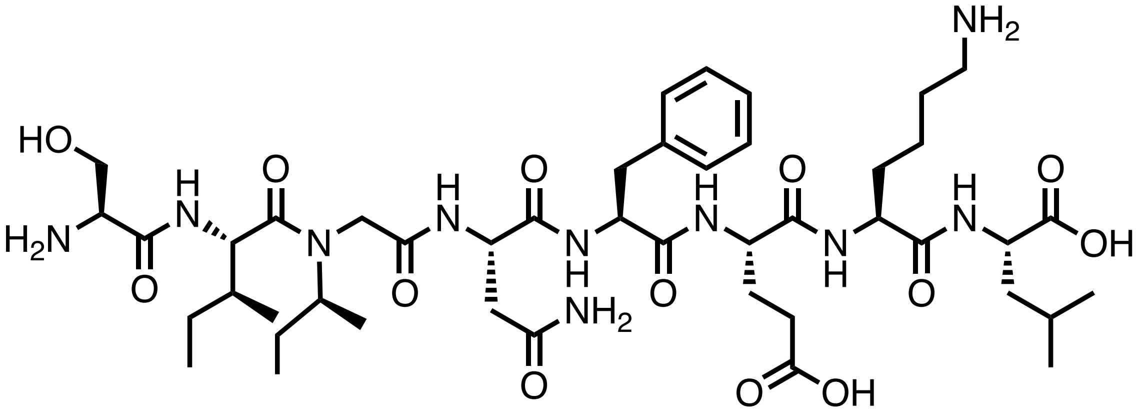

Chemical structure of **ovaNalk3**.

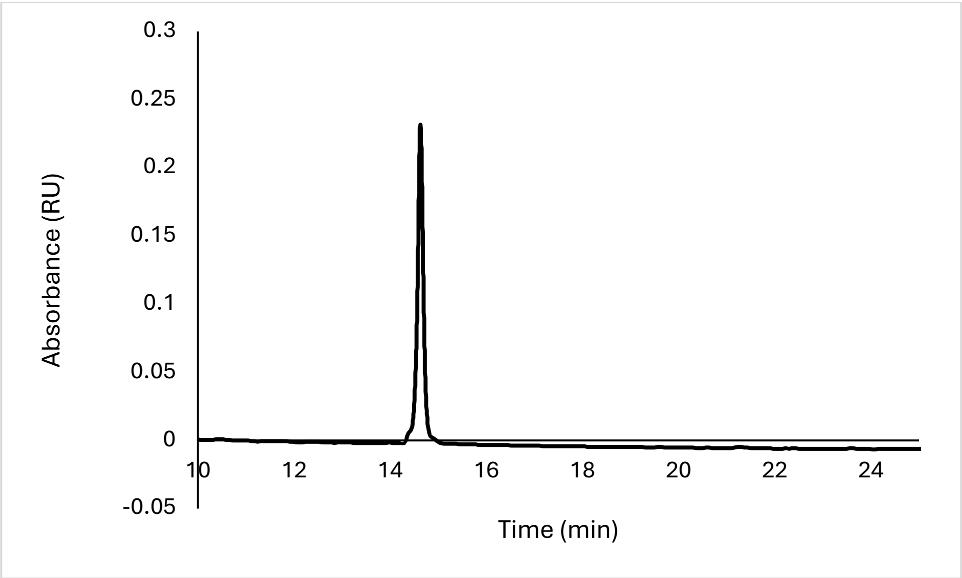


Analytical HPLC Chromatogram of **ovaNalk3**.

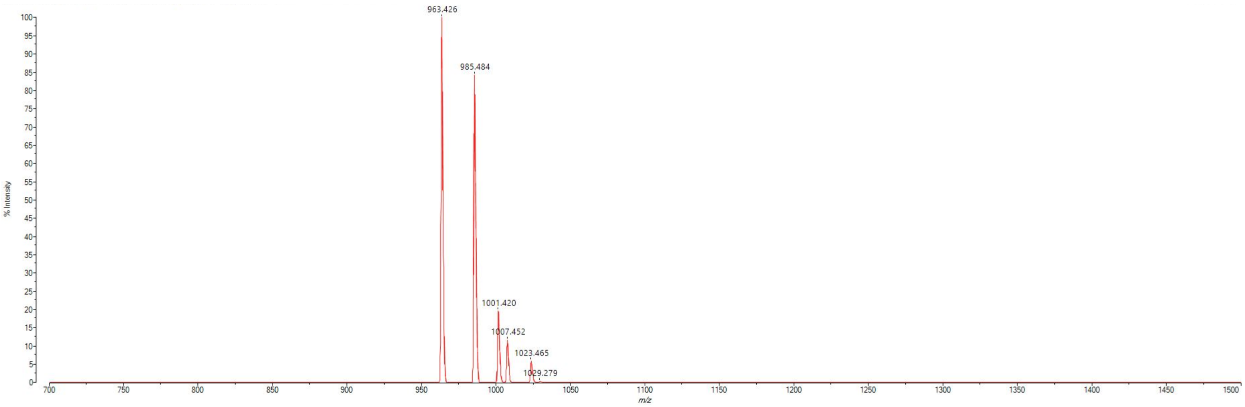


MALDI-TOF mass spectrum for **ovaNalk3** (m/z 963.551 for $[M+H]^+$, found 963.426).

**ovaNalk4**

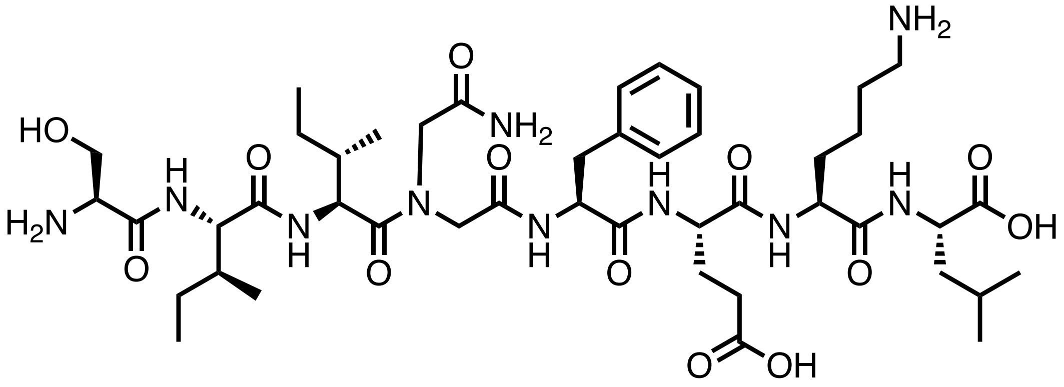


Chemical structure of **ovaNalk4**.

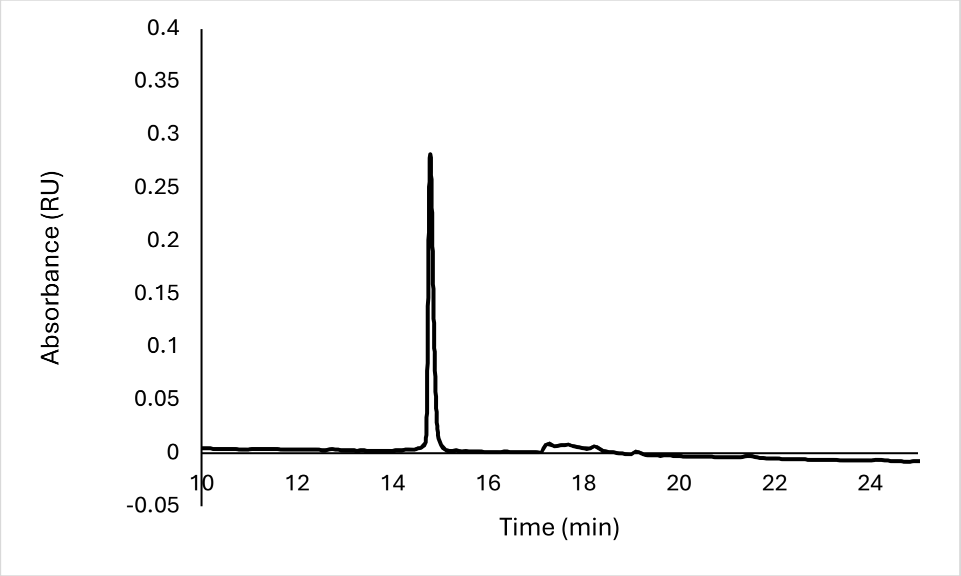


Analytical HPLC Chromatogram of **ovaNalk4**.

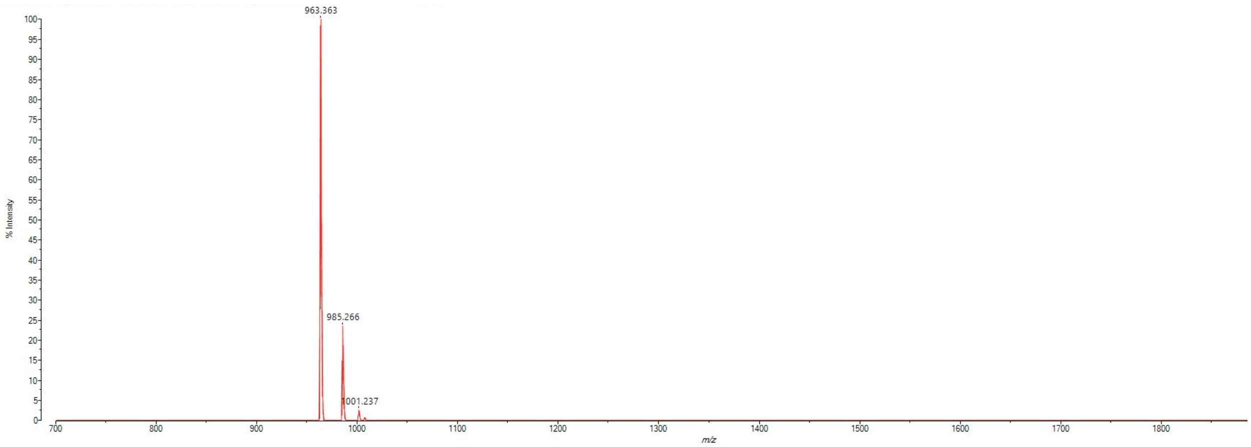


MALDI-TOF mass spectrum for **ovaNalk4** (m/z 963.551 for $[M+H]^+$, found 963.363).

**ovaNalk5**

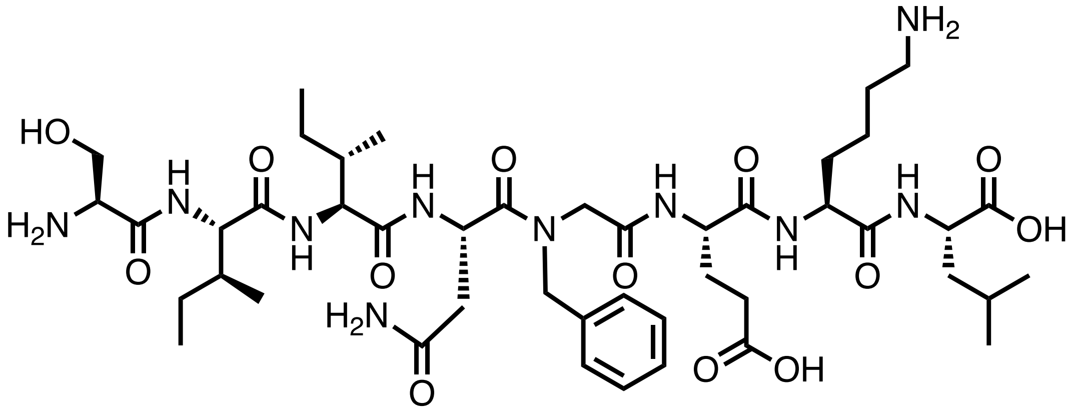


Chemical structure of **ovaNalk5**.

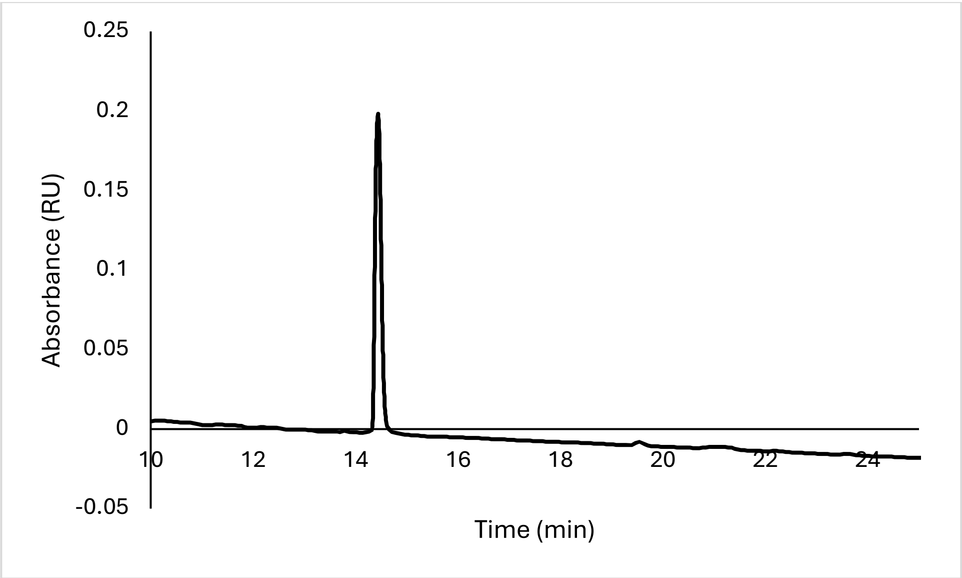


Analytical HPLC Chromatogram of **ovaNalk5**.

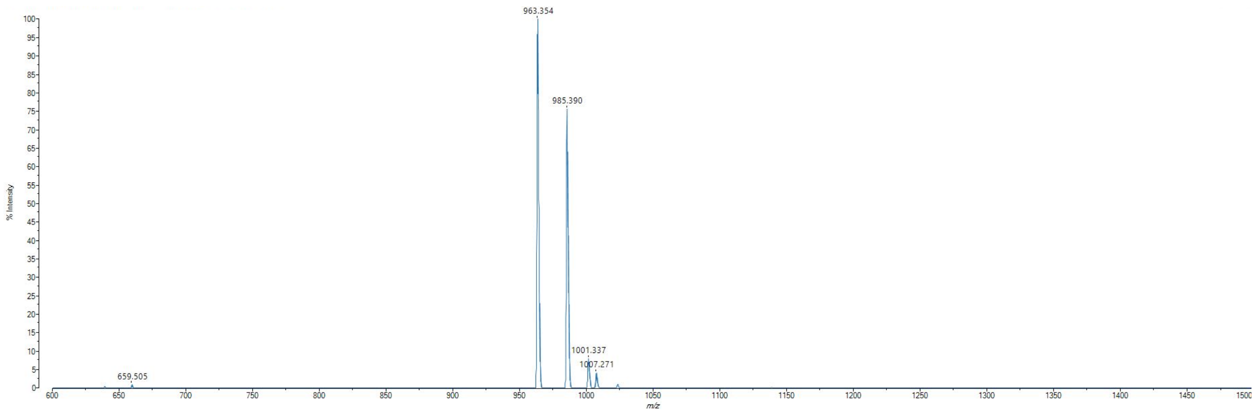


MALDI-TOF mass spectrum for **ovaNalk5** (m/z 963.551 for $[M+H]^+$, found 963.354).

**ovaNalk6**

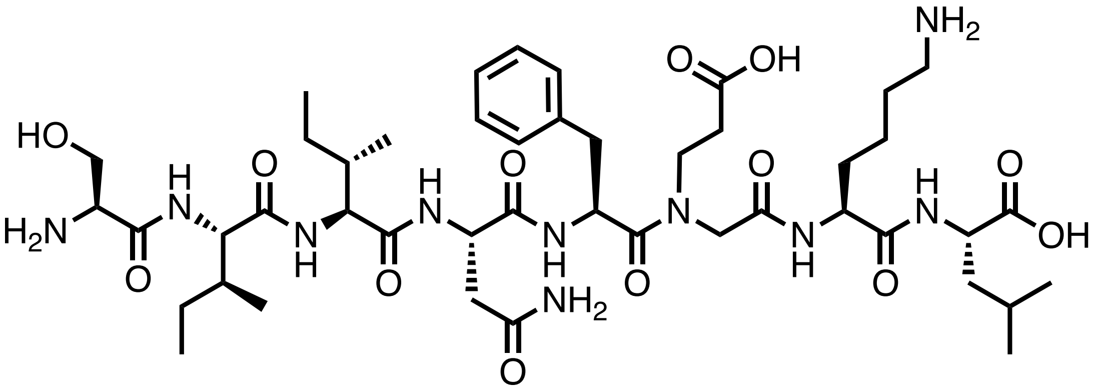

Chemical structure of **ovaNalk6**.

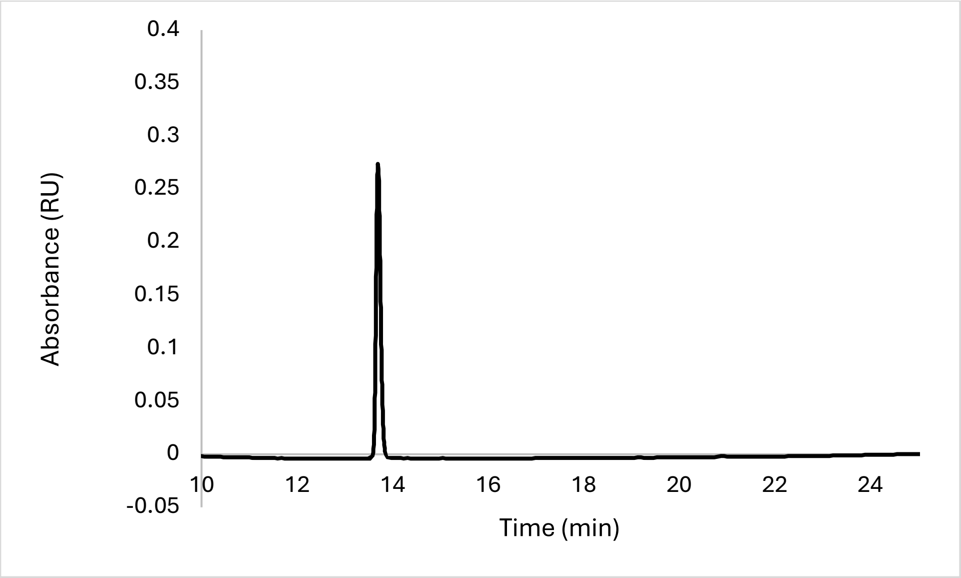


Analytical HPLC Chromatogram of **ovaNalk6**.

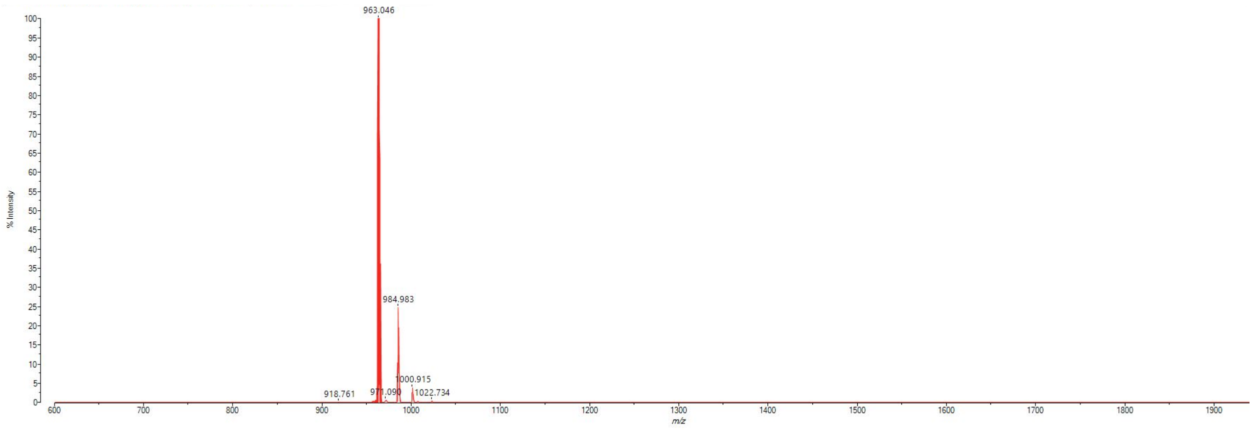


MALDI-TOF mass spectrum for **ovaNalk6** (m/z 963.551 for $[M+H]^+$, found 963.046).

**ovaNalk7**

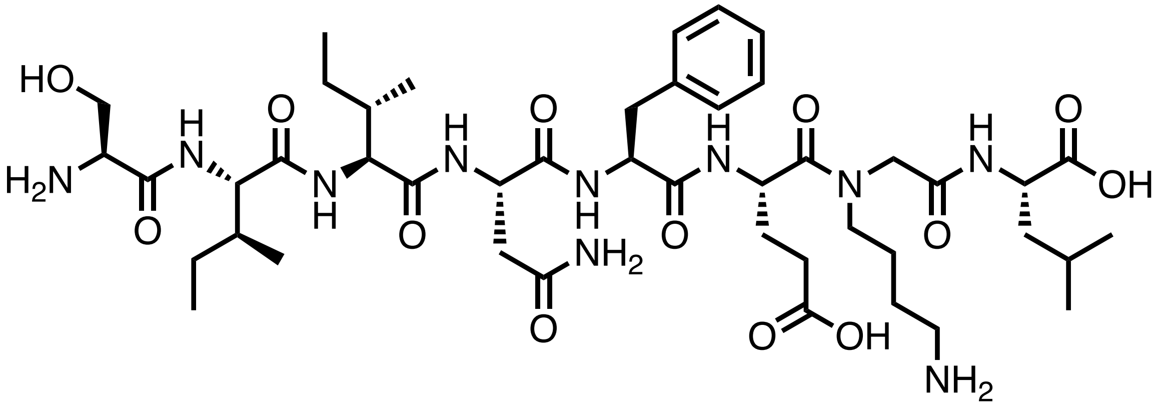

Chemical structure of **ovaNalk7**.

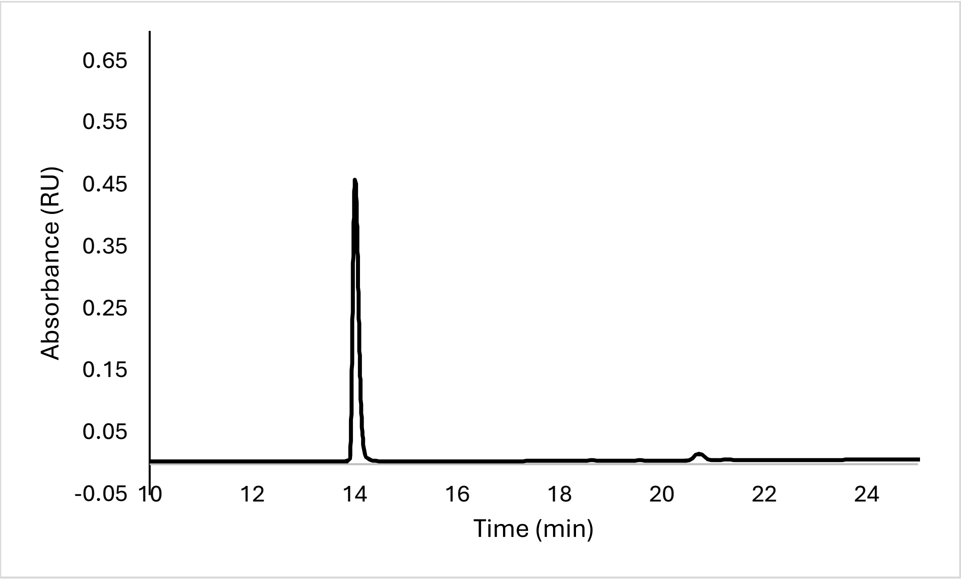


Analytical HPLC Chromatogram of **ovaNalk7**.

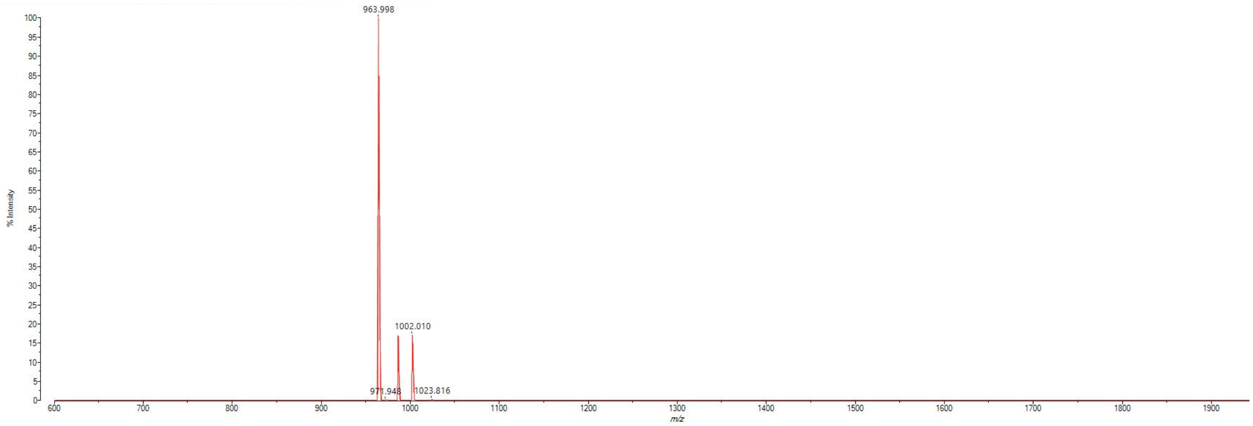


MALDI-TOF mass spectrum for **ovaNalk7** (m/z 963.551 for $[M+H]^+$, found 963.998).

**ovaNalk8**

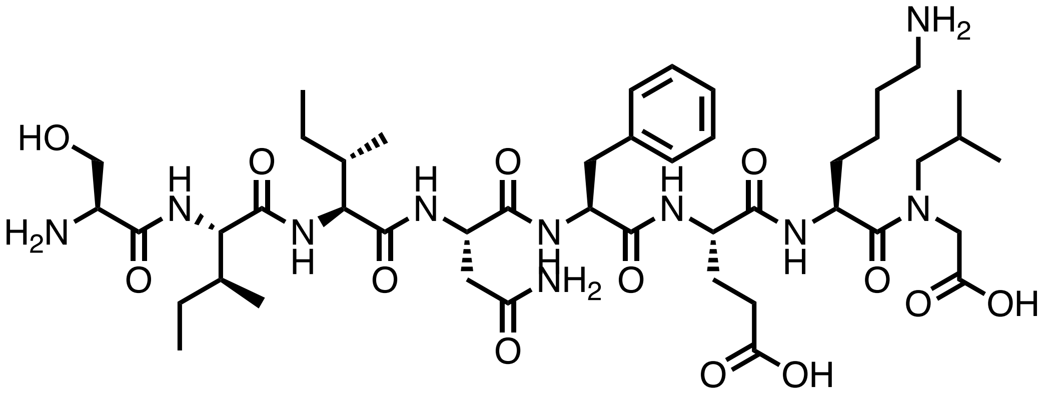

Chemical structure of **ovaNalk8**.

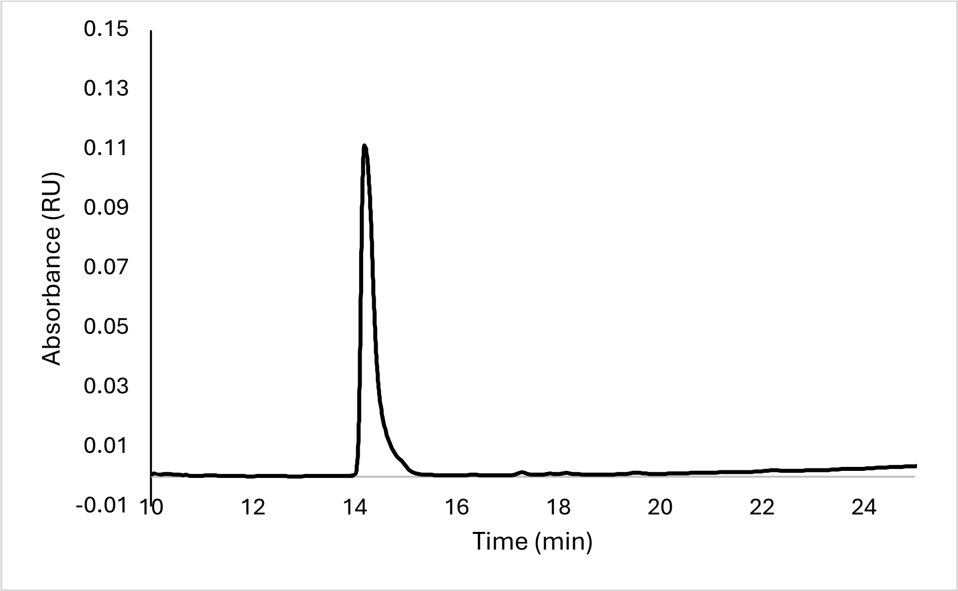


Analytical HPLC Chromatogram of **ovaNalk8**.

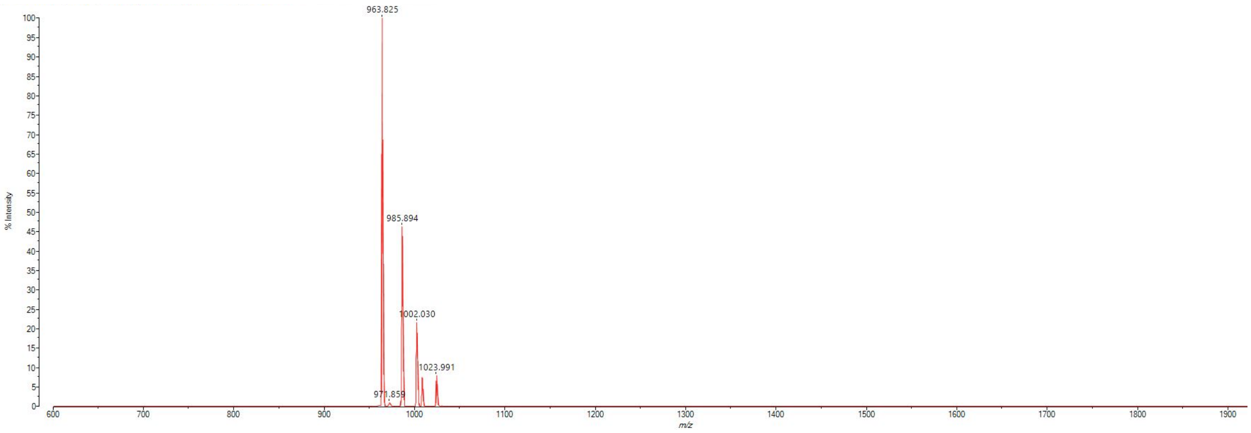


MALDI-TOF mass spectrum for **ovaNalk8** (m/z 963.551 for $[M+H]^+$, found 963.825).

**ovaD1**

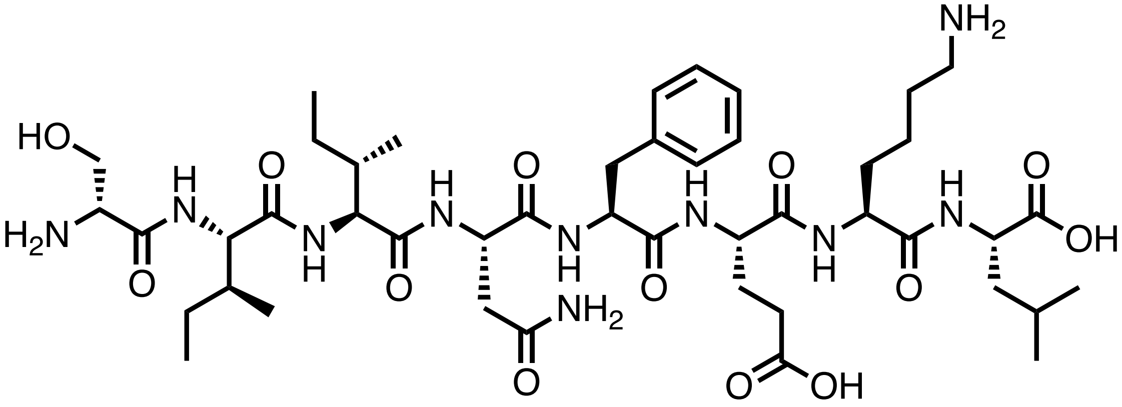

Chemical structure of **ovaD1**.

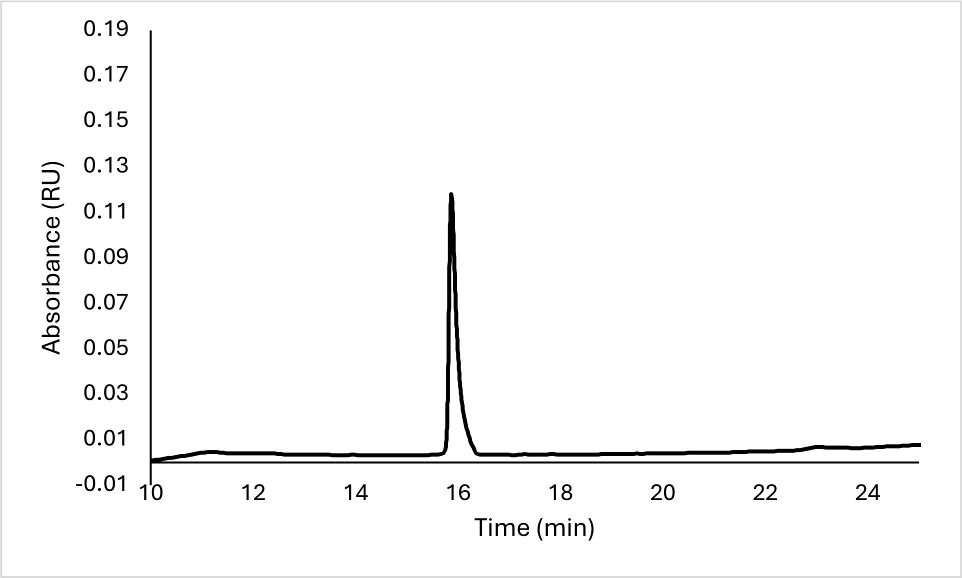


Analytical HPLC Chromatogram of **ovaD1**.

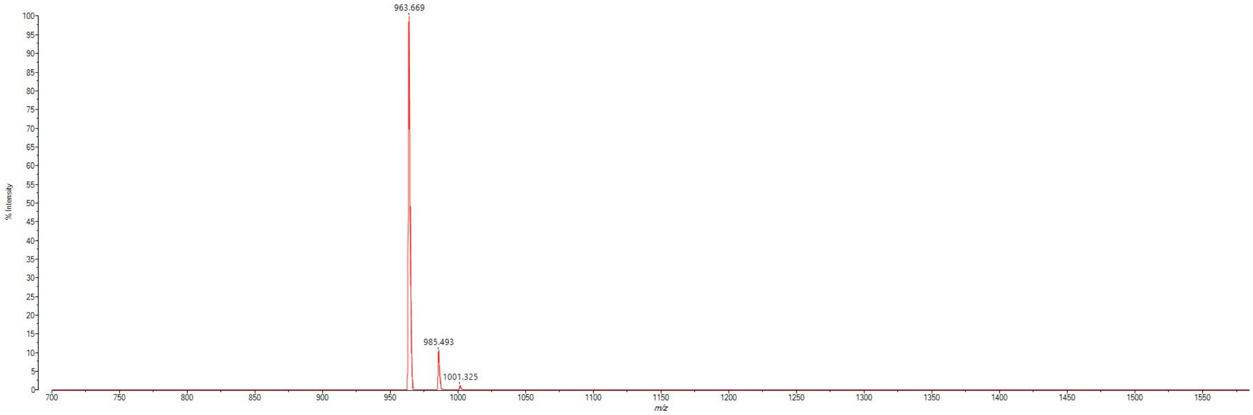


MALDI-TOF mass spectrum for **ovaD1** (m/z 963.551 for $[M+H]^+$, found 963.669).

**ovaD2**

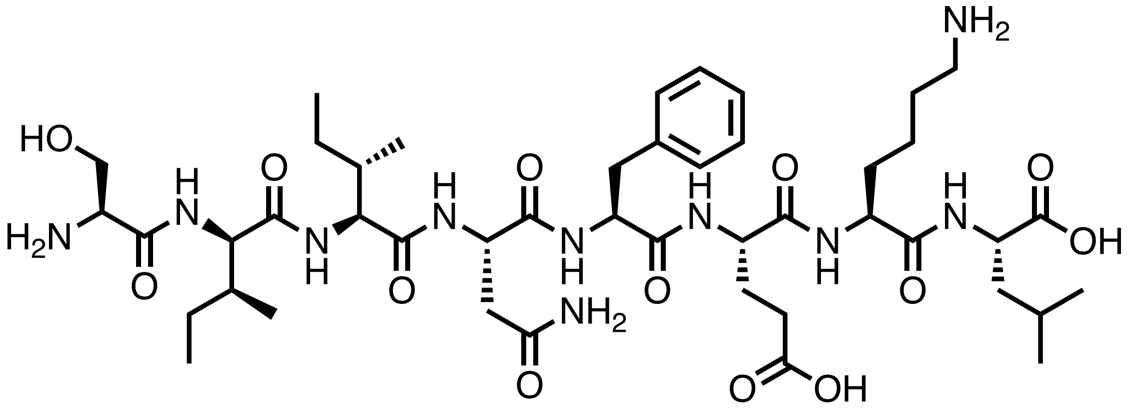


Chemical structure of **ovaD2**.

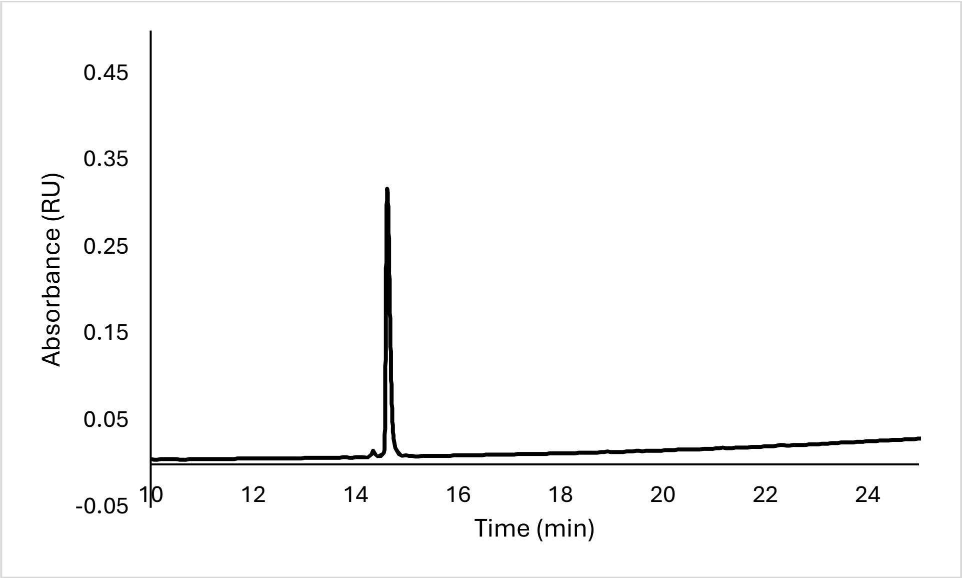


Analytical HPLC Chromatogram of **ovaD2**.

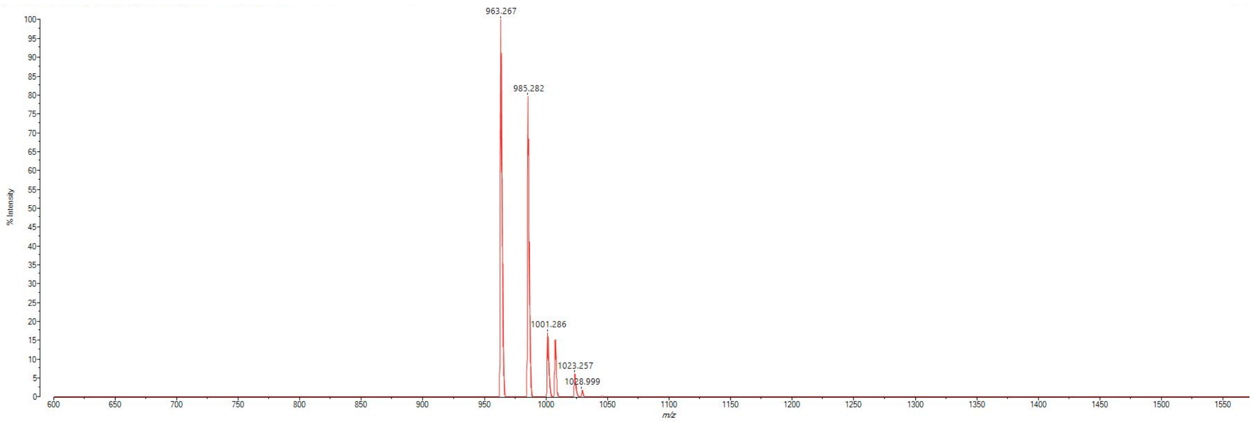


MALDI-TOF mass spectrum for **ovaD2** (m/z 963.551 for $[M+H]^+$, found 963.267).

**ovaD3**

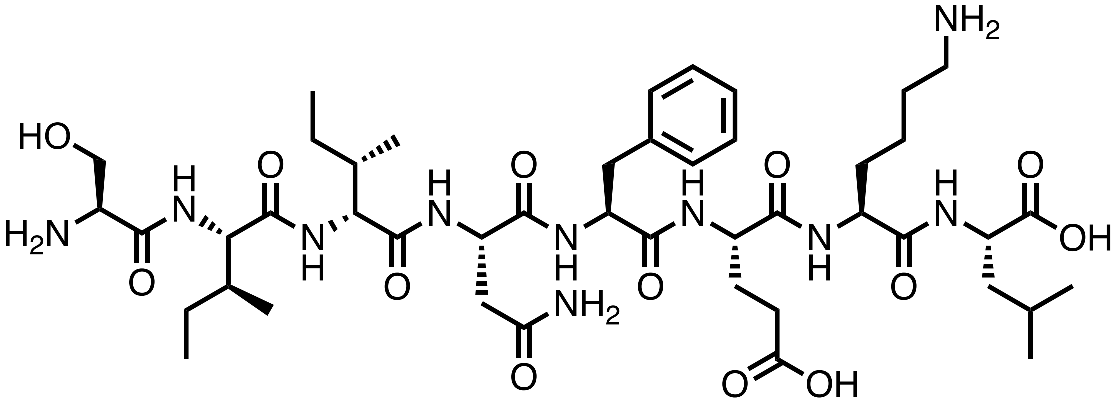

Chemical structure of **ovaD3**.

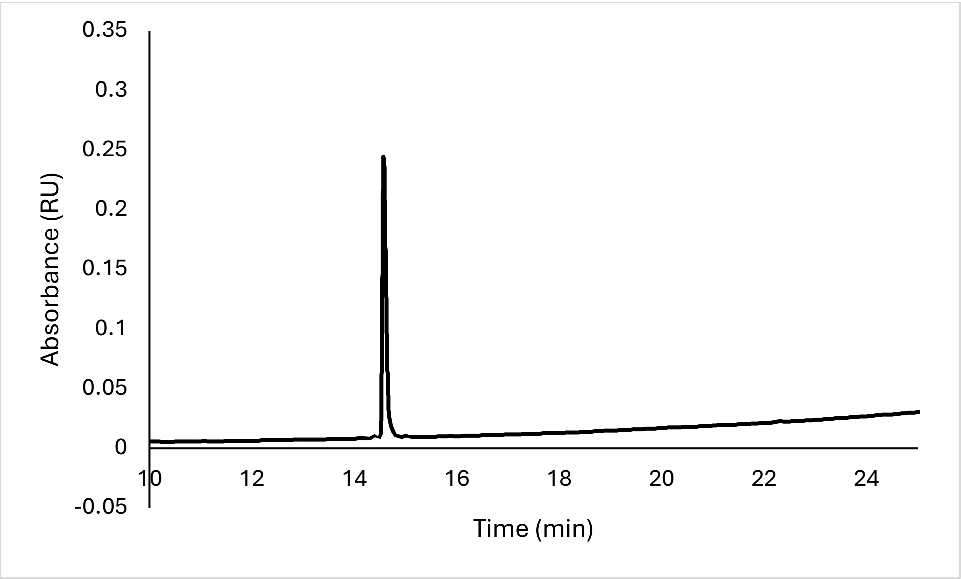


Analytical HPLC Chromatogram of **ovaD3**.

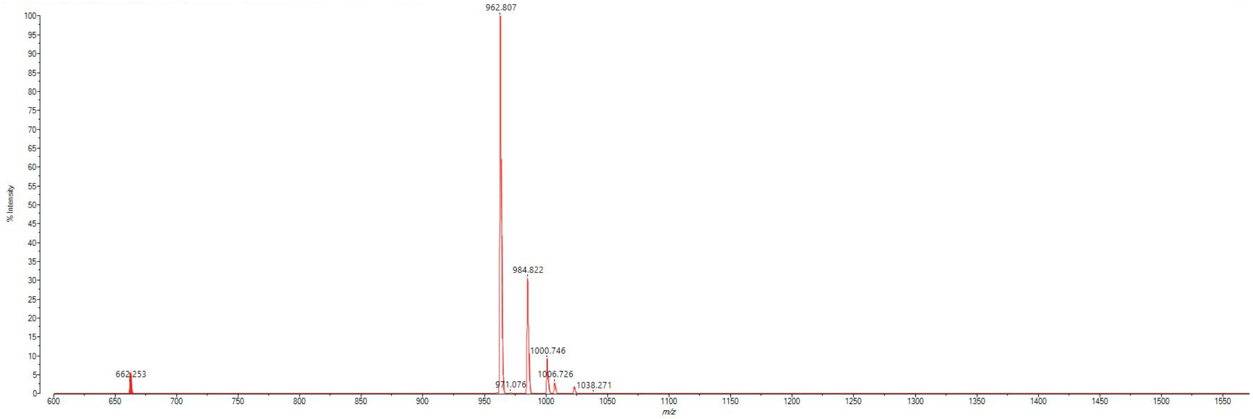


MALDI-TOF mass spectrum for **ovaD3** (m/z 963.551 for $[M+H]^+$, found 962.807).

**ovaD4**

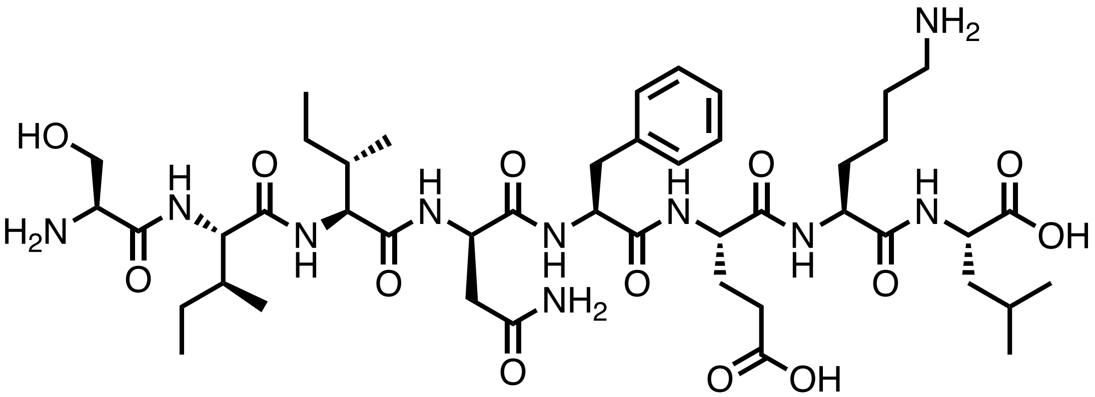

Chemical structure of **ovaD4**.

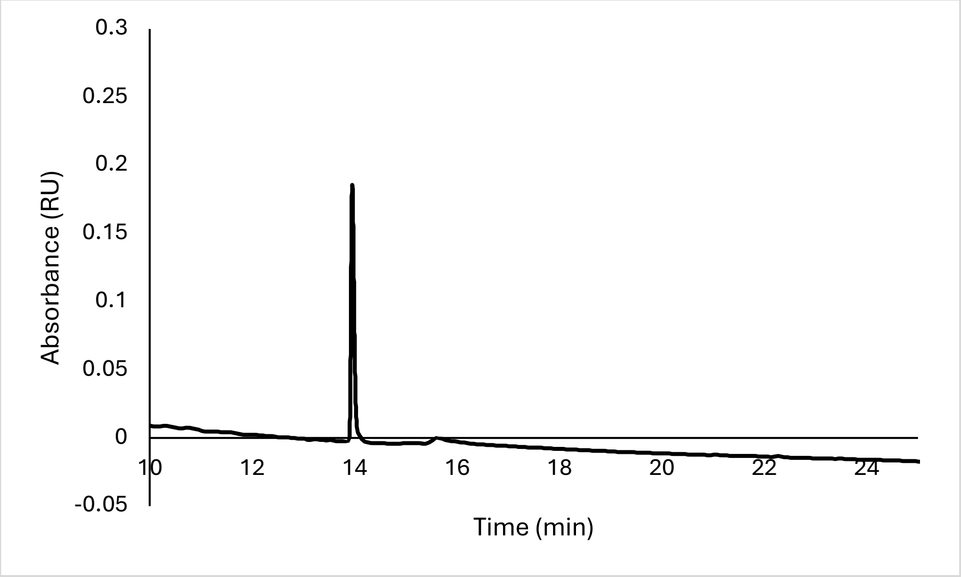


Analytical HPLC Chromatogram of **ovaD4**.

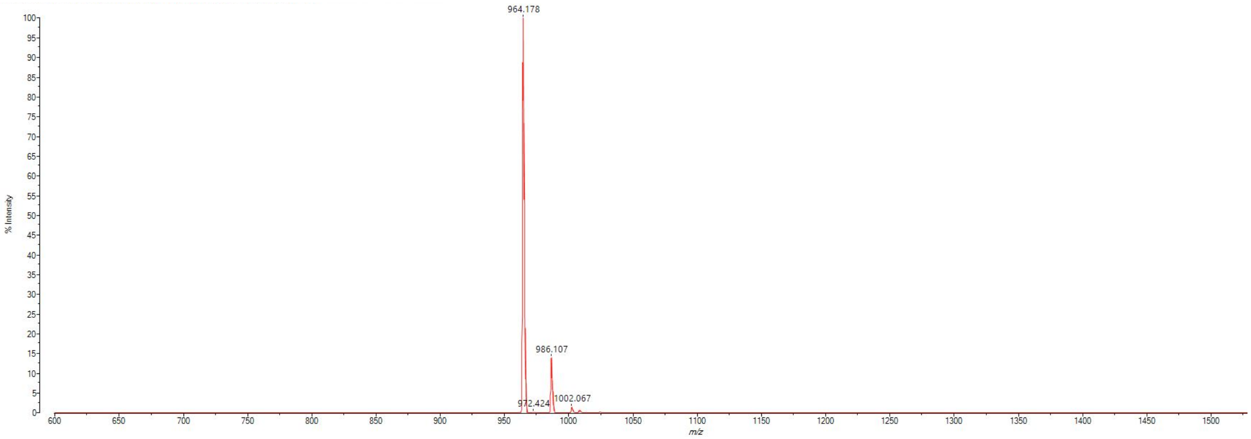


MALDI-TOF mass spectrum for **ovaD4** (m/z 963.551 for $[M+H]^+$, found 964.178).

**ovaD5**

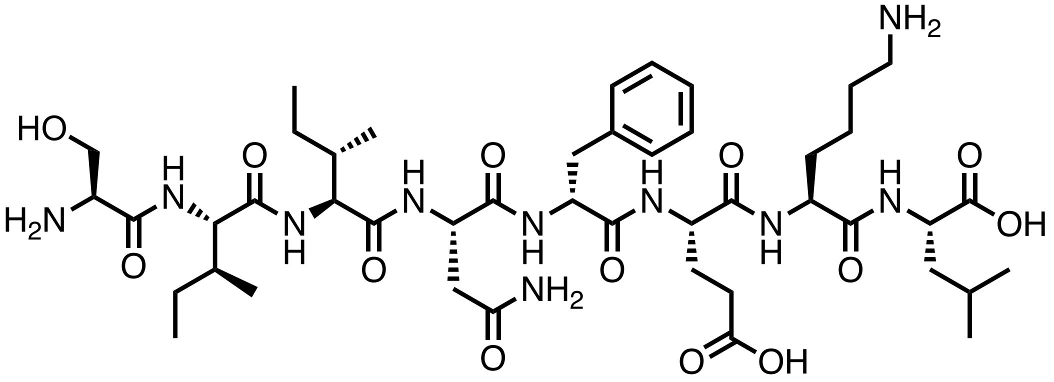

Chemical structure of **ovaD5**.

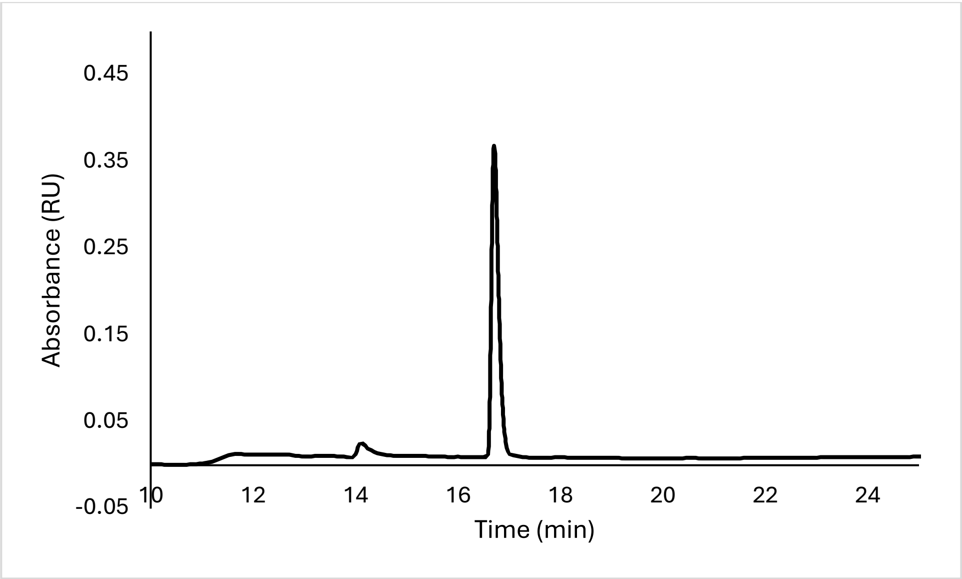


Analytical HPLC Chromatogram of **ovaD5**.

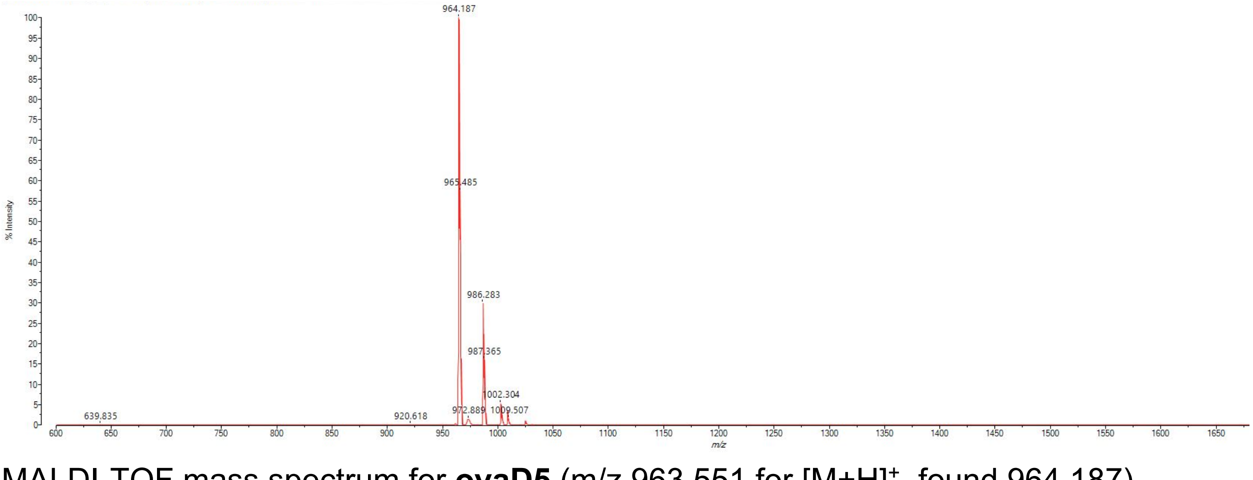


MALDI-TOF mass spectrum for **ovaD5** (m/z 963.551 for $[M+H]^+$, found 964.187).

**ovaD6**

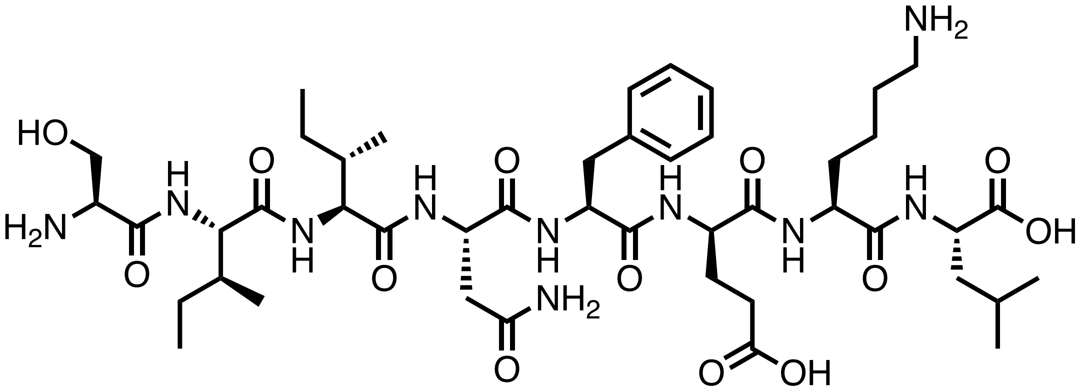

Chemical structure of **ovaD6**.

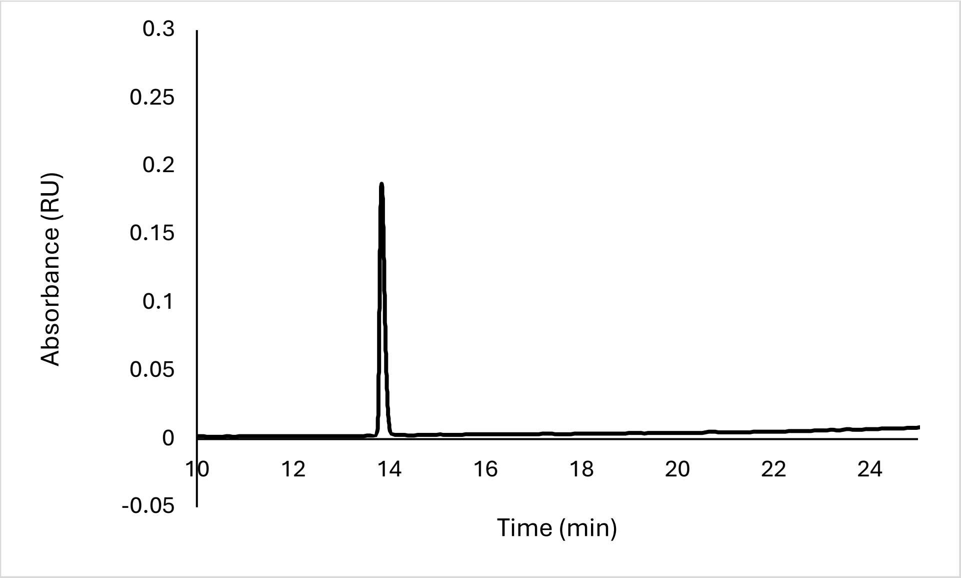


Analytical HPLC Chromatogram of **ovaD6**.

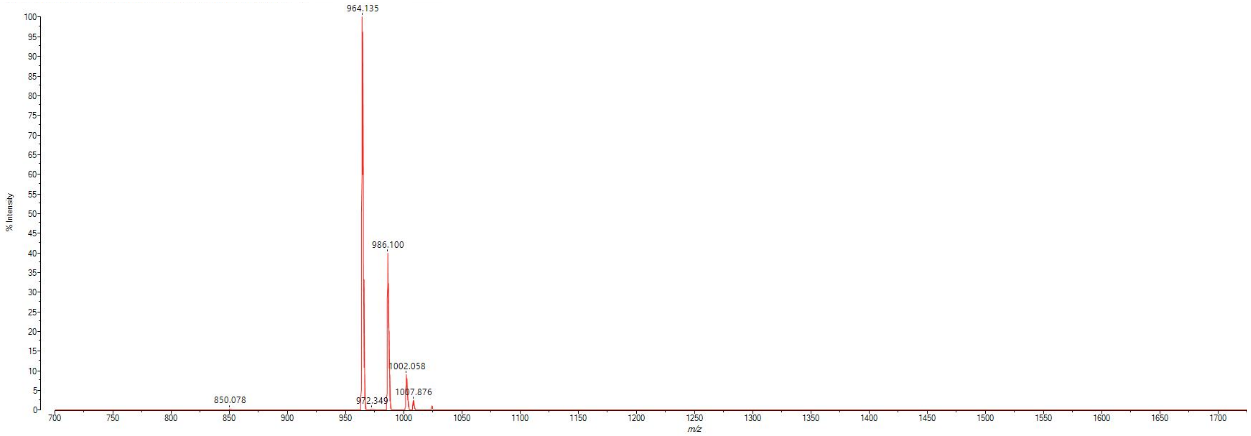


MALDI-TOF mass spectrum for **ovaD6** (m/z 963.551 for $[M+H]^+$, found 964.135).

**ovaD7**

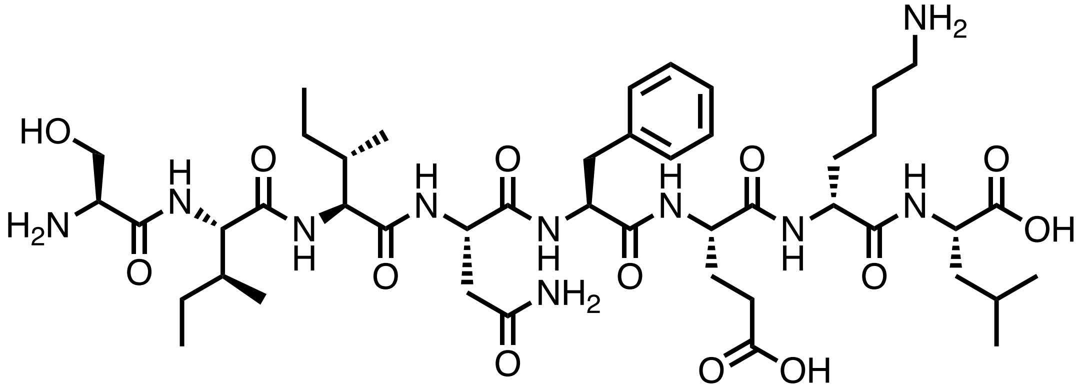

Chemical structure of **ovaD7**.

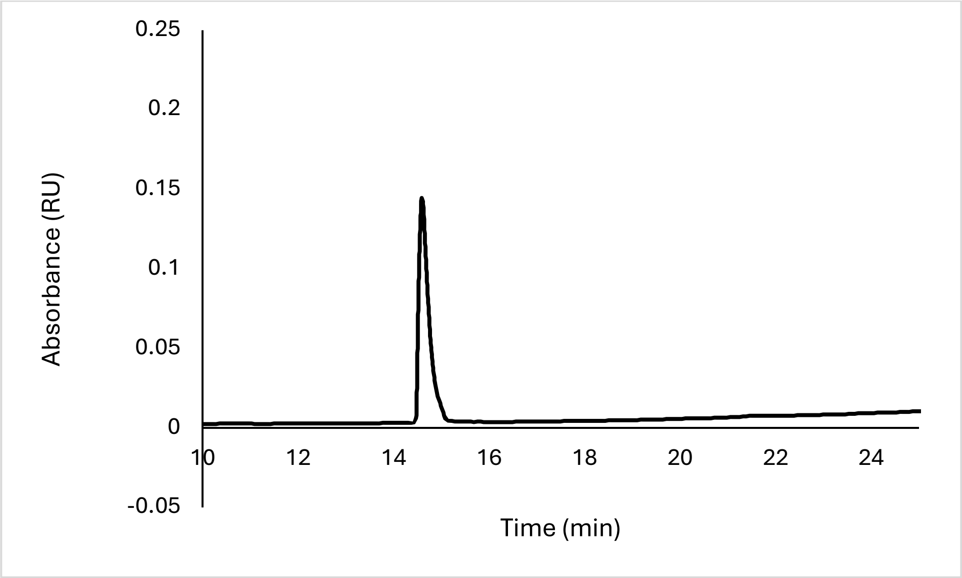


Analytical HPLC Chromatogram of **ovaD7**.

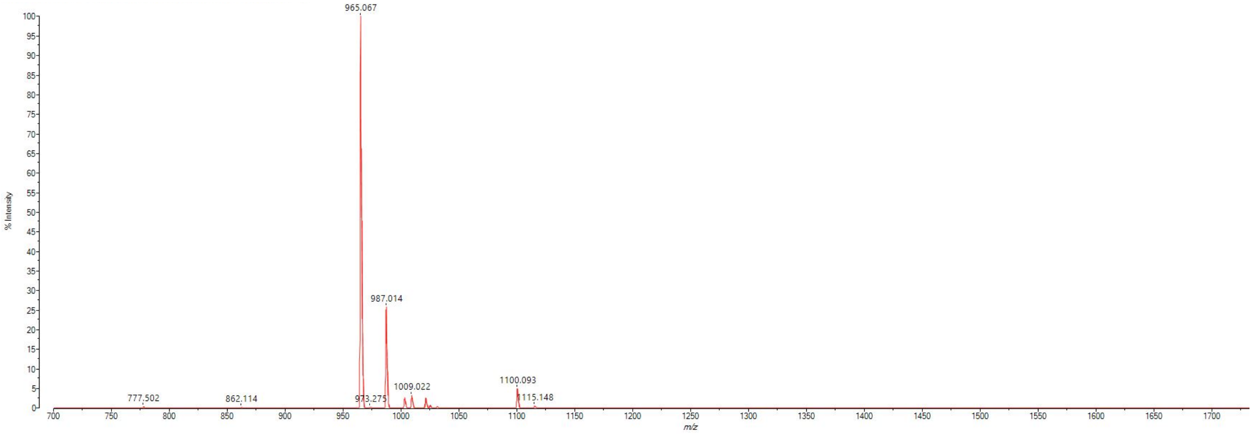


MALDI-TOF mass spectrum for **ovaD7** (m/z 963.551 for $[M+H]^+$, found 965.067).

**ovaD8**

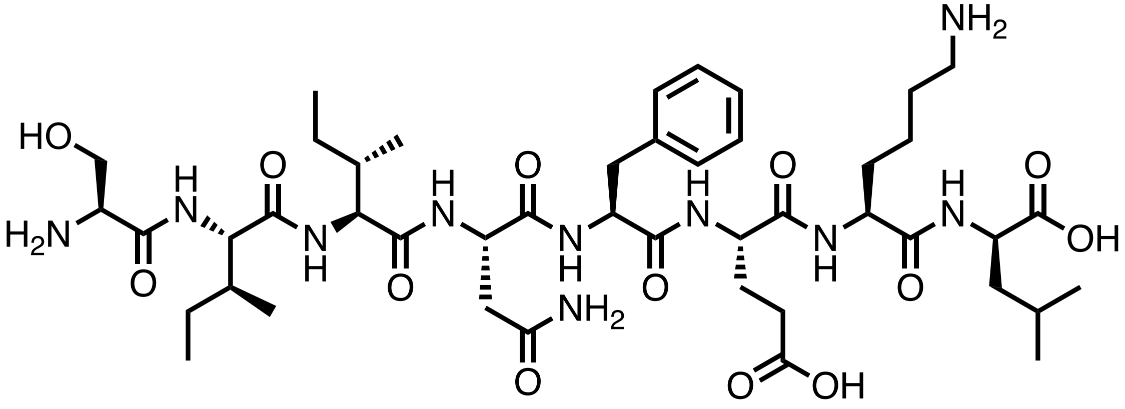


Chemical structure of **ovaD8**.

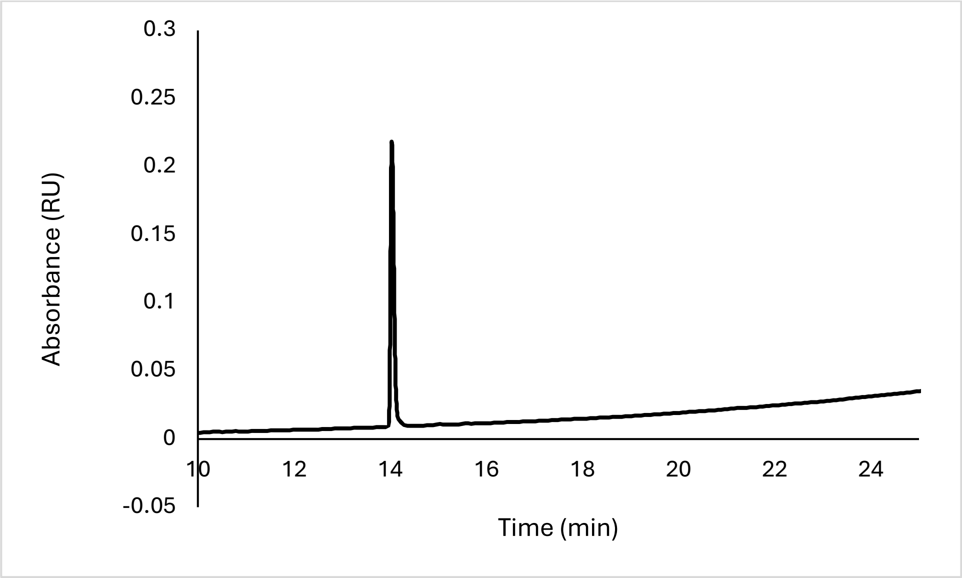


Analytical HPLC Chromatogram of **ovaD8**.

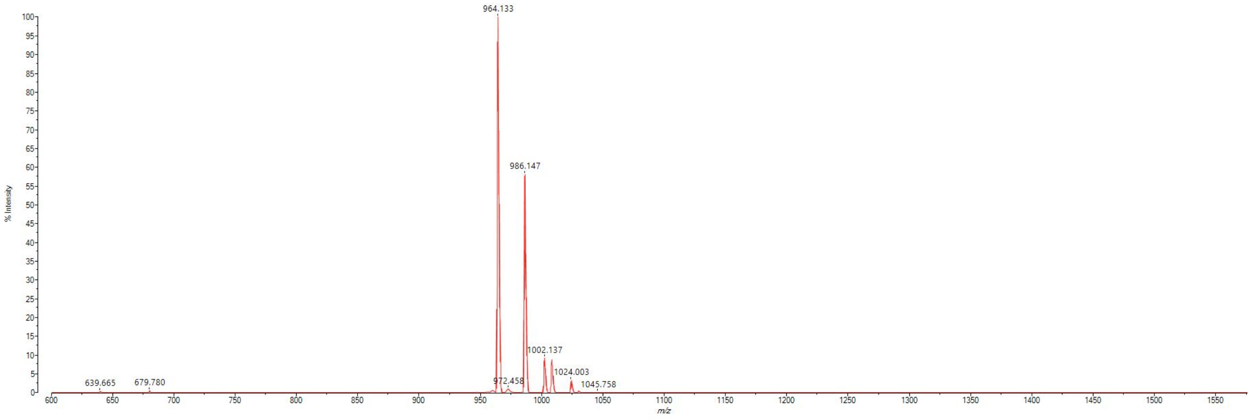


MALDI-TOF mass spectrum for **ovaD8** (m/z 963.551 for $[M+H]^+$, found 964.133).

**ovaRI**

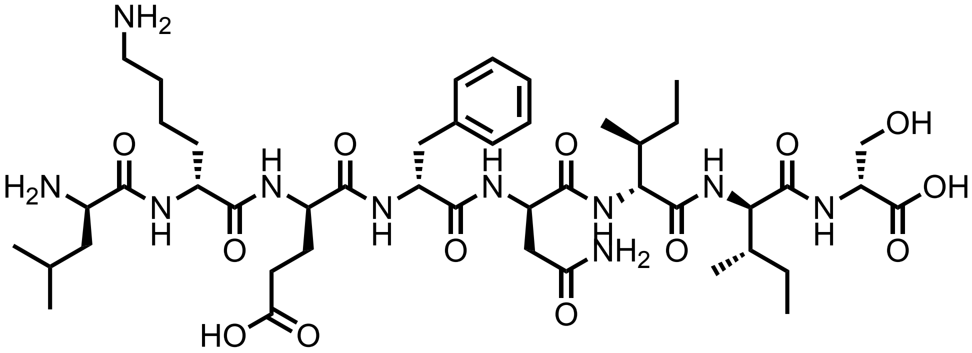


Chemical structure of **ovaRI**.

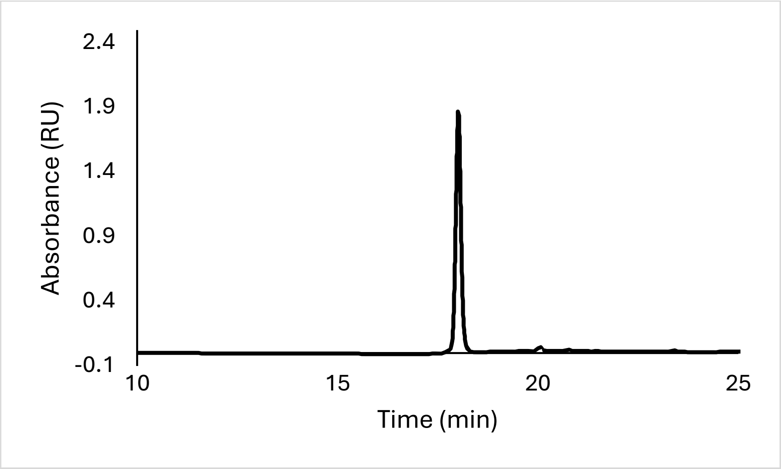


Analytical HPLC Chromatogram of **ovaRI**.

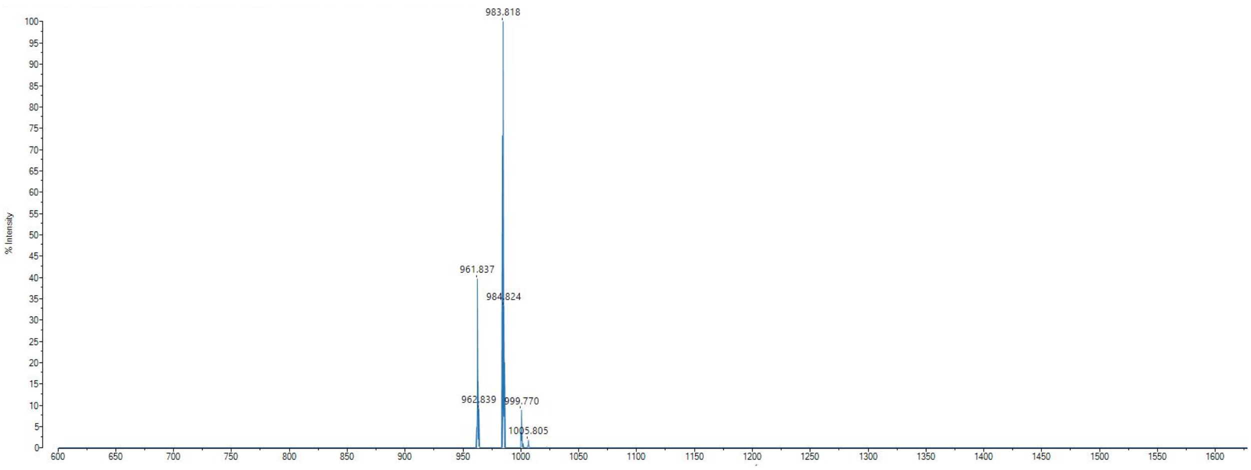


MALDI-TOF mass spectrum for **ovaRI** (m/z 963.551 for $[M+H]^+$, found 961.837).

**ovaDiMod**

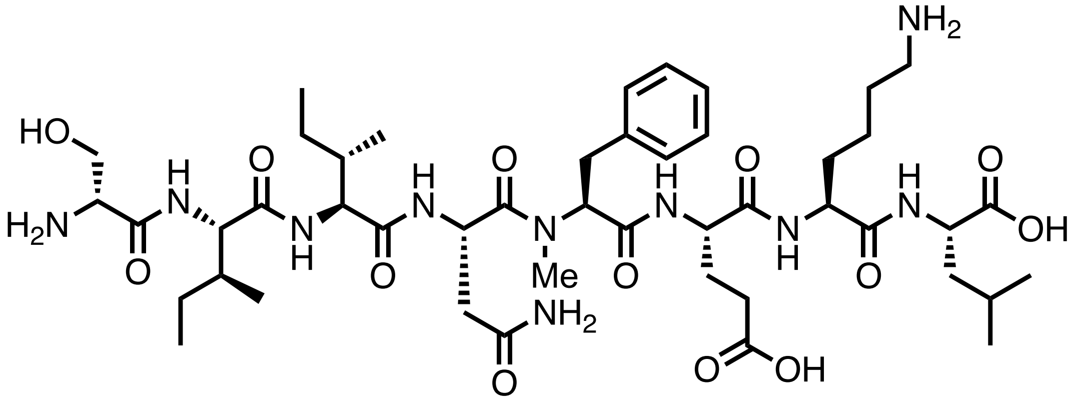

Chemical structure of **ovaDiMod**.

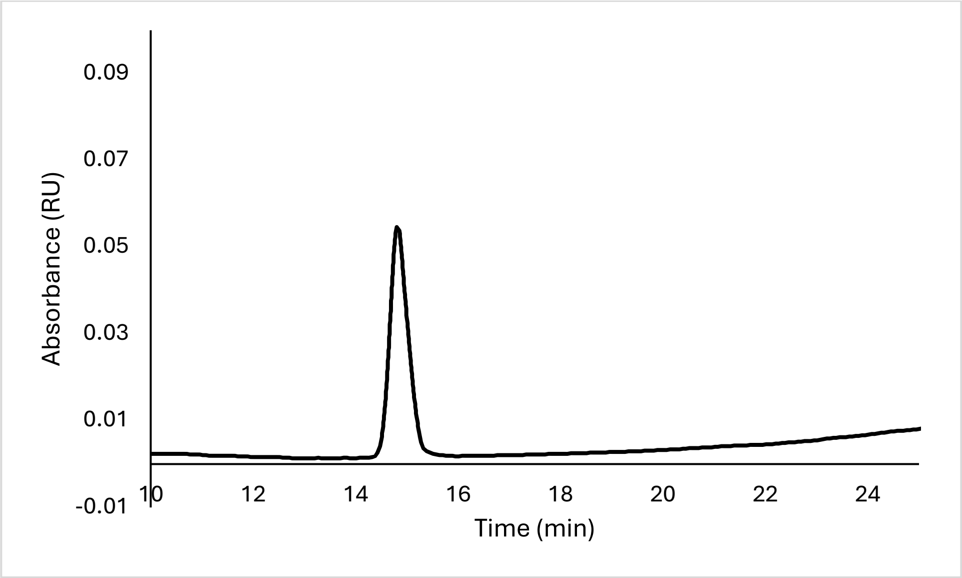


Analytical HPLC Chromatogram of **ovaDiMod**.

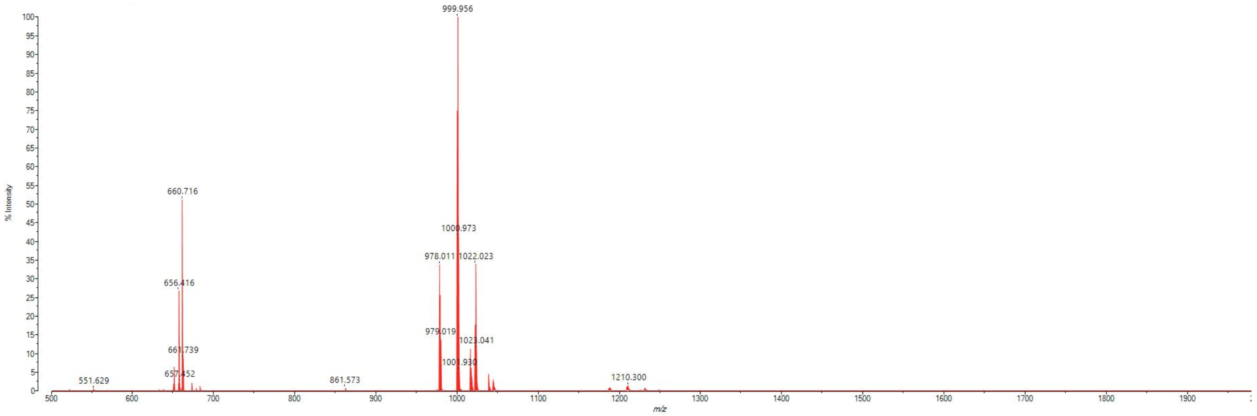


MALDI-TOF mass spectrum for **ovaDiMod** (m/z 977.567 for $[M+H]^+$, found 978.011).

**ovaTriMod**

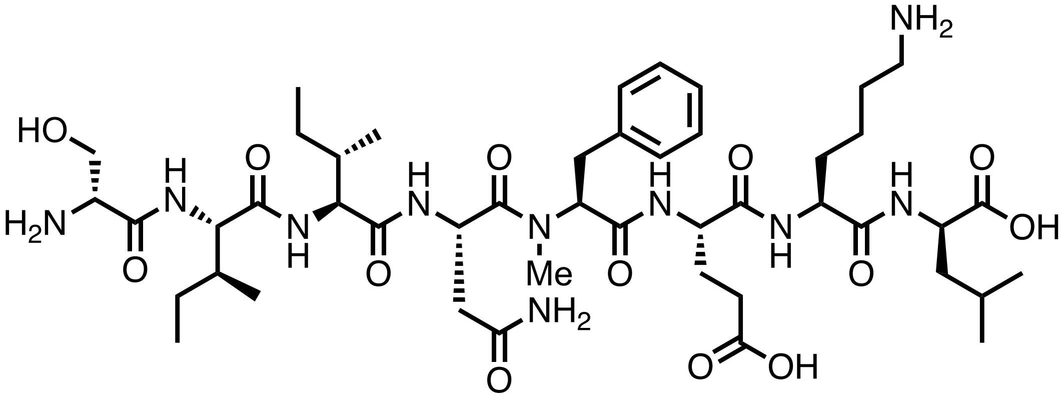

Chemical structure of **ovaTriMod**.

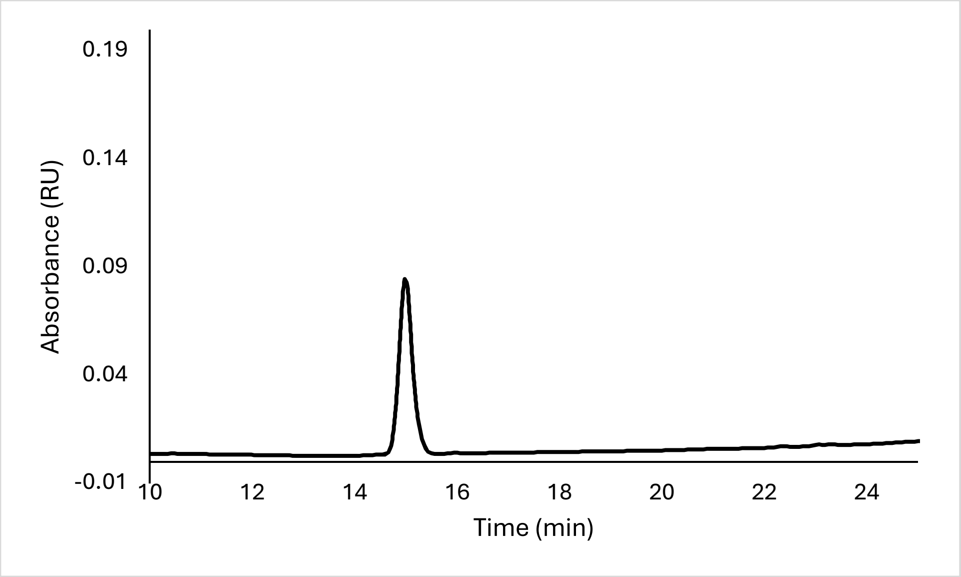


Analytical HPLC Chromatogram of **ovaTriMod**.

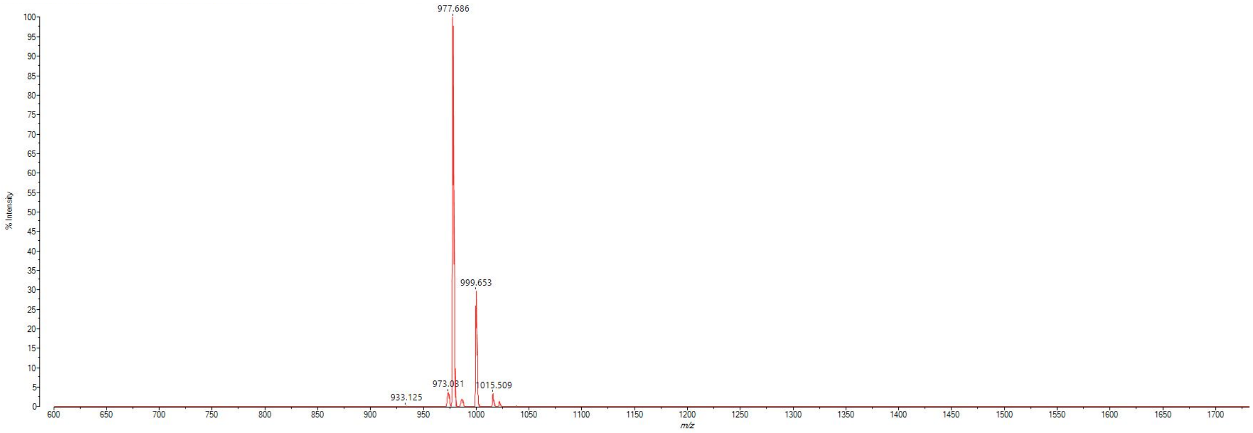


MALDI-TOF mass spectrum for **ovaTriMod** (m/z 977.567 for $[M+H]^+$, found 977.686).

**az-ovaWT**

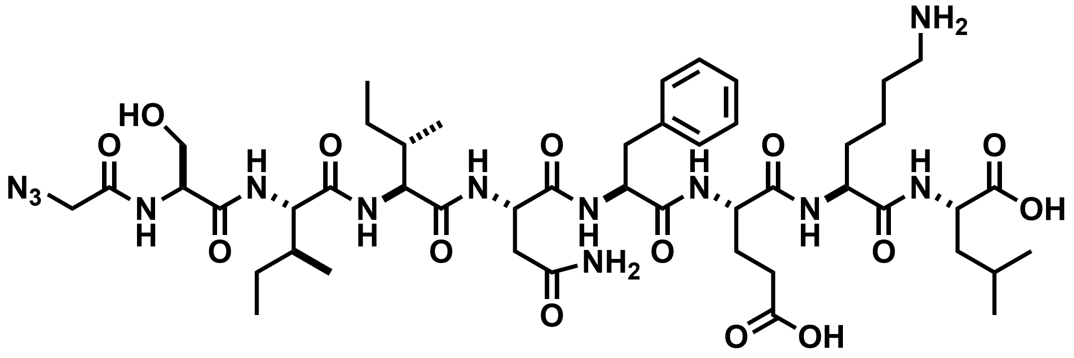


Chemical structure of **az-ovaWT**.

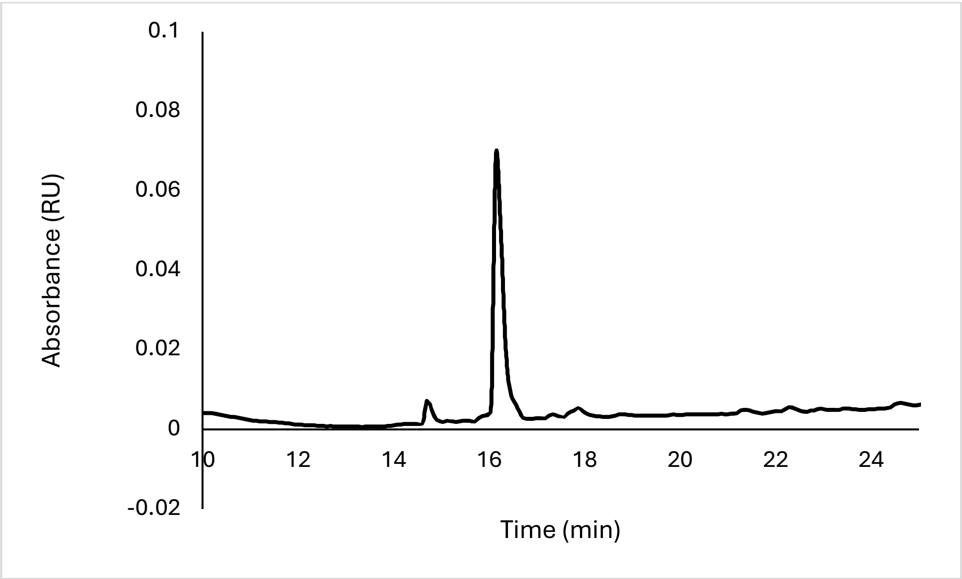


Analytical HPLC Chromatogram of **az-ovaWT**.

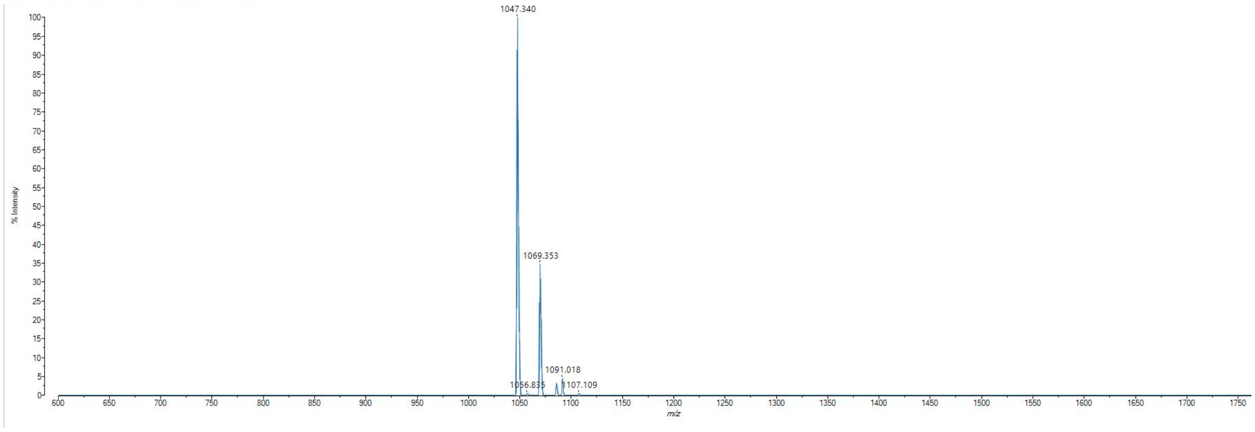


MALDI-TOF mass spectrum for **az-ovaWT** (m/z 1046.563 for $[M+H]^+$, found 1047.340).

**ovaNmet1**

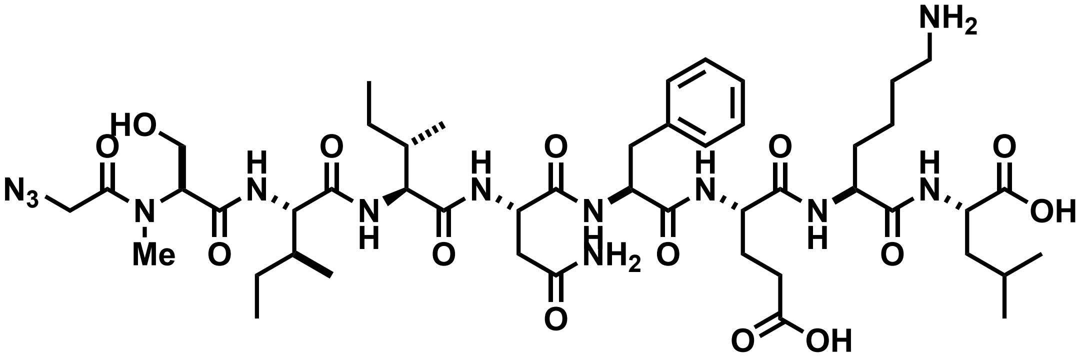

Chemical structure of **az-ovaNmet1**.

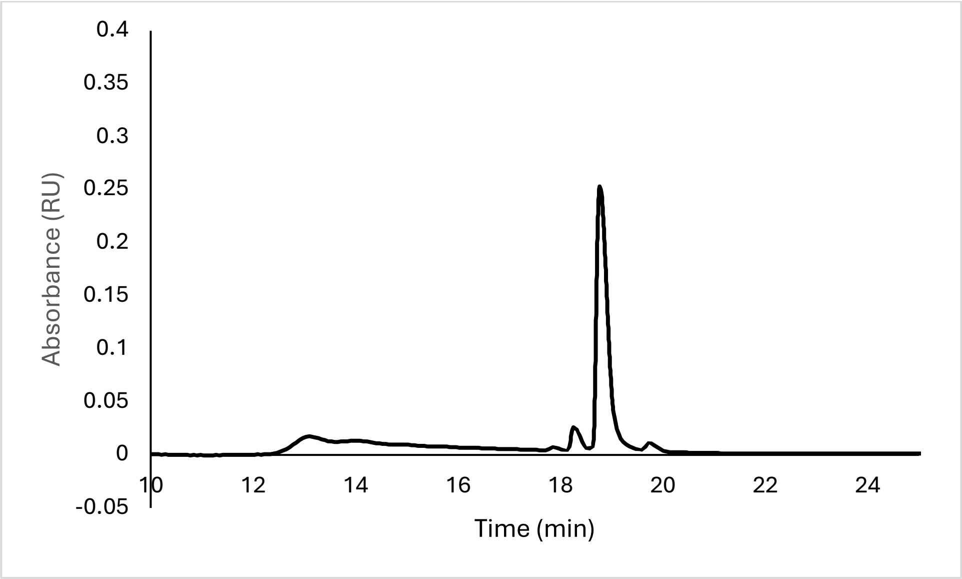


Analytical HPLC Chromatogram of **az-ovaNmet1**.

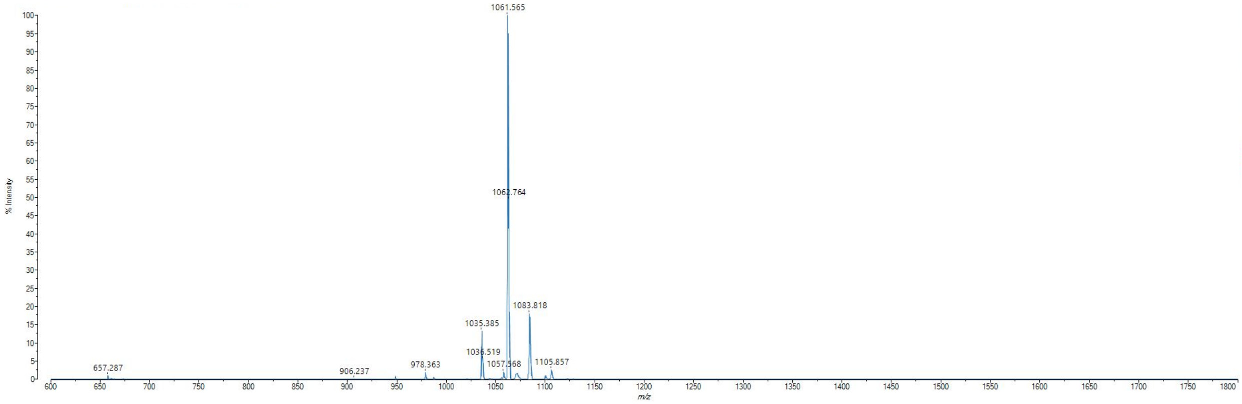


MALDI-TOF mass spectrum for **az-ovaNmet1** (m/z 1060.579 for $[M+H]^+$, found 1061.565).

**az-ovaNmet2**

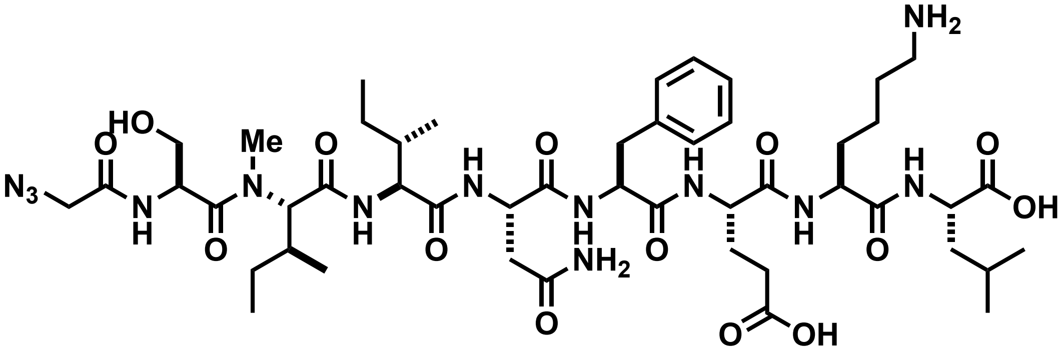


Chemical structure of **az-ovaNmet2**.

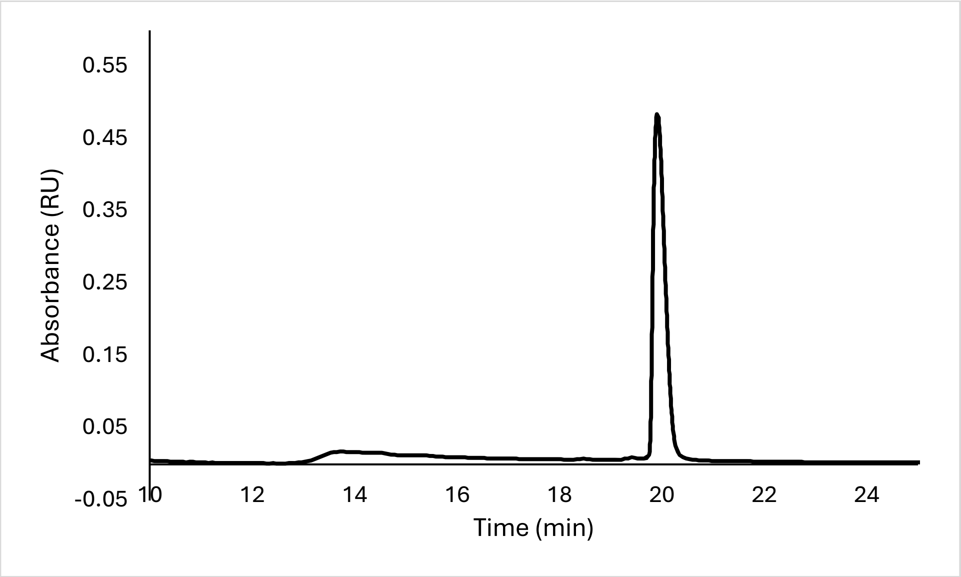


Analytical HPLC Chromatogram of **az-ovaNmet2**.

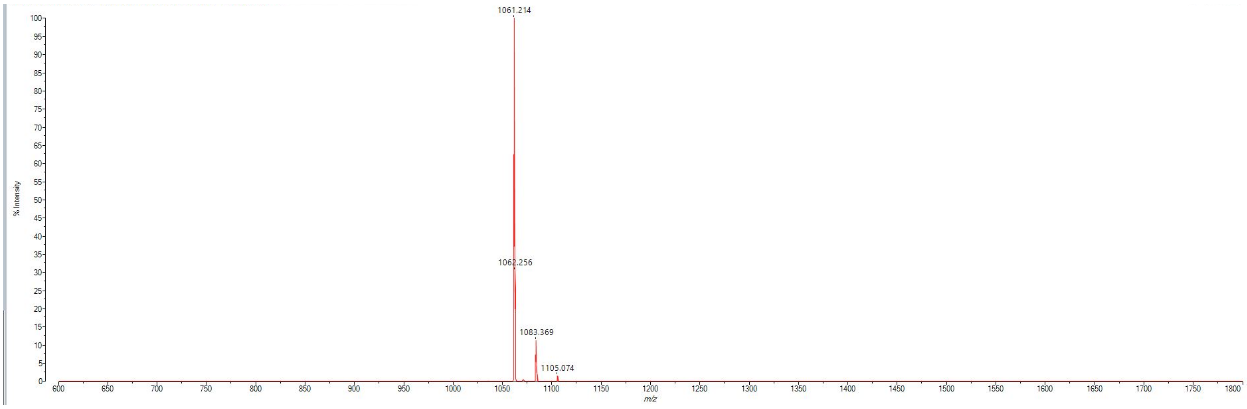


MALDI-TOF mass spectrum for **az-ovaNmet2** (m/z 1060.579 for $[M+H]^+$, found 1061.214).

**az-ovaNmet3**

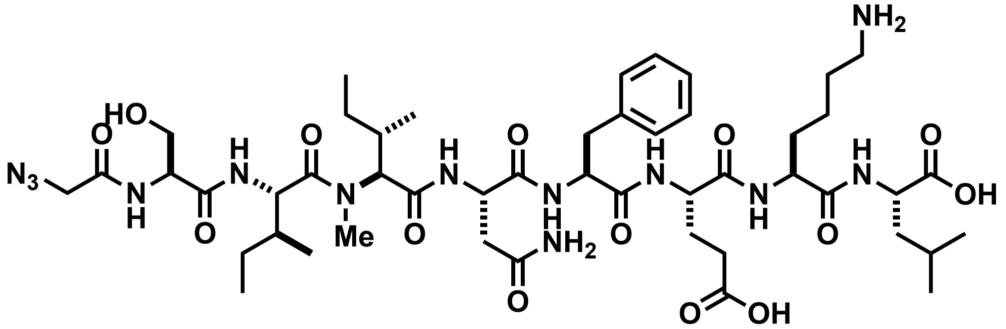


Chemical structure of **az-ovaNmet3**.

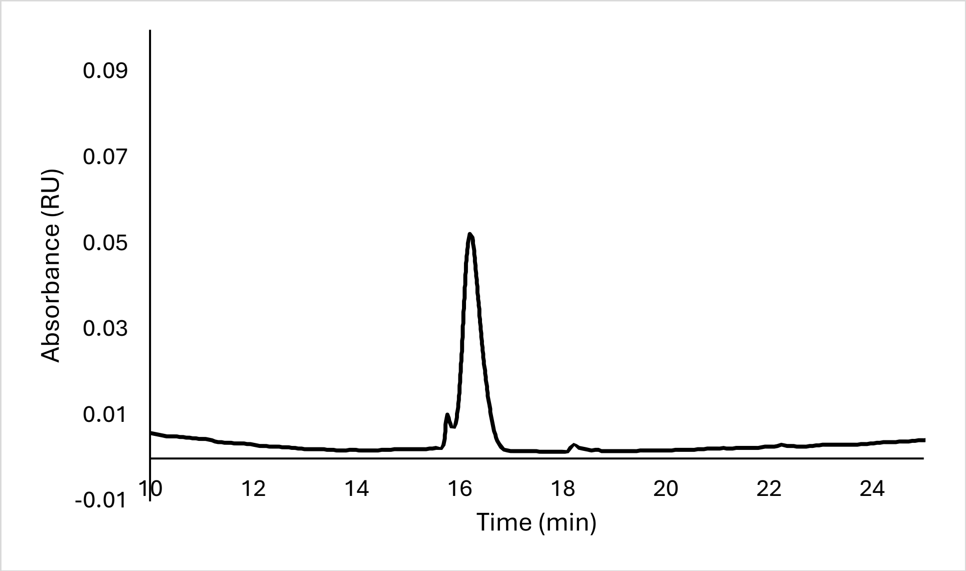


Analytical HPLC Chromatogram of **az-ovaNmet3**.

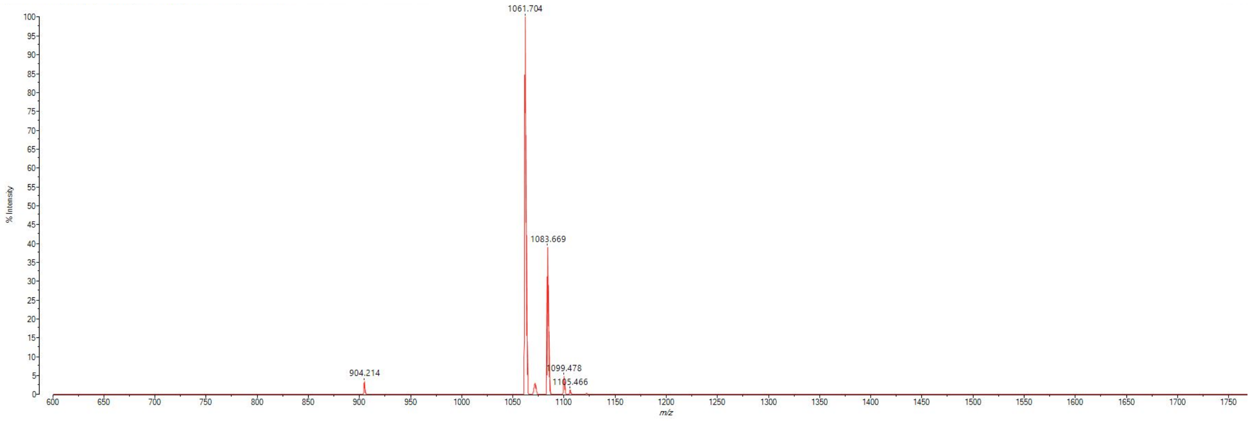


MALDI-TOF mass spectrum for **az-ovaNmet3** (m/z 1060.579 for $[M+H]^+$, found 1061.704).

**az-ovaNmet4**

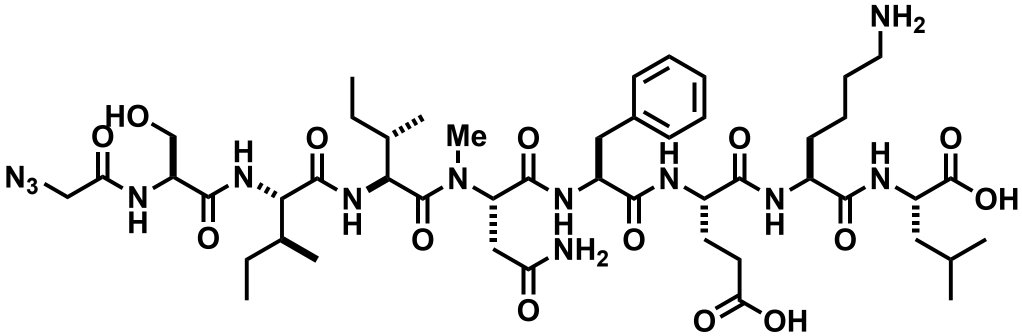

Chemical structure of **az-ovaNmet4**.

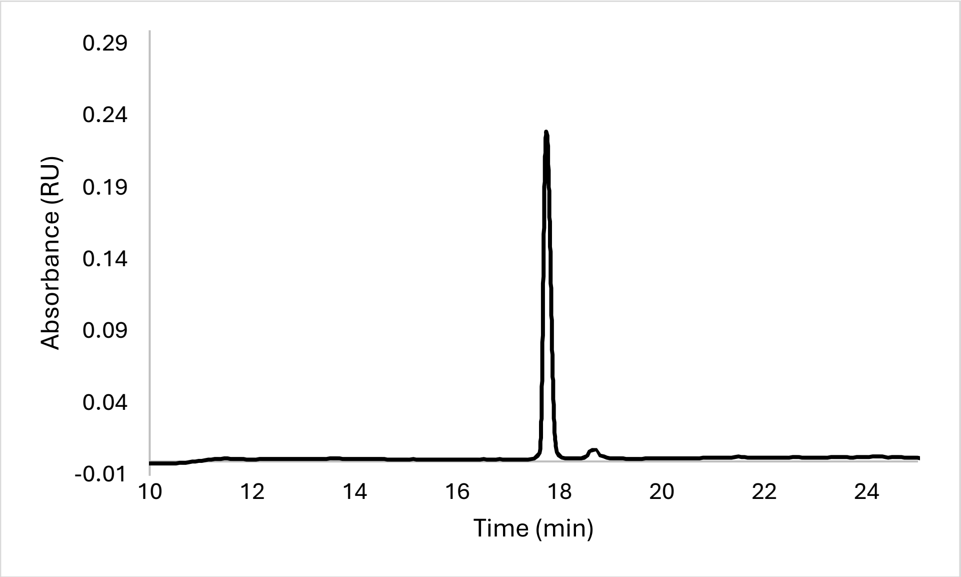


Analytical HPLC Chromatogram of **az-ovaNmet4**.

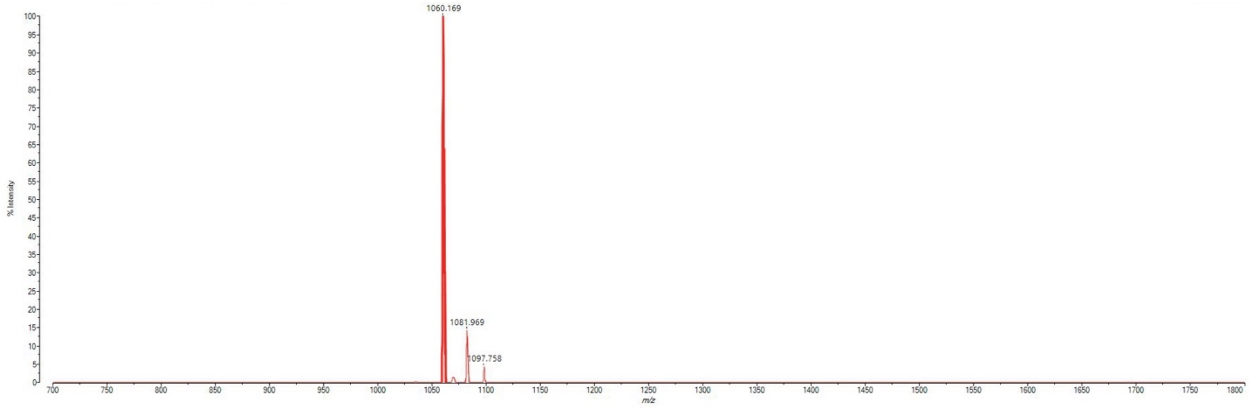


MALDI-TOF mass spectrum for **az-ovaNmet4** (m/z 1060.579 for $[M+H]^+$, found 1060.169).

**az-ovaNmet5**

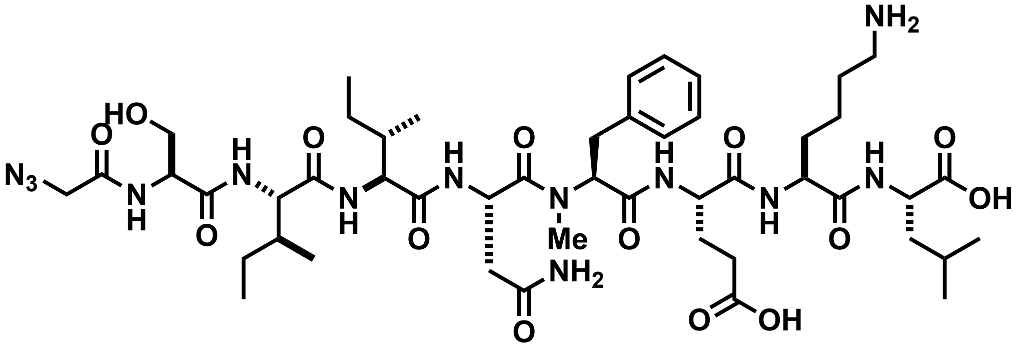

Chemical structure of **az-ovaNmet5**.

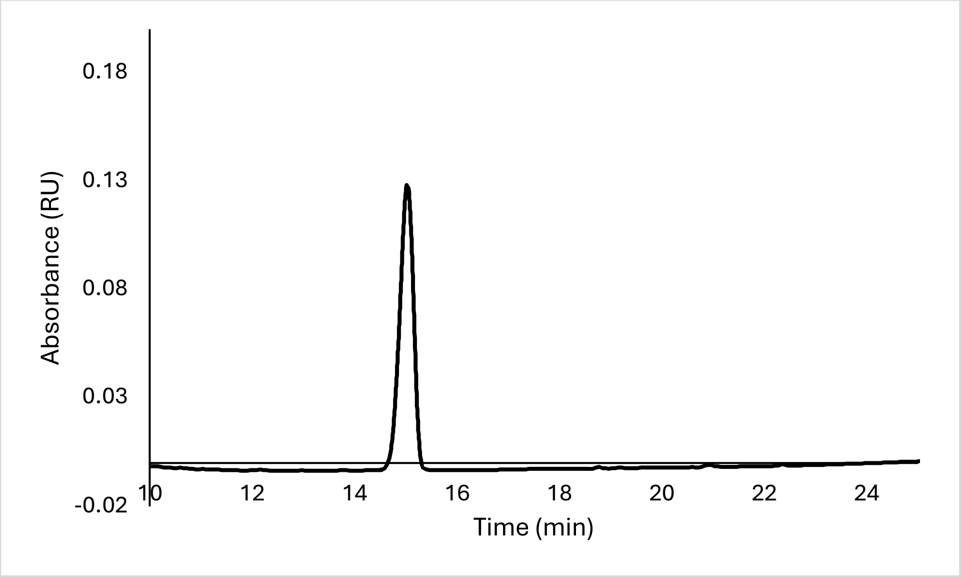


Analytical HPLC Chromatogram of **az-ovaNmet5**.

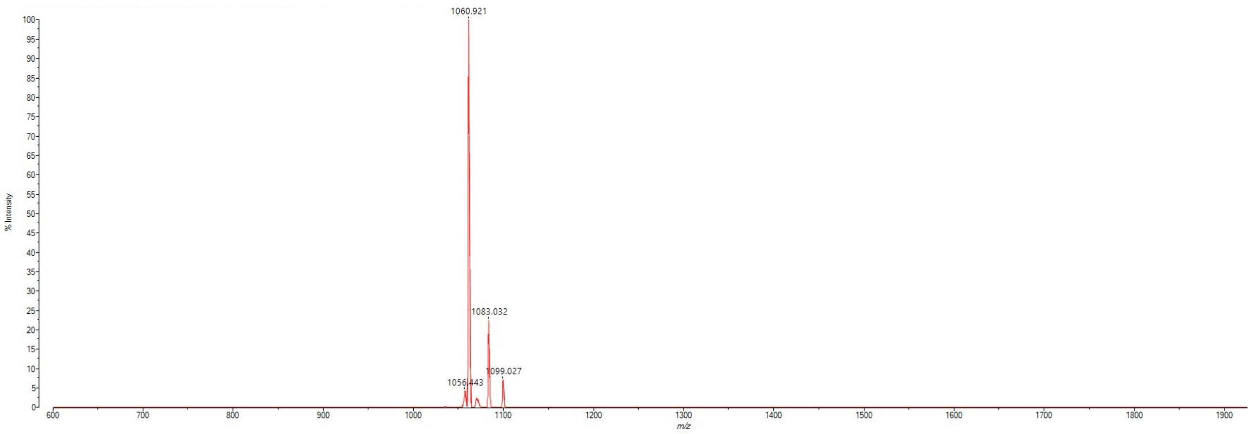


MALDI-TOF mass spectrum for **az-ovaNmet5** (m/z 1060.579 for $[M+H]^+$, found 1060.921).

**az-ovaNmet6**

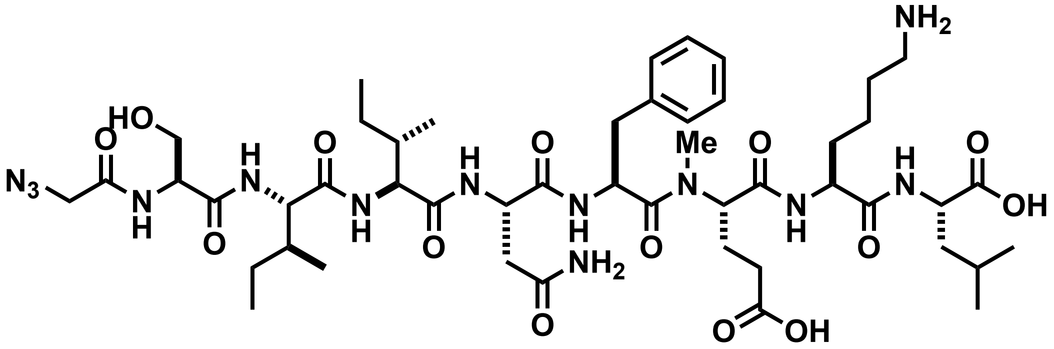


Chemical structure of **az-ovaNmet6**.

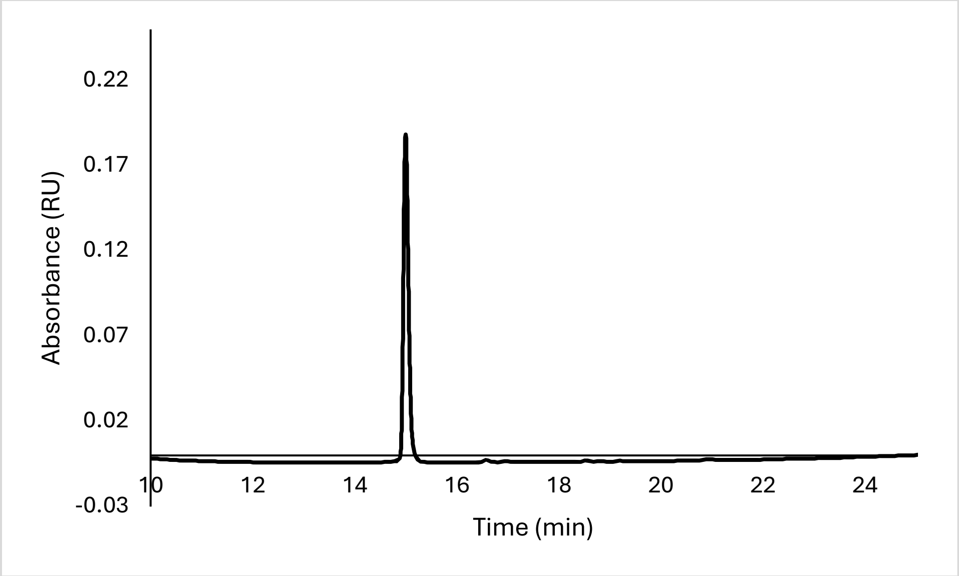


Analytical HPLC Chromatogram of **az-ovaNmet6**.

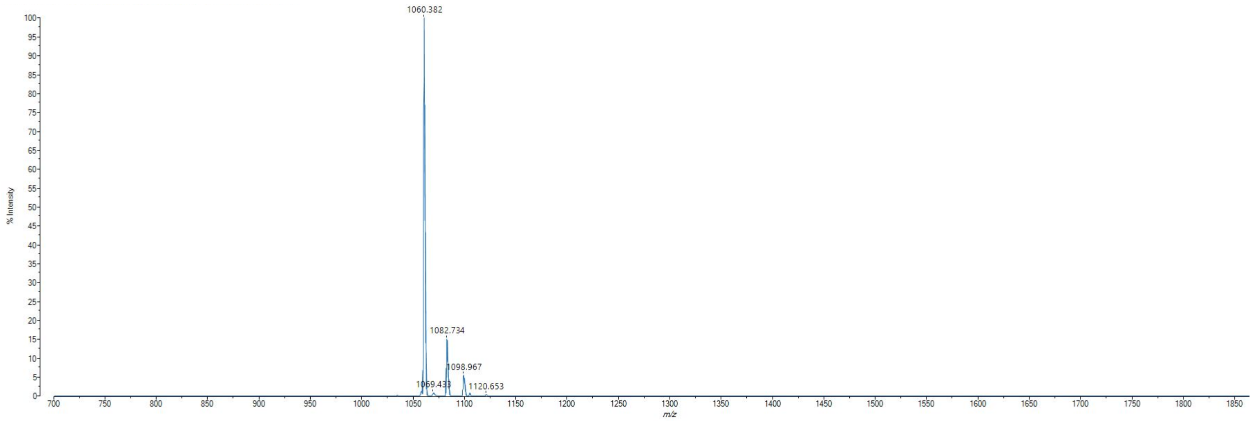


MALDI-TOF mass spectrum for **az-ovaNmet6** (m/z 1060.579 for $[M+H]^+$, found 1060.382).

**az-ovaNmet7**

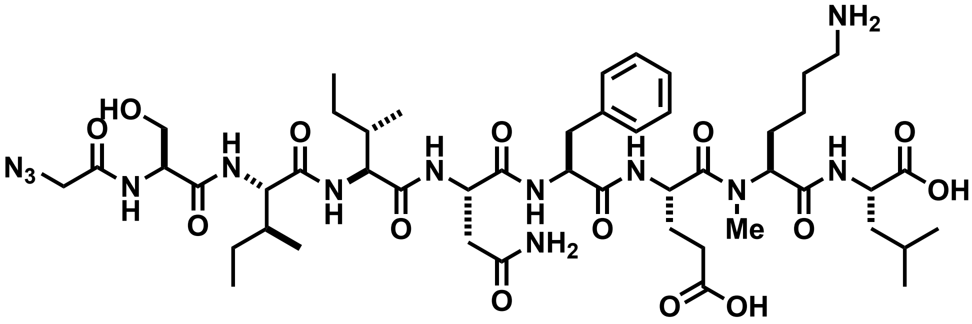

Chemical structure of **az-ovaNmet7**.

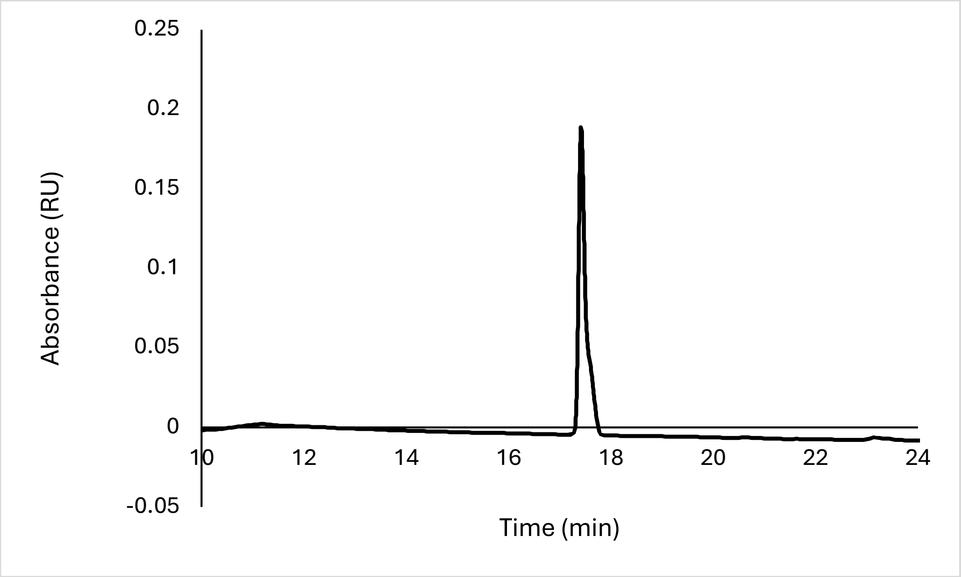


Analytical HPLC Chromatogram of **az-ovaNmet7**.

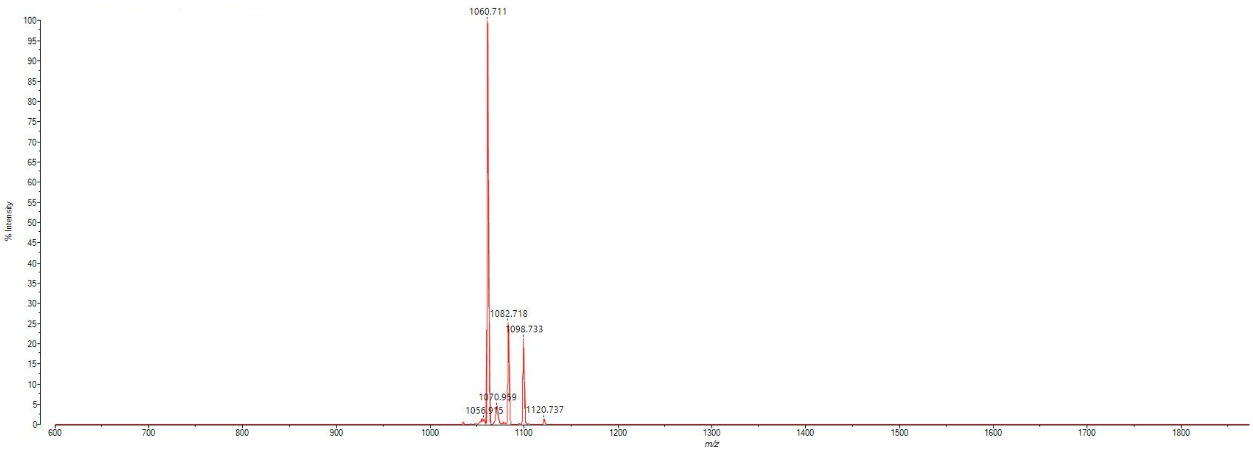


MALDI-TOF mass spectrum for **az-ovaNmet7** (m/z 1060.579 for $[M+H]^+$, found 1060.711).

**az-ovaNmet8**

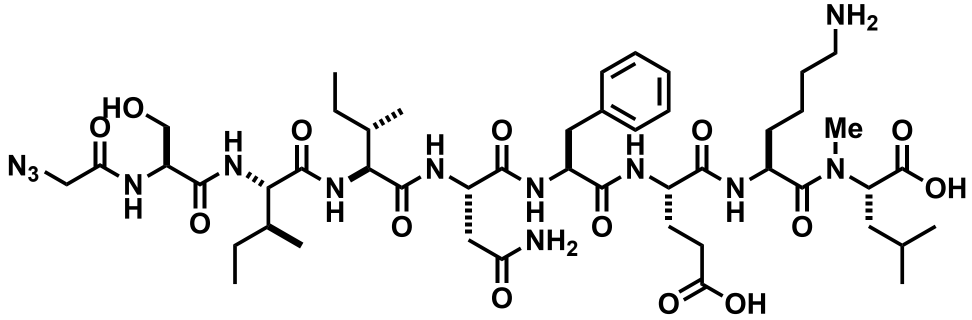


Chemical structure of **az-ovaNmet8**.

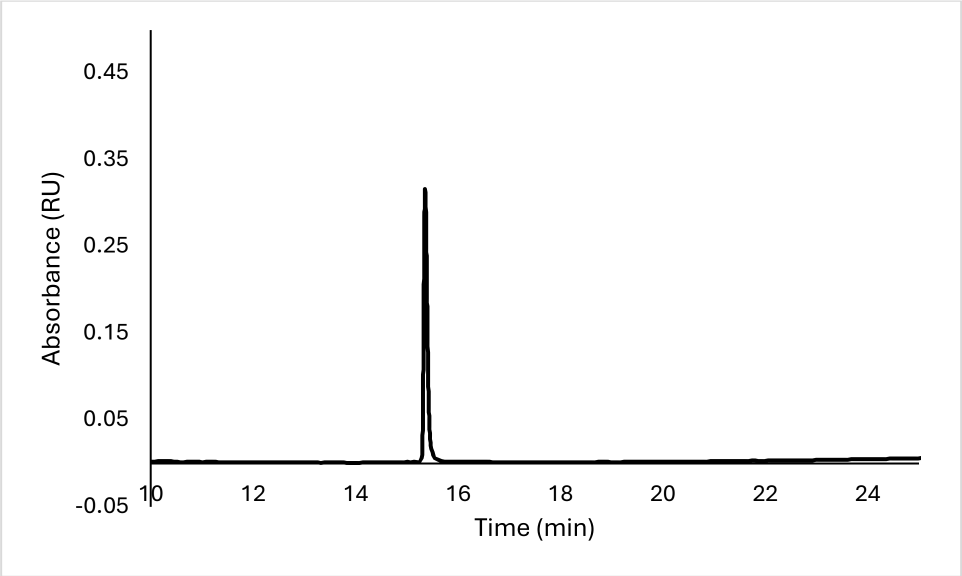


Analytical HPLC Chromatogram of **az-ovaNmet8**.

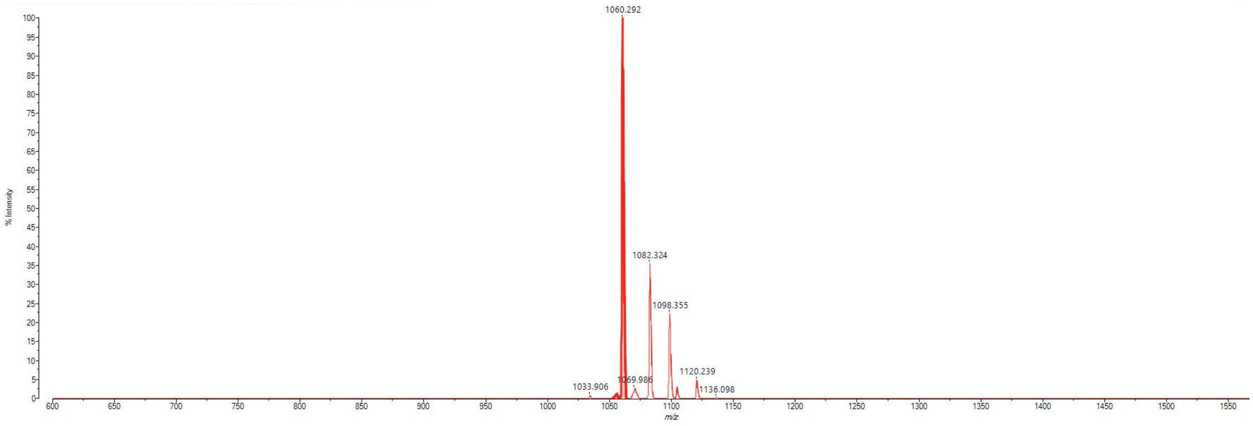


MALDI-TOF mass spectrum for **az-ovaNmet8** (m/z 1060.579 for $[M+H]^+$, found 1060.292).

**az-ovaNalk1**

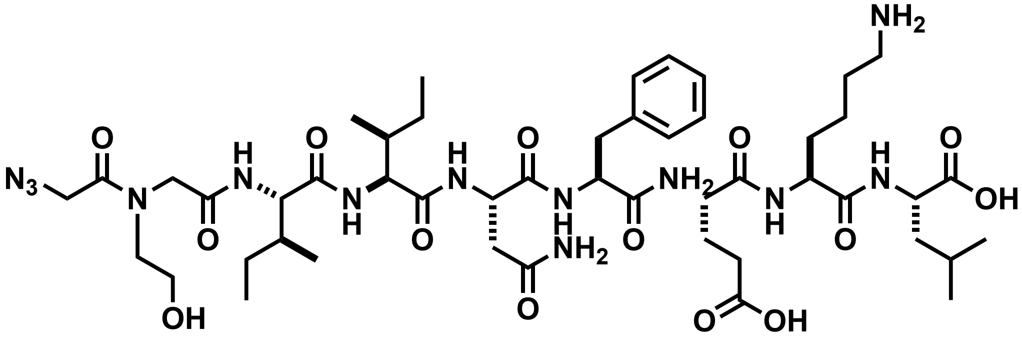

Chemical structure of **az-ovaNalk1**.

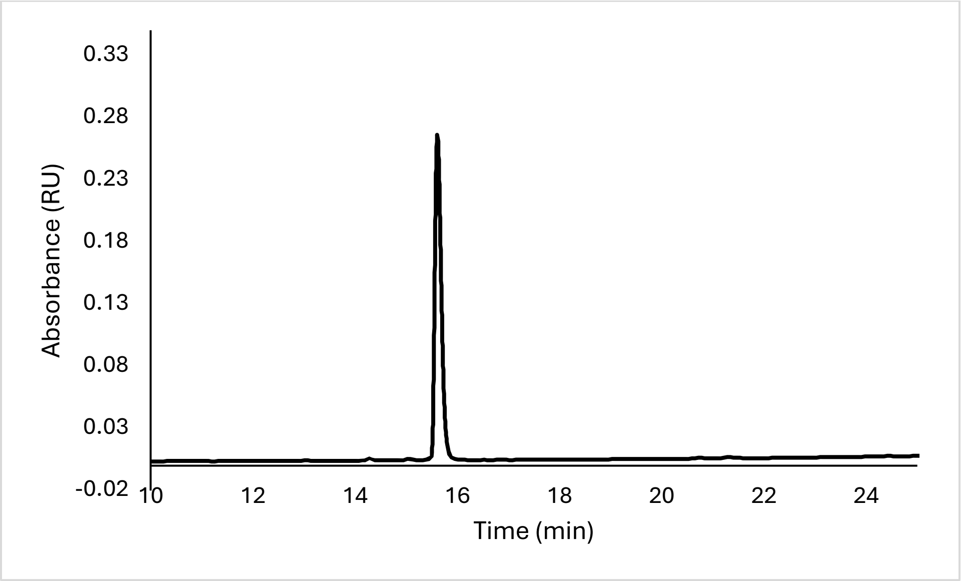


Analytical HPLC Chromatogram of **az-ovaNalk1**.

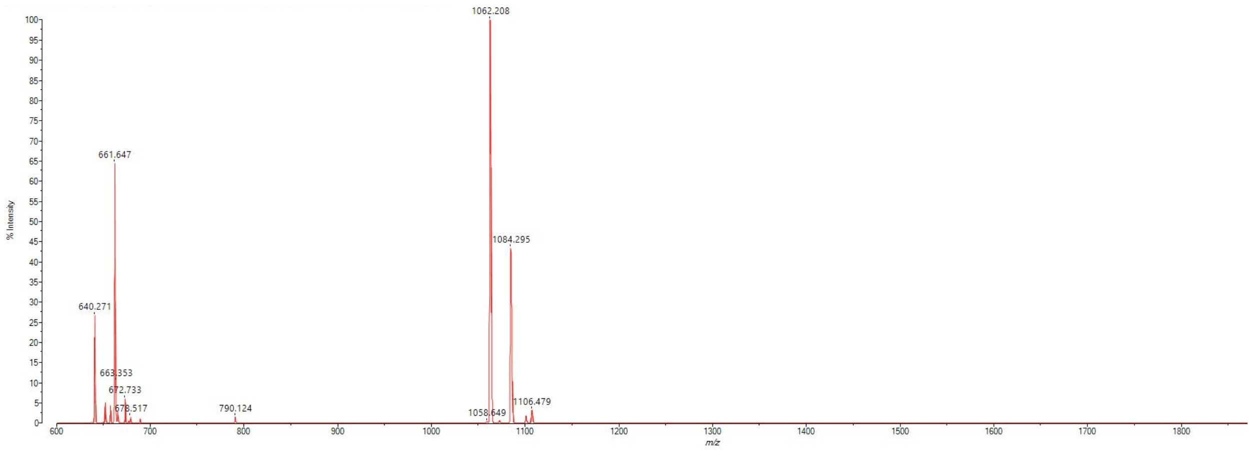


MALDI-TOF mass spectrum for **az-ovaNalk1** (m/z 1060.579 for $[M+H]^+$, found 1062.208).

**az-ovaNalk2**

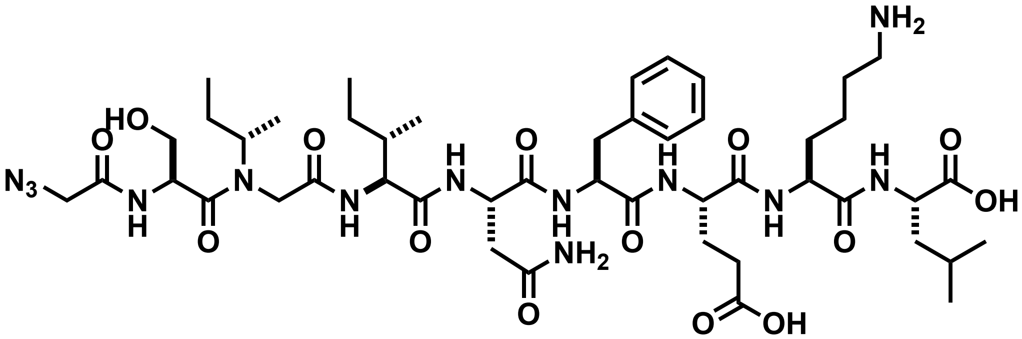

Chemical structure of **az-ovaNalk2**.

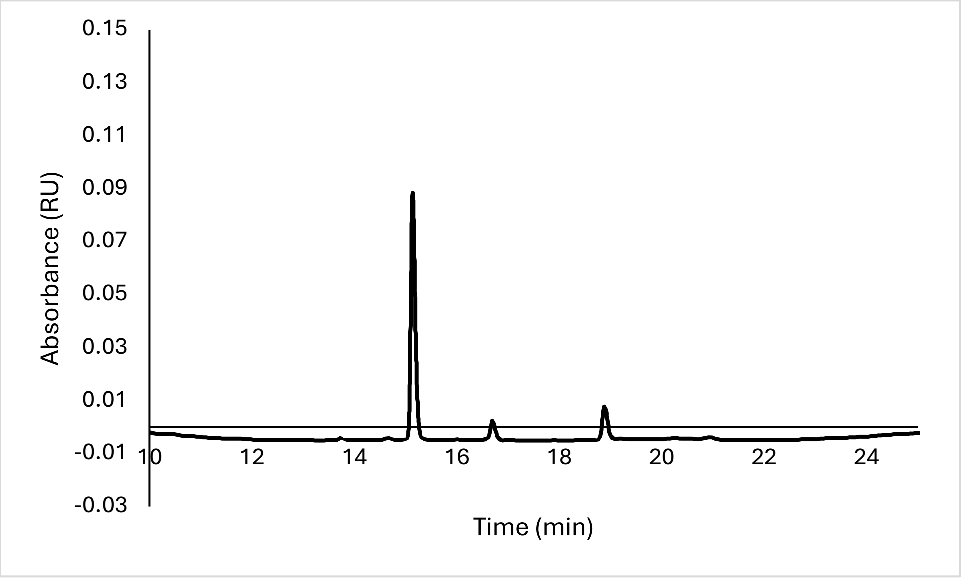


Analytical HPLC Chromatogram of **az-ovaNalk2**.

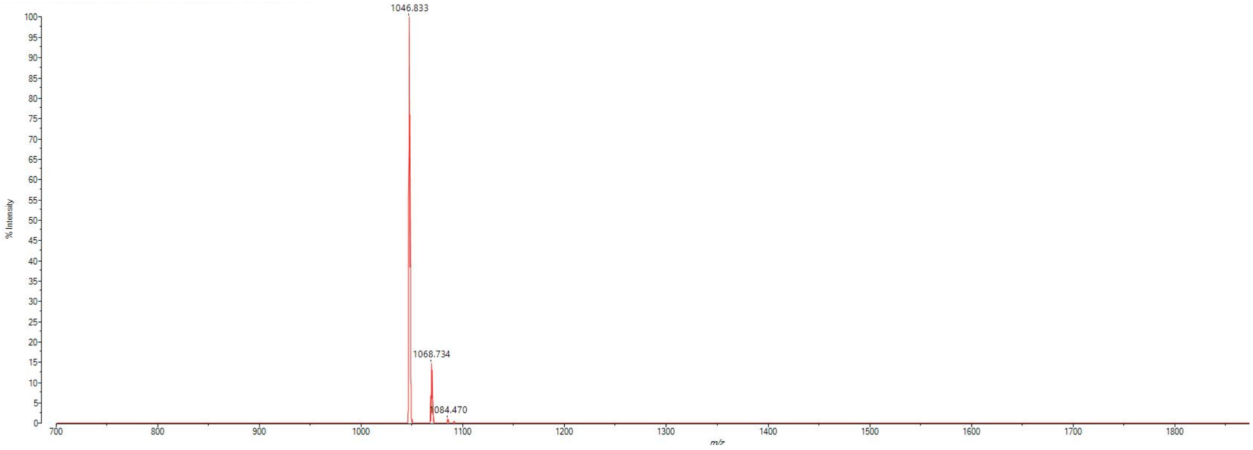


MALDI-TOF mass spectrum for **az-ovaNalk2** (m/z 1046.563 for $[M+H]^+$, found 1046.833).

**az-ovaNalk3**

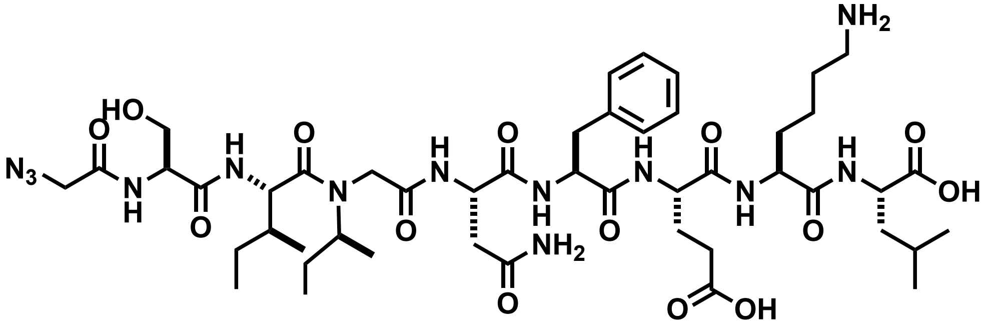


Chemical structure of **az-ovaNalk3**.

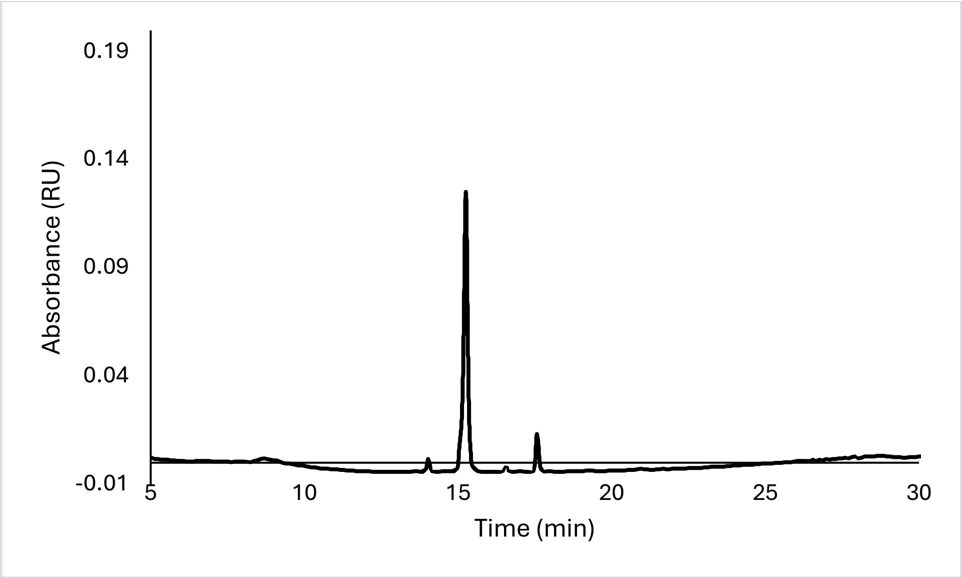


Analytical HPLC Chromatogram of **az-ovaNalk3**.

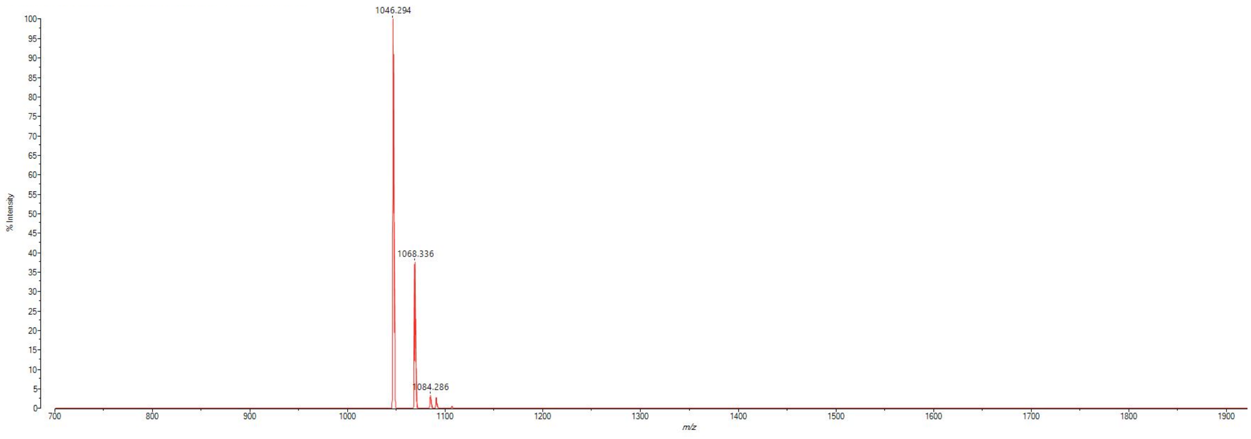


MALDI-TOF mass spectrum for **az-ovaNalk3** (m/z 1046.563 for $[M+H]^+$, found 1046.294).

**az-ovaNalk4**

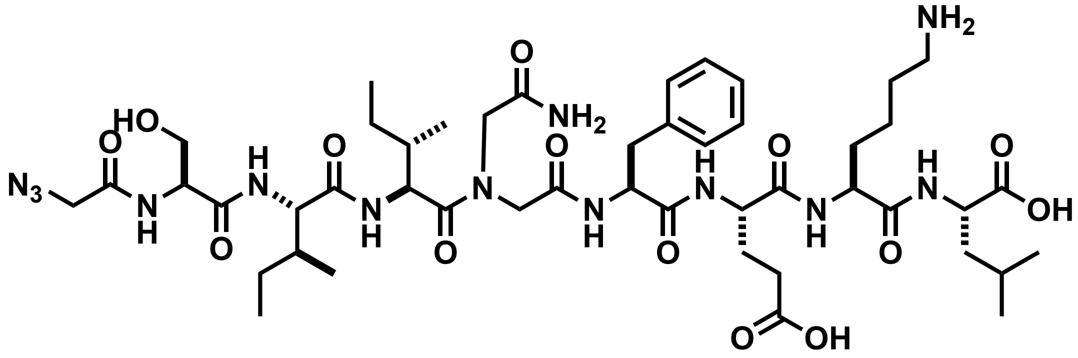

Chemical structure of **az-ovaNalk4**.

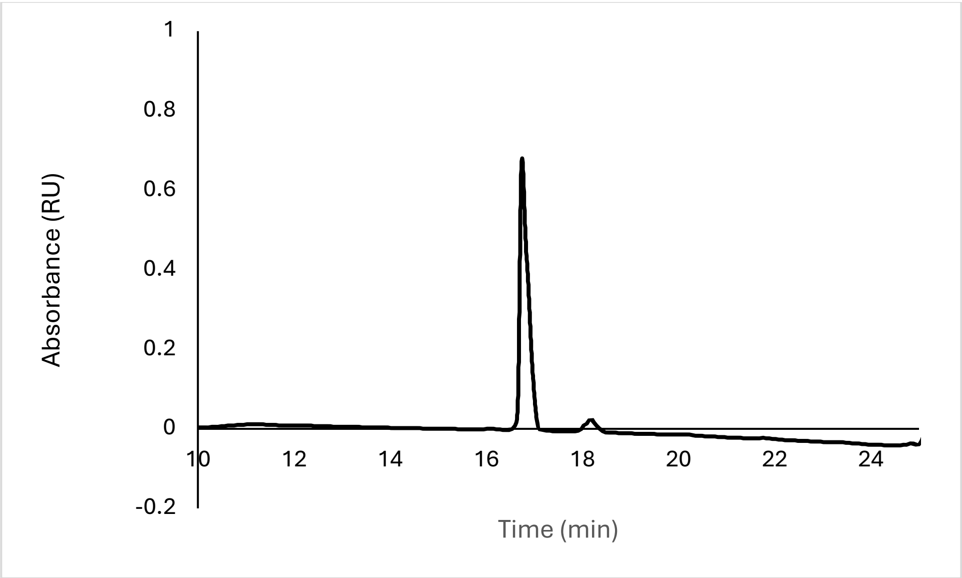


Analytical HPLC Chromatogram of **az-ovaNalk4**.

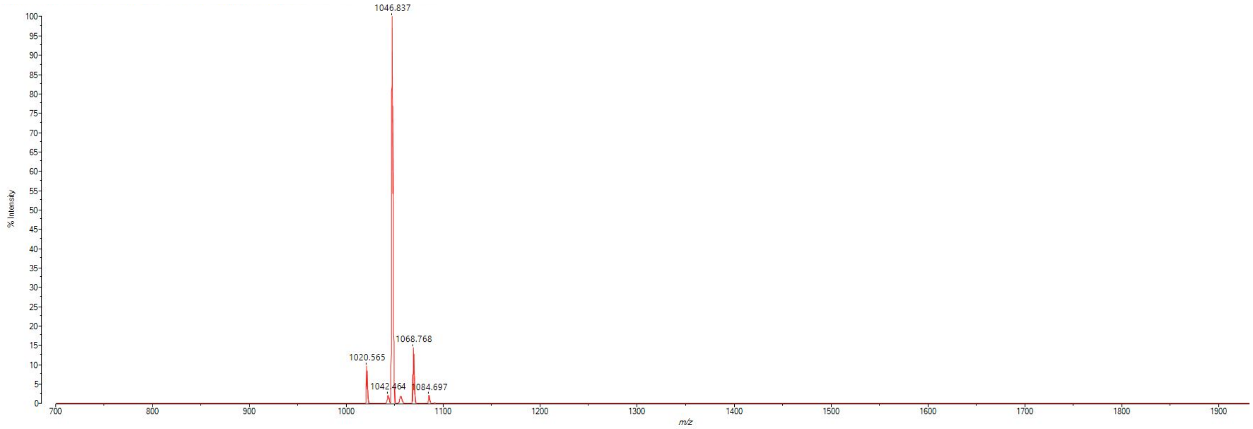


MALDI-TOF mass spectrum for **az-ovaNalk4** (m/z 1046.563 for $[M+H]^+$, found 1046.837).

**az-ovaNalk5**

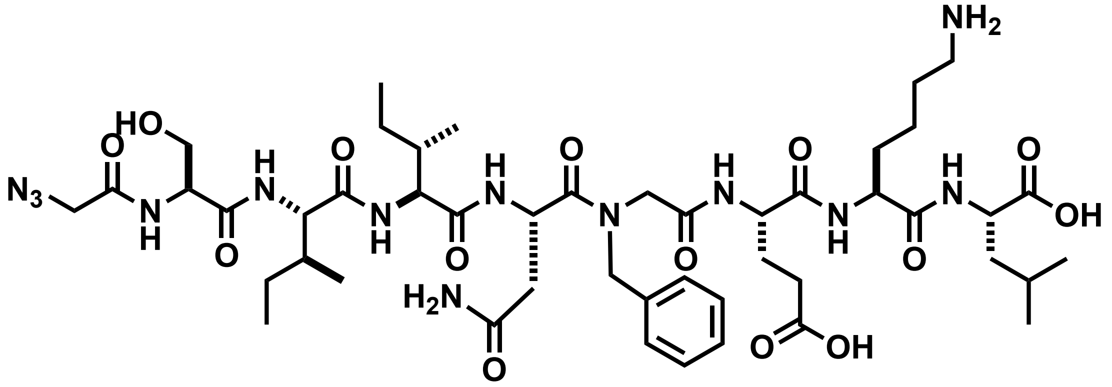


Chemical structure of **az-ovaNalk5**.

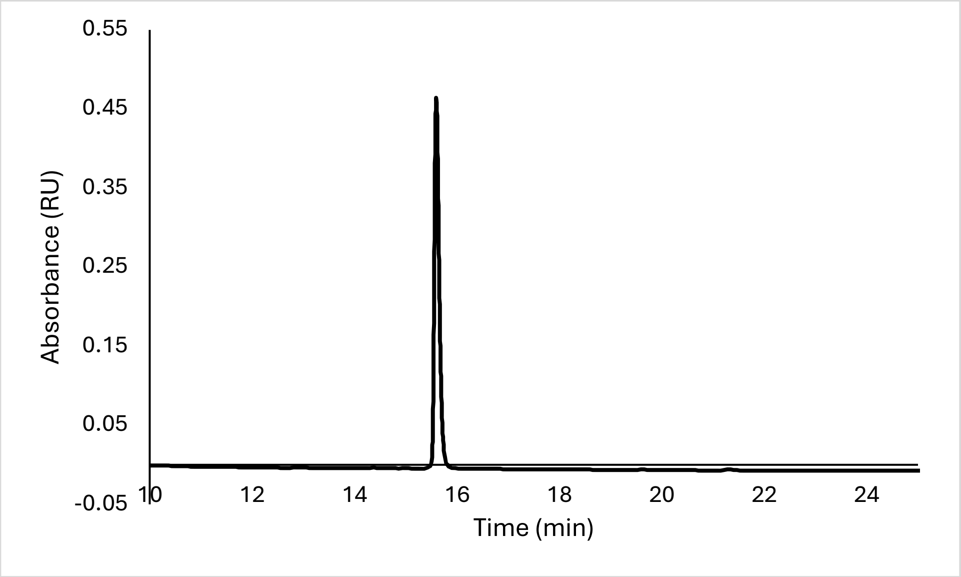


Analytical HPLC Chromatogram of **az-ovaNalk5**.

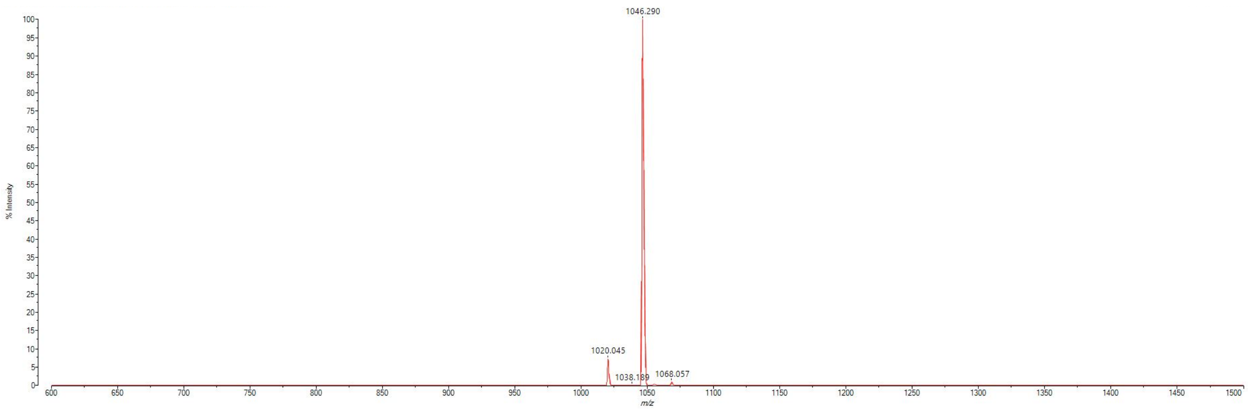


MALDI-TOF mass spectrum for **az-ovaNalk5** (m/z 1046.563 for $[M+H]^+$, found 1046.290).

**az-ovaNalk6**

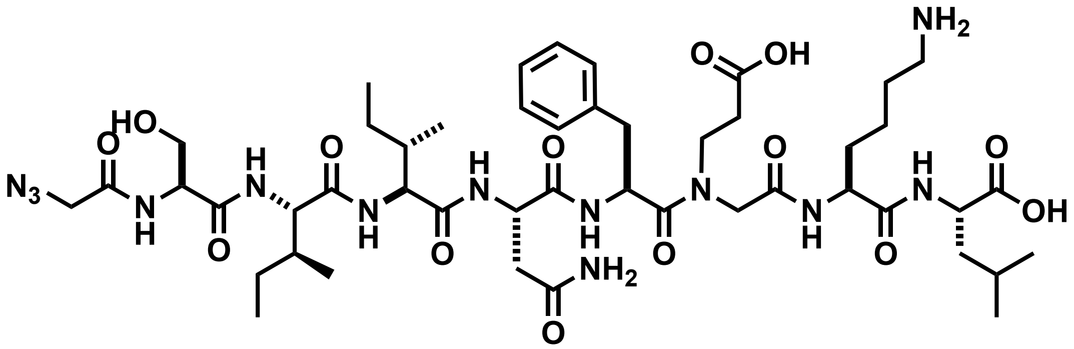


Chemical structure of **az-ovaNalk6**.

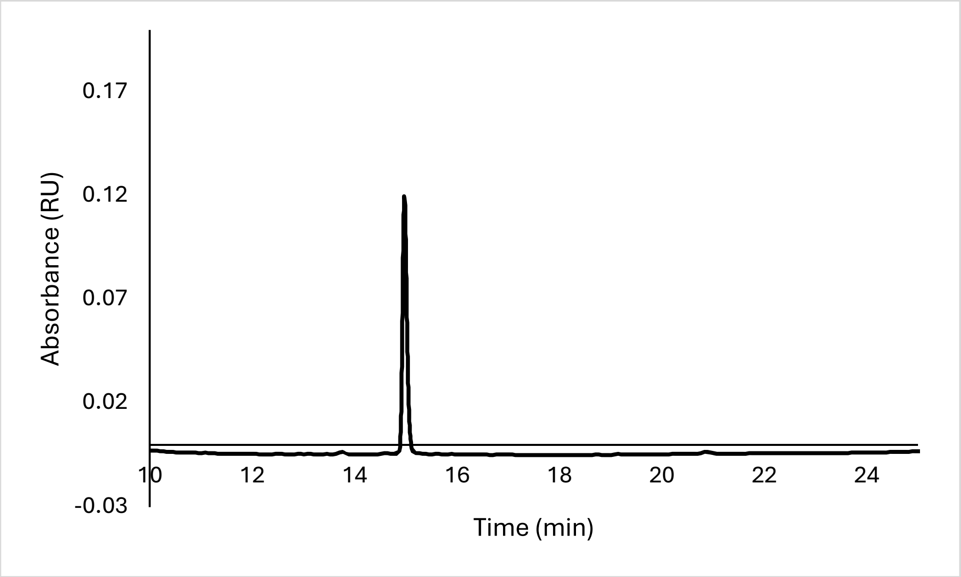


Analytical HPLC Chromatogram of **az-ovaNalk6**.

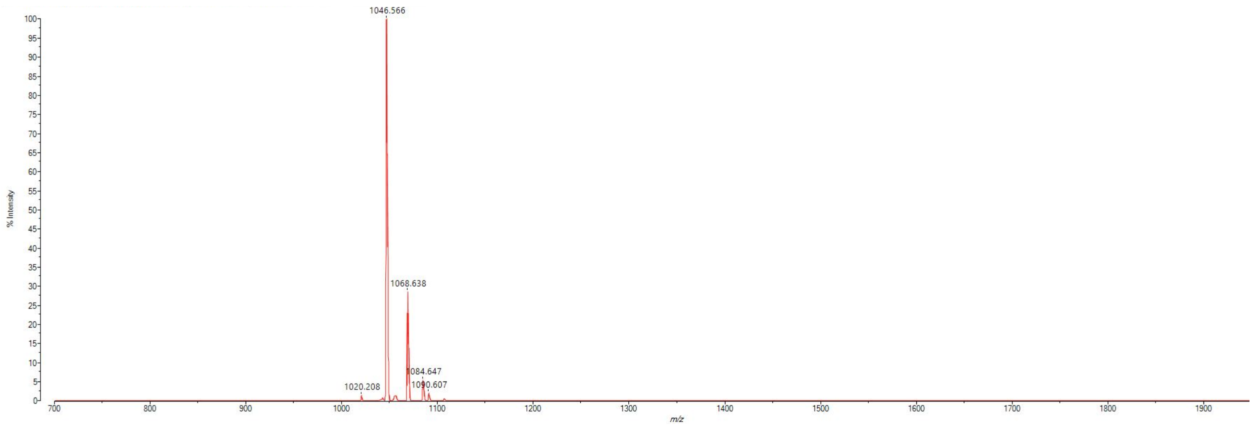


MALDI-TOF mass spectrum for **az-ovaNalk6** (m/z 1046.563 for $[M+H]^+$, found 1046.566).

**az-ovaNalk7**

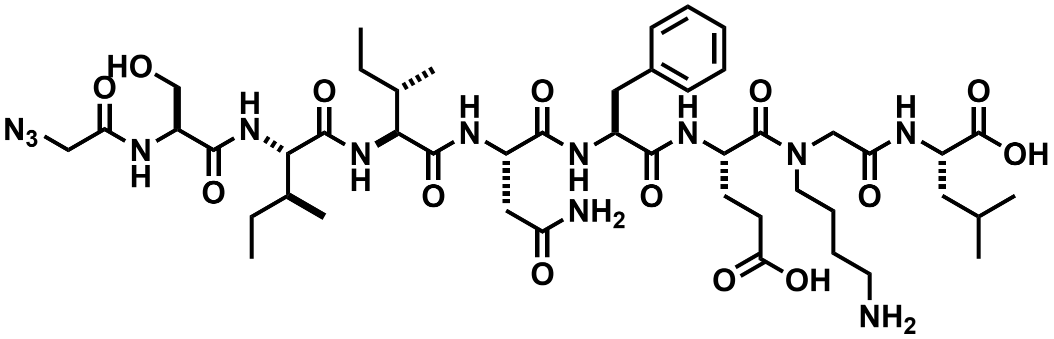


Chemical structure of **az-ovaNalk7**.

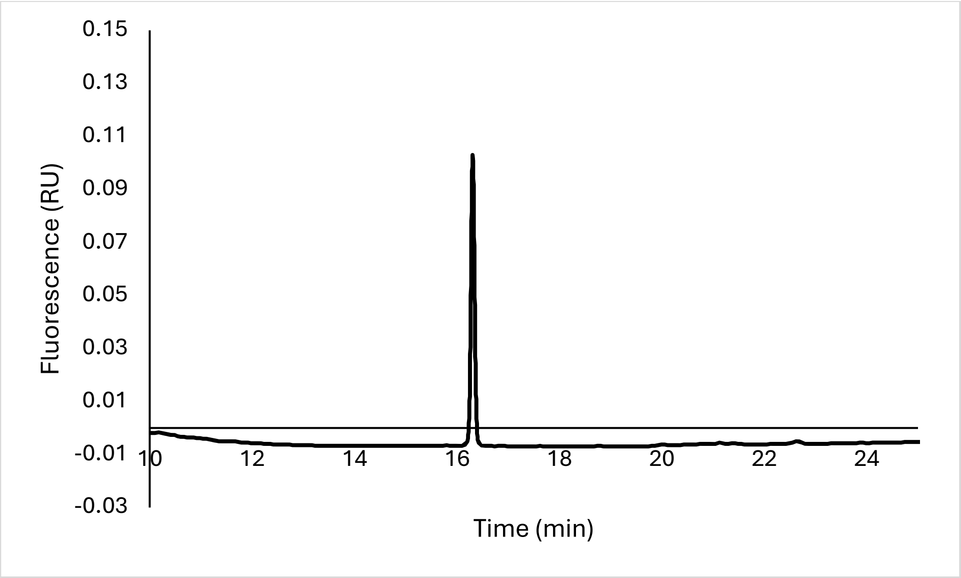


Analytical HPLC Chromatogram of **az-ovaNalk7**.

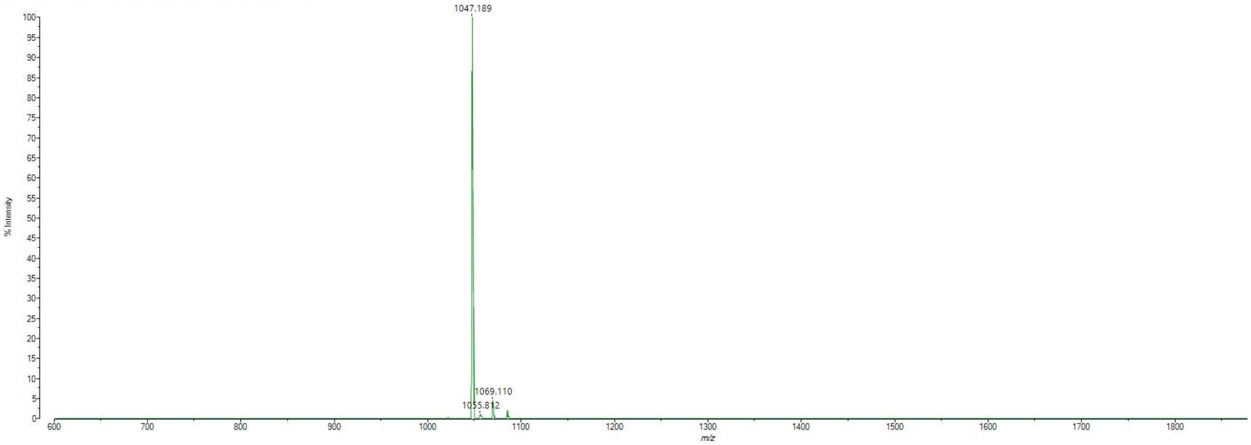


MALDI-TOF mass spectrum for **az-ovaNalk7** (m/z 1046.563 for $[M+H]^+$, found 1047.189).

**az-ovaNalk8**

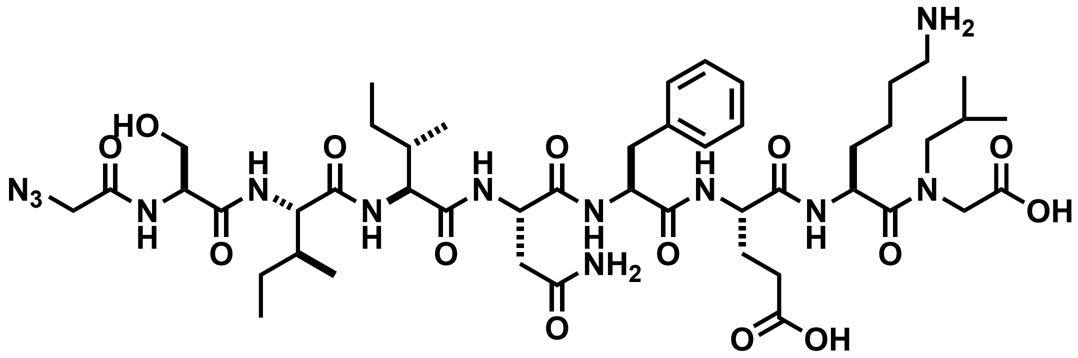


Chemical structure of **az-ovaNalk8**.

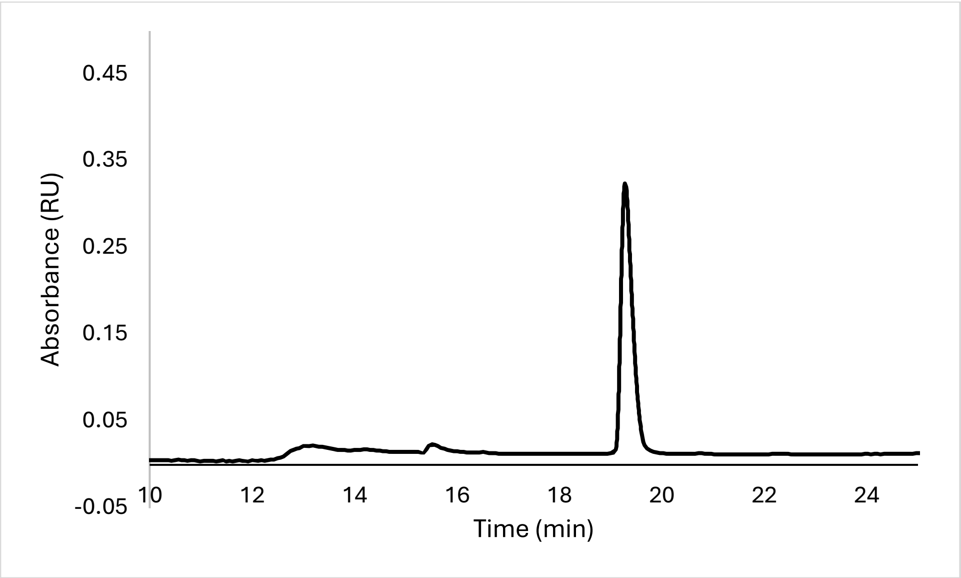


Analytical HPLC Chromatogram of **az-ovaNalk8**.

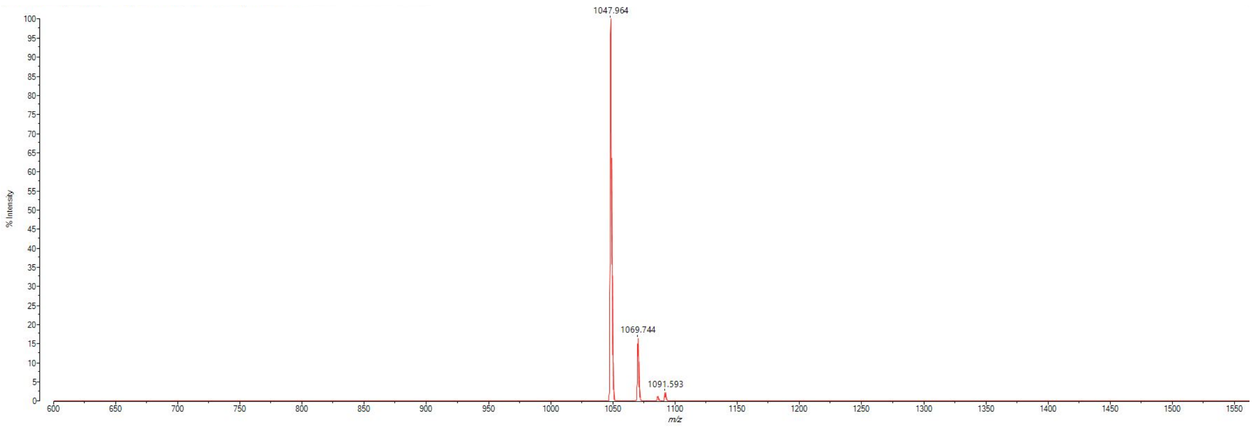


MALDI-TOF mass spectrum for **az-ovaNalk8** (m/z 1046.563 for $[M+H]^+$, found 1047.964).

**az-ovaD1**

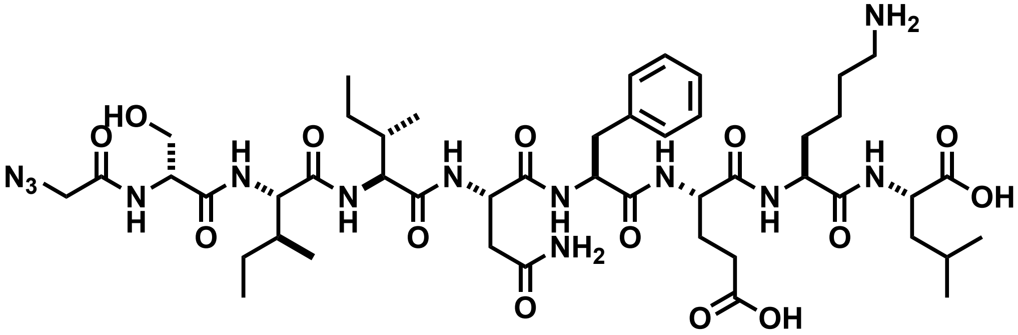


Chemical structure of **az-ovaD1**.

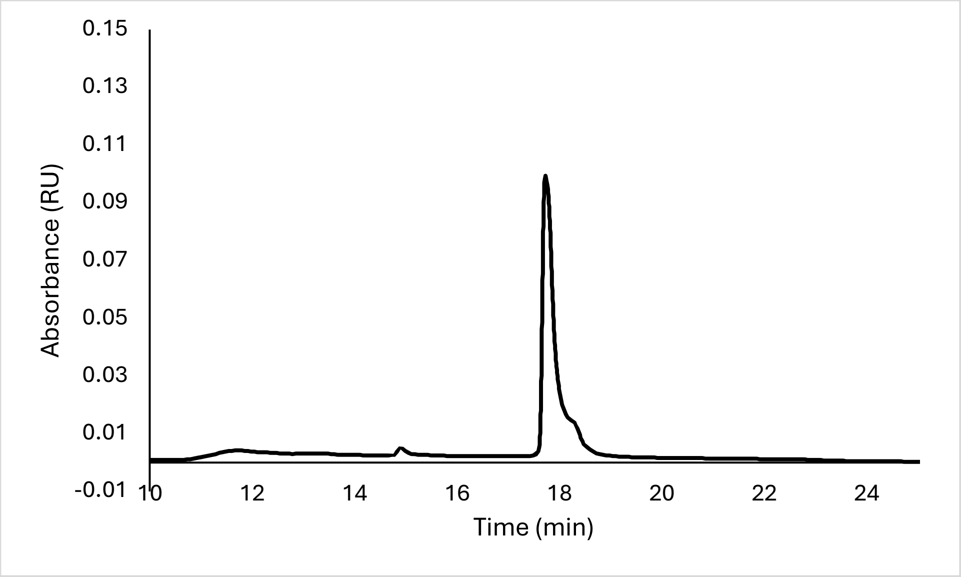


Analytical HPLC Chromatogram of **az-ovaD1**.

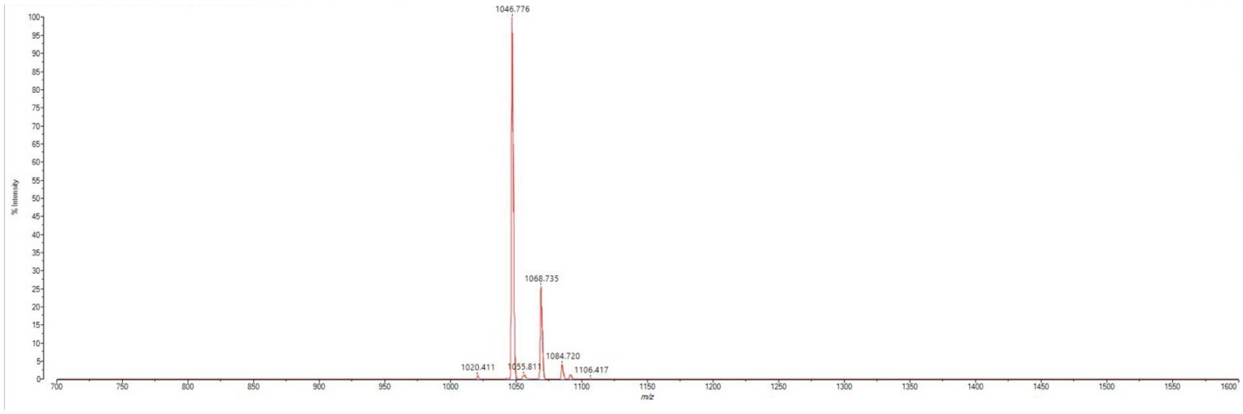


MALDI-TOF mass spectrum for **az-ovaD1** (m/z 1046.563 for $[M+H]^+$, found 1046.776).

**az-ovaD2**

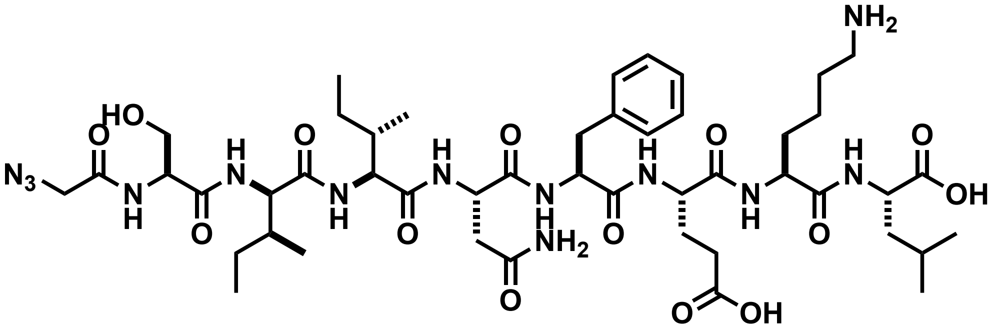

Chemical structure of **az-ovaD2**.

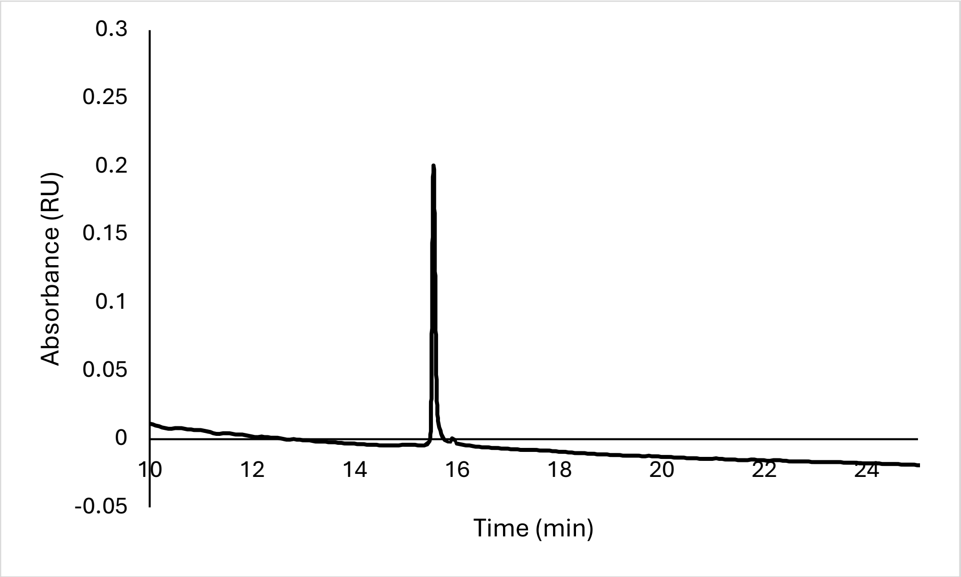


Analytical HPLC Chromatogram of **az-ovaD2**.

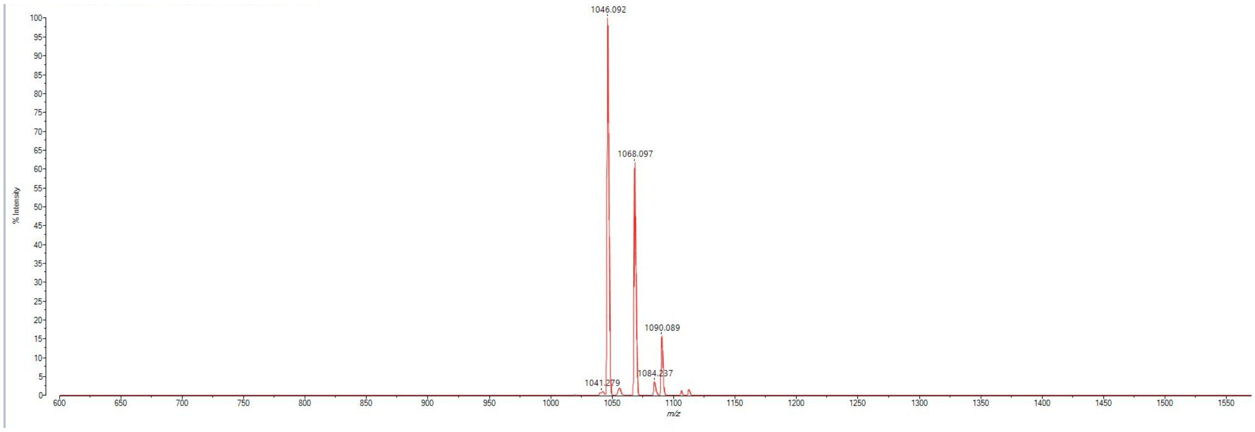


MALDI-TOF mass spectrum for **az-ovaD2** (m/z 1046.563 for $[M+H]^+$, found 1046.092).

**az-ovaD3**

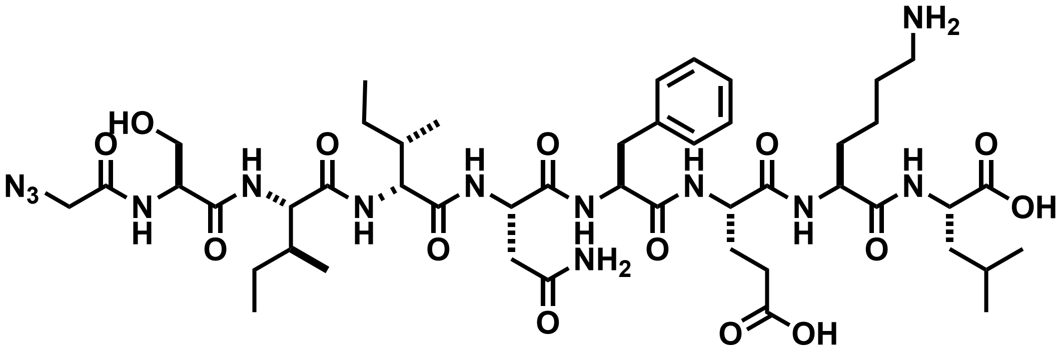

Chemical structure of **az-ovaD3**.

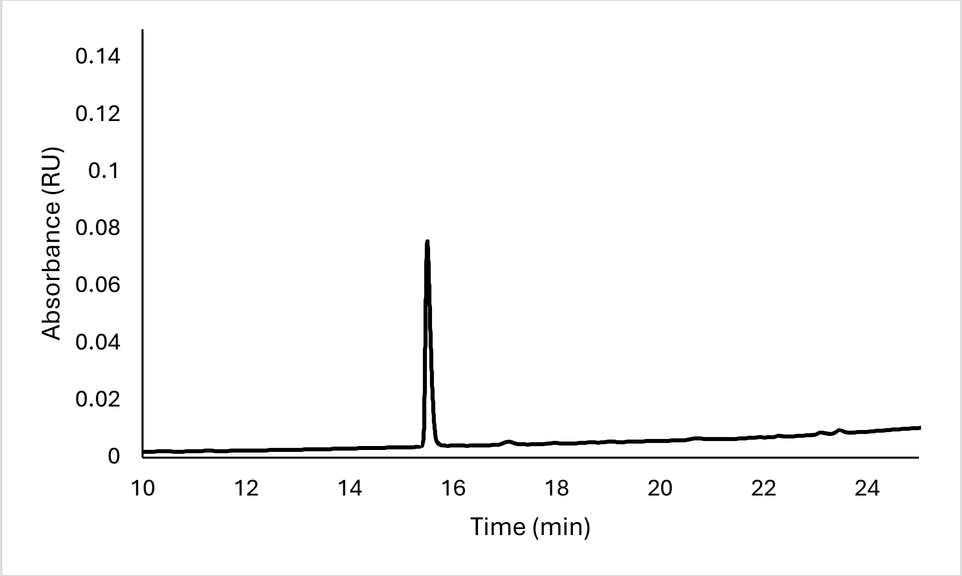


Analytical HPLC Chromatogram of **az-ovaD3**.

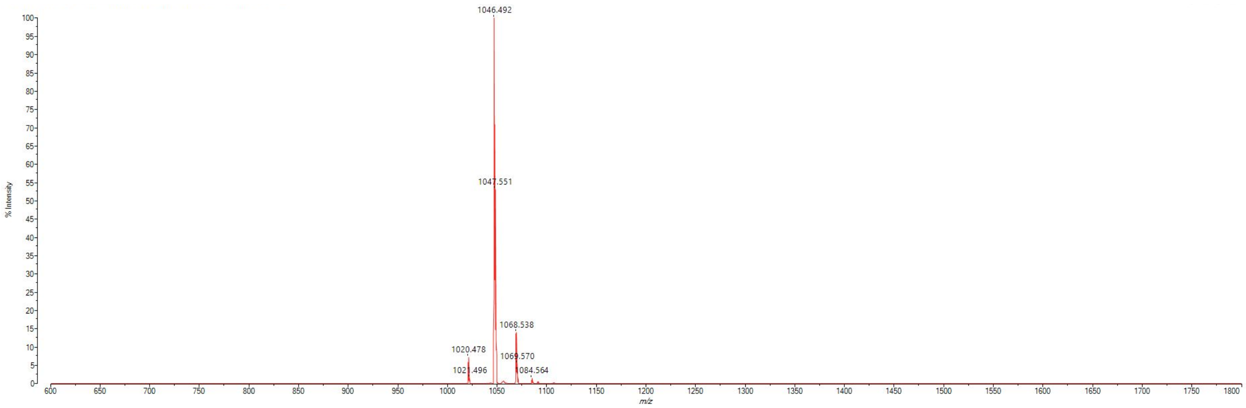


MALDI-TOF mass spectrum for **az-ovaD3** (m/z 1046.563 for $[M+H]^+$, found 1046.492).

**az-ovaD4**

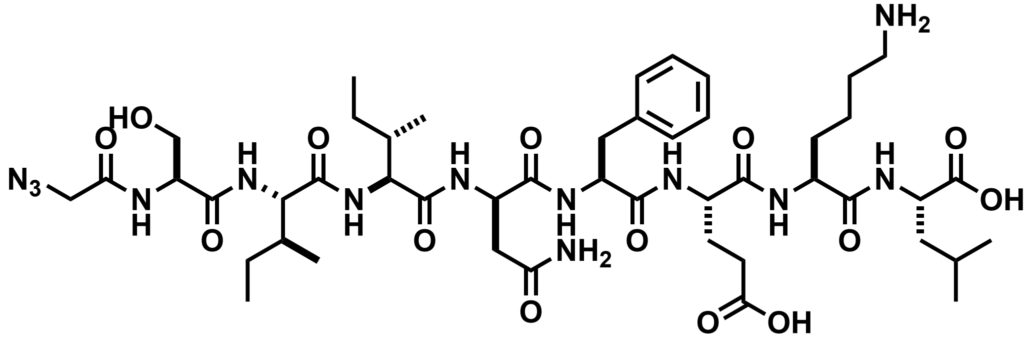


Chemical structure of **az-ovaD4**.

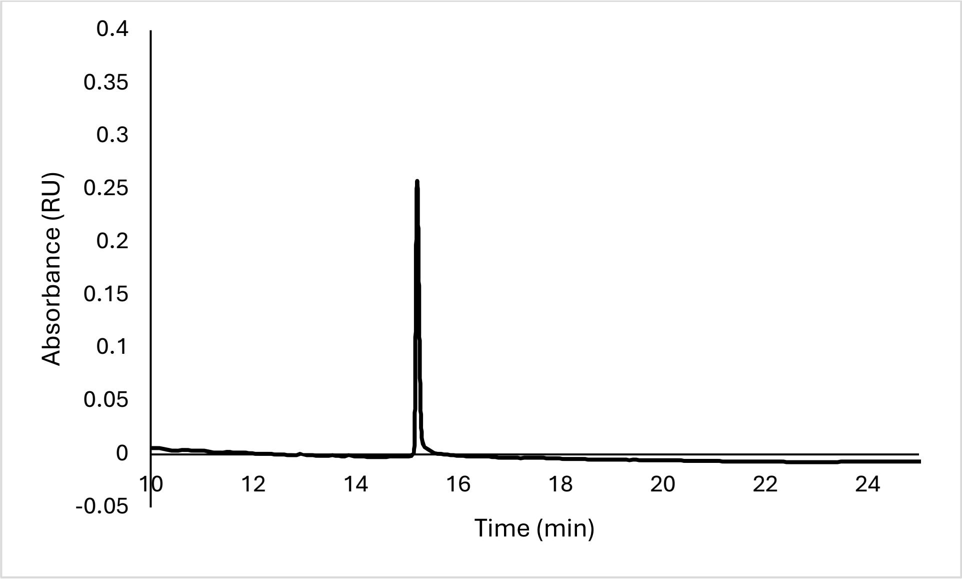


Analytical HPLC Chromatogram of **az-ovaD4**.

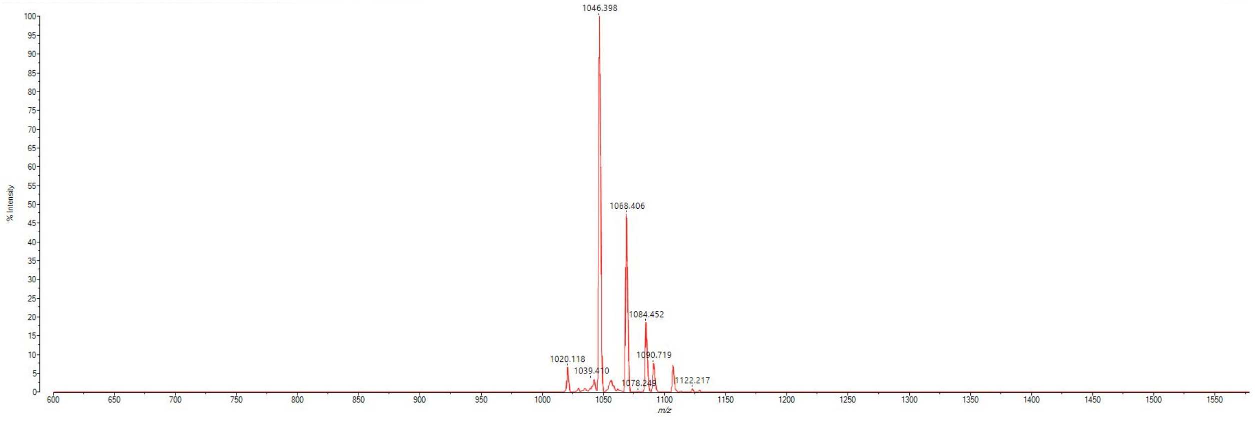


MALDI-TOF mass spectrum for **az-ovaD4** (m/z 1046.563 for $[M+H]^+$, found 1046.398).

**az-ovaD5**

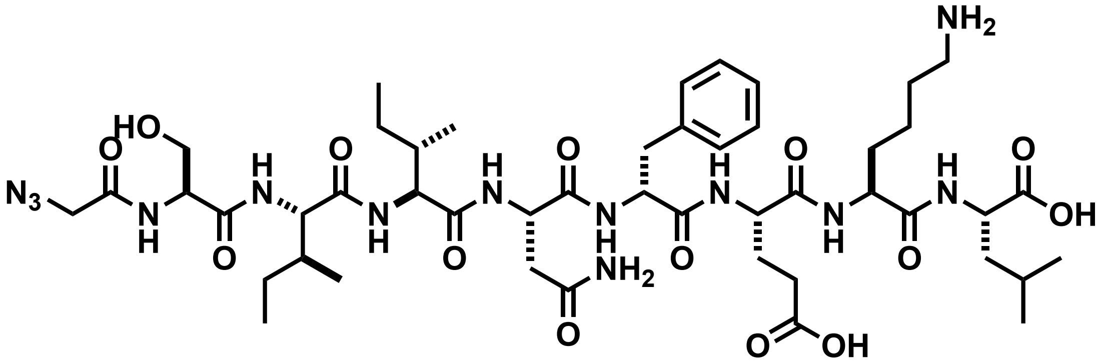

Chemical structure of **az-ovaD5**.

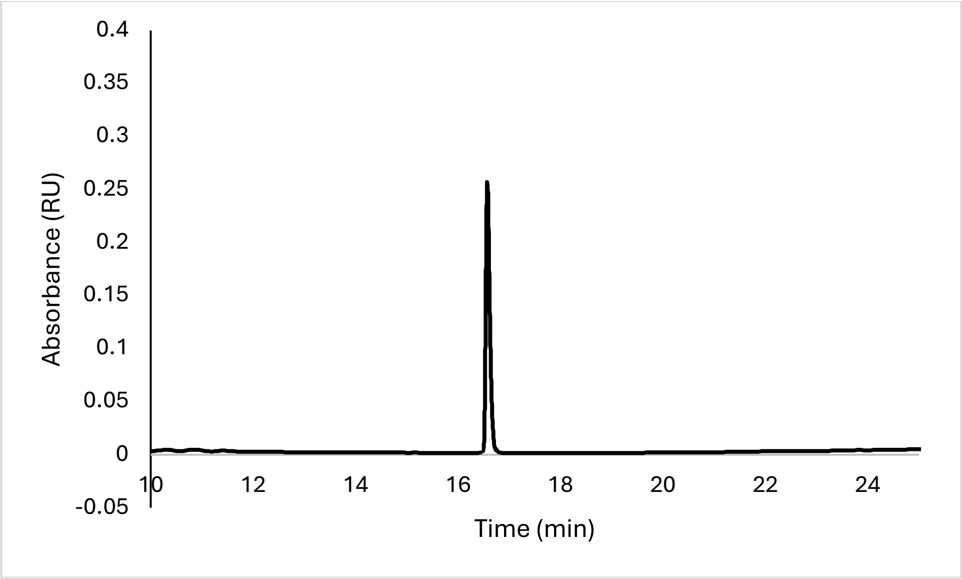


Analytical HPLC Chromatogram of **az-ovaD5**.

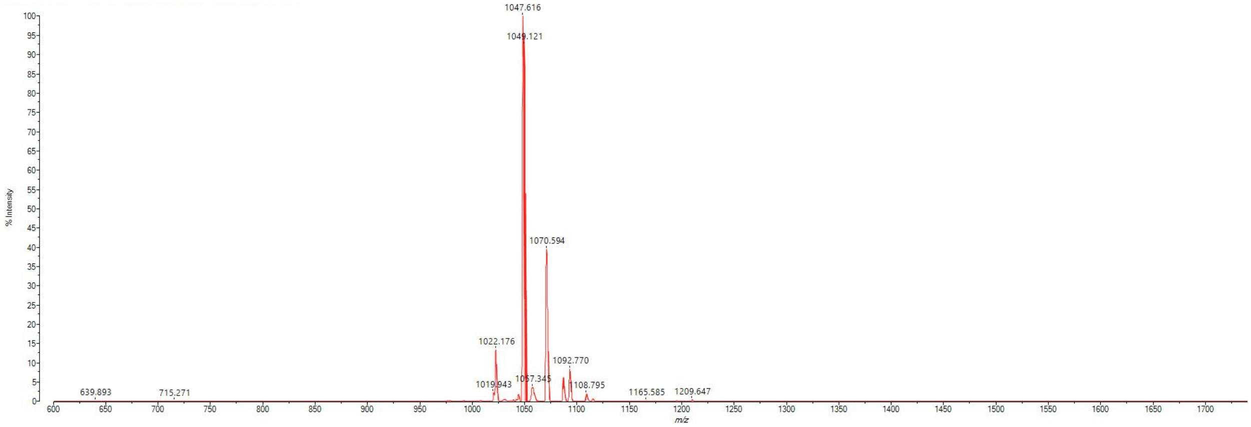


MALDI-TOF mass spectrum for **az-ovaD5** (m/z 1046.563 for $[M+H]^+$, found 1047.616).

**az-ovaD6**

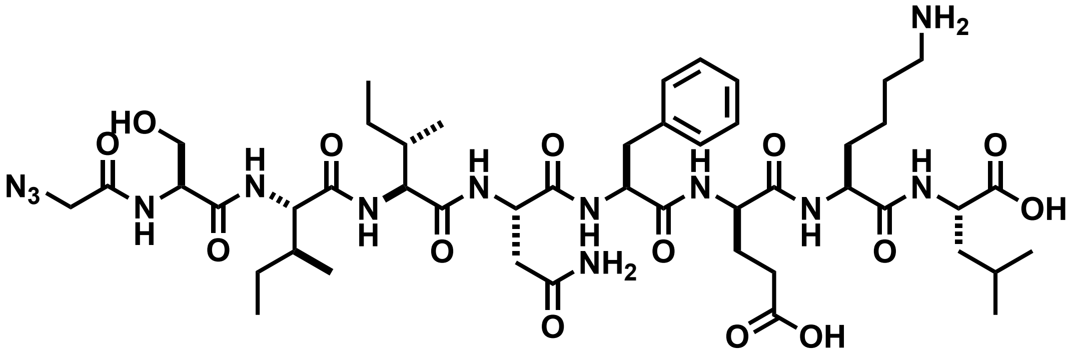


Chemical structure of **az-ovaD6**.

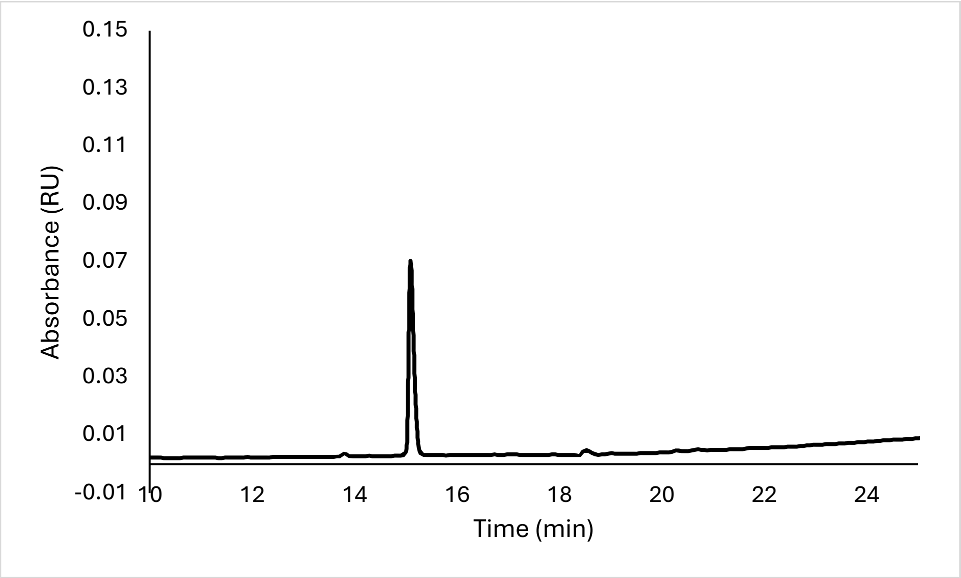


Analytical HPLC Chromatogram of **az-ovaD6**.

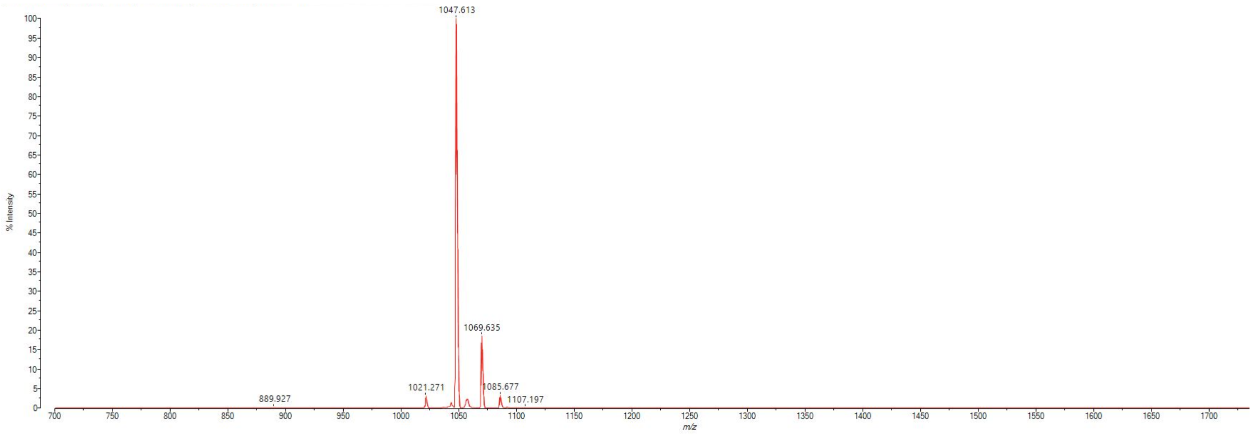


MALDI-TOF mass spectrum for **az-ovaD6** (m/z 1046.563 for $[M+H]^+$, found 1047.613).

**az-ovaD7**

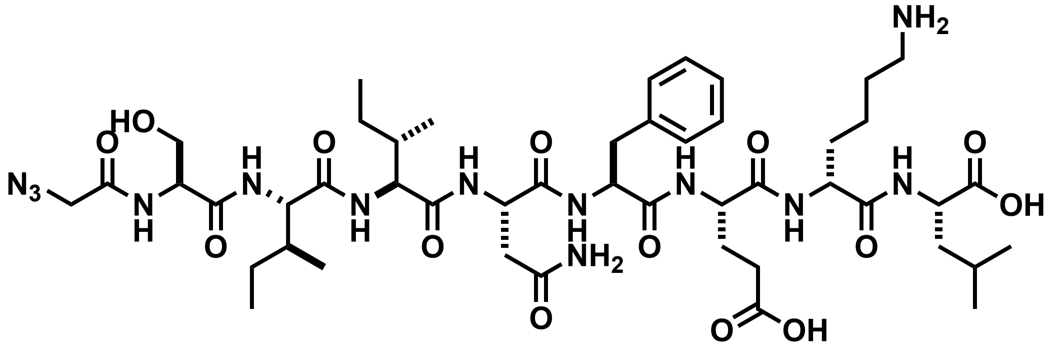

Chemical structure of **az-ovaD7**.

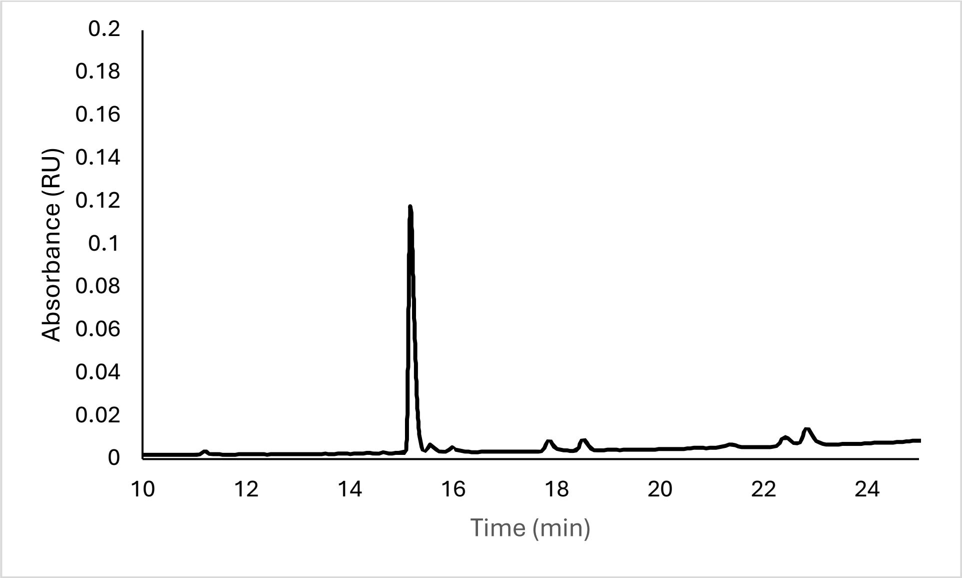


Analytical HPLC Chromatogram of **az-ovaD7**.

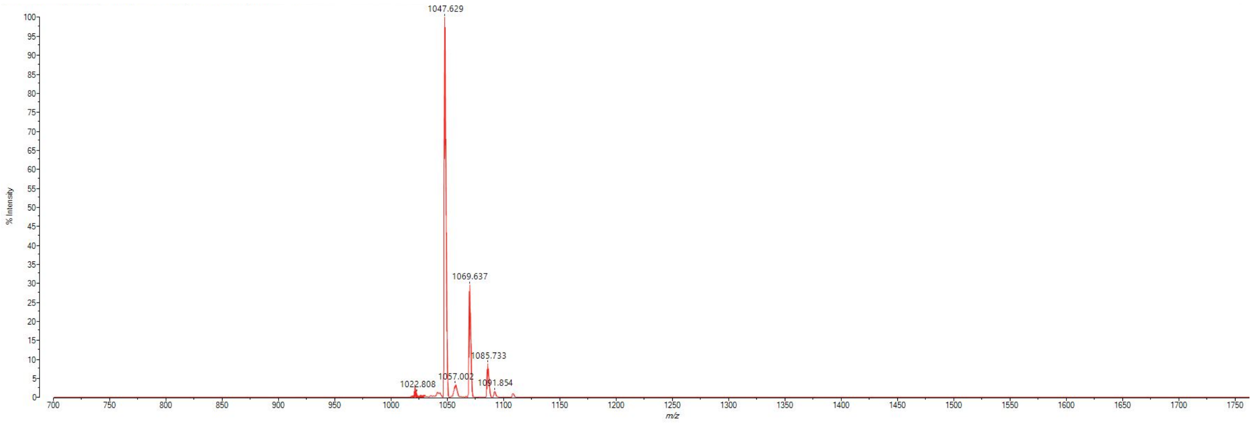


MALDI-TOF mass spectrum for **az-ovaD7** (m/z 1046.563 for $[M+H]^+$, found 1047.629).

**az-ovaD8**

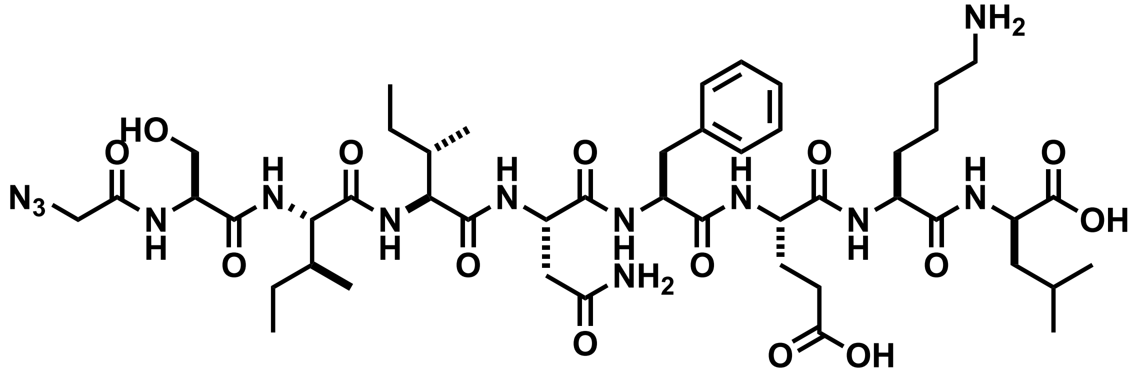

Chemical structure of **az-ovaD8**.

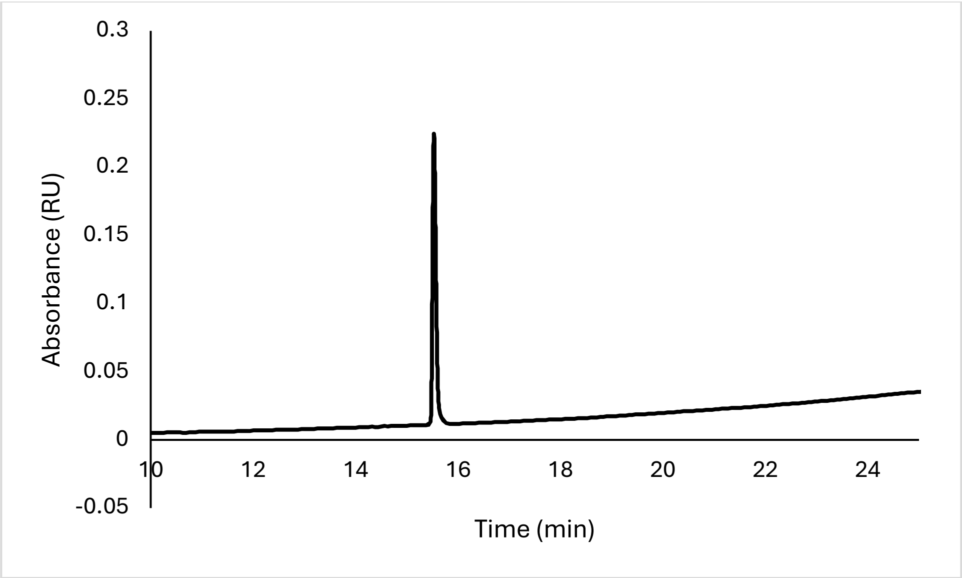


Analytical HPLC Chromatogram of **az-ovaD8**.

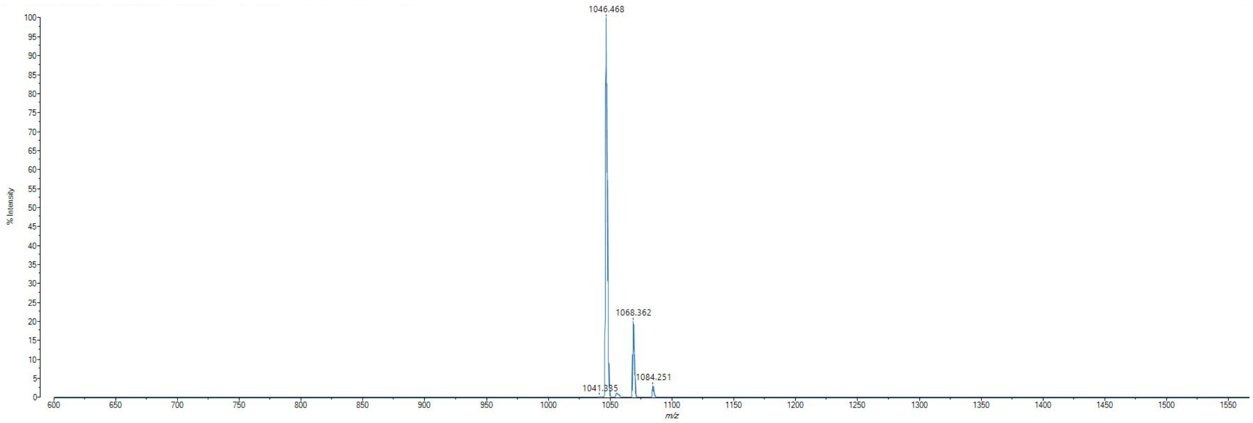


MALDI-TOF mass spectrum for **az-ovaD8** (m/z 1046.563 for $[M+H]^+$, found 1046.468).
